# An accumbal cholinergic ISR-HCN2 axis sustains maladaptive cue motivation

**DOI:** 10.64898/2026.09.18.752826

**Authors:** Shihao Huang, Jingliang Zhang, Yue Li, Xinyou Lv, Zhihao Song, Zengbo Ding, Zhonghao Li, Yu Tian, Xiangyang Zang, Yuchuan Hu, Yixiao Luo, Haoyu Li, Jie Shi, Lin Lu, Zhuo Huang, Yan-Xue Xue

## Abstract

Environmental cues associated with rewarding outcomes can exert persistent control over behavior long after the reward is absent, yet the cellular adaptations that stabilize such maladaptive motivation remain unclear. Here we identify nucleus accumbens cholinergic interneurons (CINs) as causal regulators of persistent cue-driven seeking. Self-administration of addictive drugs or high-fat reward, but not sucrose, selectively increased CIN pacemaker firing. Cocaine-paired cues evoked rapid acetylcholine release in the nucleus accumbens core, and bidirectional manipulation of CINs suppressed or enhanced cue-induced cocaine seeking. Mechanistically, cocaine experience engaged the integrated stress response (ISR) in CINs, increased HCN2 expression and *I*_h_, and stabilized a hyperexcitable state. CIN-specific ISR suppression or HCN2 knockdown reduced cocaine seeking, whereas HCN2 overexpression enhanced it. Local HCN blockade and systemic treatment with the clinically approved HCN blocker ivabradine reduced cue-driven seeking across addictive-drug and high-fat-reward paradigms while sparing sucrose seeking. These findings identify an ISR-HCN2-dependent cholinergic mechanism that stabilizes maladaptive cue-driven motivation and nominate HCN-dependent excitability as a tractable target for therapeutic intervention.

## Introduction

Environmental cues guide adaptive behavior by allowing organisms to predict, prioritize, and pursue biologically valuable outcomes^1,2^. In maladaptive motivational states, however, this normally adaptive process becomes pathologically persistent. Cues associated with addictive drugs, and in some contexts highly palatable rewards, can acquire excessive motivational power and trigger seeking long after the primary reward is absent^3–8^. This persistent cue-driven motivation is a central feature of relapse and compulsive reward seeking. Although dopamine signaling is critical for reward prediction, reinforcement learning, and incentive motivation, phasic dopamine transients alone do not fully explain how cue-triggered motivation remains stable, selective, and behaviorally potent over prolonged periods^4,9,10^. Identifying the local circuit and molecular mechanisms that sustain maladaptive cue-driven motivation therefore remains a central challenge.

The nucleus accumbens (NAc) is a key interface between reward prediction, motivational state, and action selection^11,12^. Within this structure, cholinergic interneurons (CINs) are uniquely positioned to regulate how predictive cues influence behavior^11,12^. CINs fire autonomously, provide widespread acetylcholine release throughout local accumbal microcircuits^13–16^, and regulate both dopaminergic input^14,17,18^ and medium spiny neuron output^19–21^. Through these properties, CINs can influence circuit function on longer timescales than transient neuromodulatory signals, acting as state-setting elements that gate how predictive cues are translated into action. Consistent with this role, striatal CINs contribute to learning-related adjustments of striatal output and to the mapping of cue information onto action selection^21–24^. Their firing is transiently modulated by informative environmental stimuli, increasing to cues that discourage motivated behavior^25,26^, and exhibiting the canonical pause response to reward-predictive cues^27–29^. Recent work further shows that CIN-derived acetylcholine can regulate dopaminergic signaling in a task- and state-dependent manner, supporting high-effort reward-directed actions^30^. Together, these studies suggest that CINs normally help constrain, filter, or amplify cue impact according to behavioral context.

How this cholinergic control system is reconfigured by addictive experience remains unknown. A key unresolved question is whether repeated drug taking converts CINs from adaptive regulators of cue-guided behavior into drivers of persistent maladaptive motivation. Such a shift would require not only transient cue responses, but a durable change in intrinsic excitability capable of biasing accumbal circuits toward cue reactivity over time. One candidate mechanism is the integrated stress response (ISR), a conserved translational control pathway that regulates protein synthesis in response to cellular stress and has emerged as a modulator of synaptic plasticity, memory, and neuronal excitability^31,32^. In striatal circuits, ISR signaling operates constitutively in cholinergic interneurons and influences firing properties and learning-related adaptation^33,34^. In parallel, ISR-dependent translational control has been implicated in cocaine-induced synaptic and behavioral plasticity^35,36^. These observations raise the possibility that addictive experience engages ISR signaling in NAc CINs to install a persistent hyperexcitable state. However, whether ISR activation in CINs distinguishes maladaptive cue motivation from ordinary natural reward seeking, and which downstream effectors convert ISR signaling into lasting changes in CIN firing, remain unknown.

Here, we tested the hypothesis that accumbal CINs sustain maladaptive cue-driven motivation. We show that self-administration of addictive drugs and high-fat reward, but not yoked cocaine exposure or sucrose self-administration, persistently increases CIN pacemaker firing. Cocaine-paired cues evoke rapid acetylcholine release in the NAc core, and bidirectional manipulation of CINs respectively suppresses or enhances cue-induced cocaine seeking. Mechanistically, cocaine experience engages ISR signaling in CINs, upregulates HCN2 channels, enhances *I*_h_, and stabilizes a hyperexcitable CIN state. Genetic suppression of CIN ISR signaling or CIN-specific HCN2 knockdown reduces cue-induced cocaine seeking, whereas HCN2 overexpression enhances it. Finally, local HCN blockade and systemic treatment with the FDA-approved HCN blocker ivabradine reduce cue-induced seeking across multiple addictive drugs and high-fat reward while sparing sucrose seeking. These findings define an ISR-HCN2 axis in accumbal CINs that sustains maladaptive cue-driven motivation and provides a tractable entry point for relapse prevention.

## Results

### Addictive drug and high-fat self-administration induce persistent CIN hyperexcitability in the NAc

To determine whether maladaptive cue-driven motivation is associated with altered intrinsic excitability of NAc CINs, we first performed cell-attached recordings from NAc CINs after cocaine self-administration (SA). Rats underwent 10 days of cocaine or saline SA, during which active nose pokes delivered intravenous cocaine paired with discrete cues or saline infusions, followed by 5 days of home-cage withdrawal before *ex vivo* recordings (**Fig. 1a**). In slices from these rats, large, aspiny CINs, but not neighboring MSNs, exhibited autonomous pacemaker firing (**Fig. 1b,c**). Compared with saline SA rats, CINs from cocaine SA rats displayed markedly elevated firing rates (**Fig. 1d,e**), indicating that a history of contingent cocaine intake enhances the intrinsic activity of NAc CINs. In contrast, CINs from rats that received yoked cocaine infusions, matched in total cocaine exposure but without instrumental learning, did not differ from saline SA controls, suggesting that CIN hyperexcitability depends on the associative components of drug taking rather than passive pharmacological exposure alone.

**Fig. 1.**
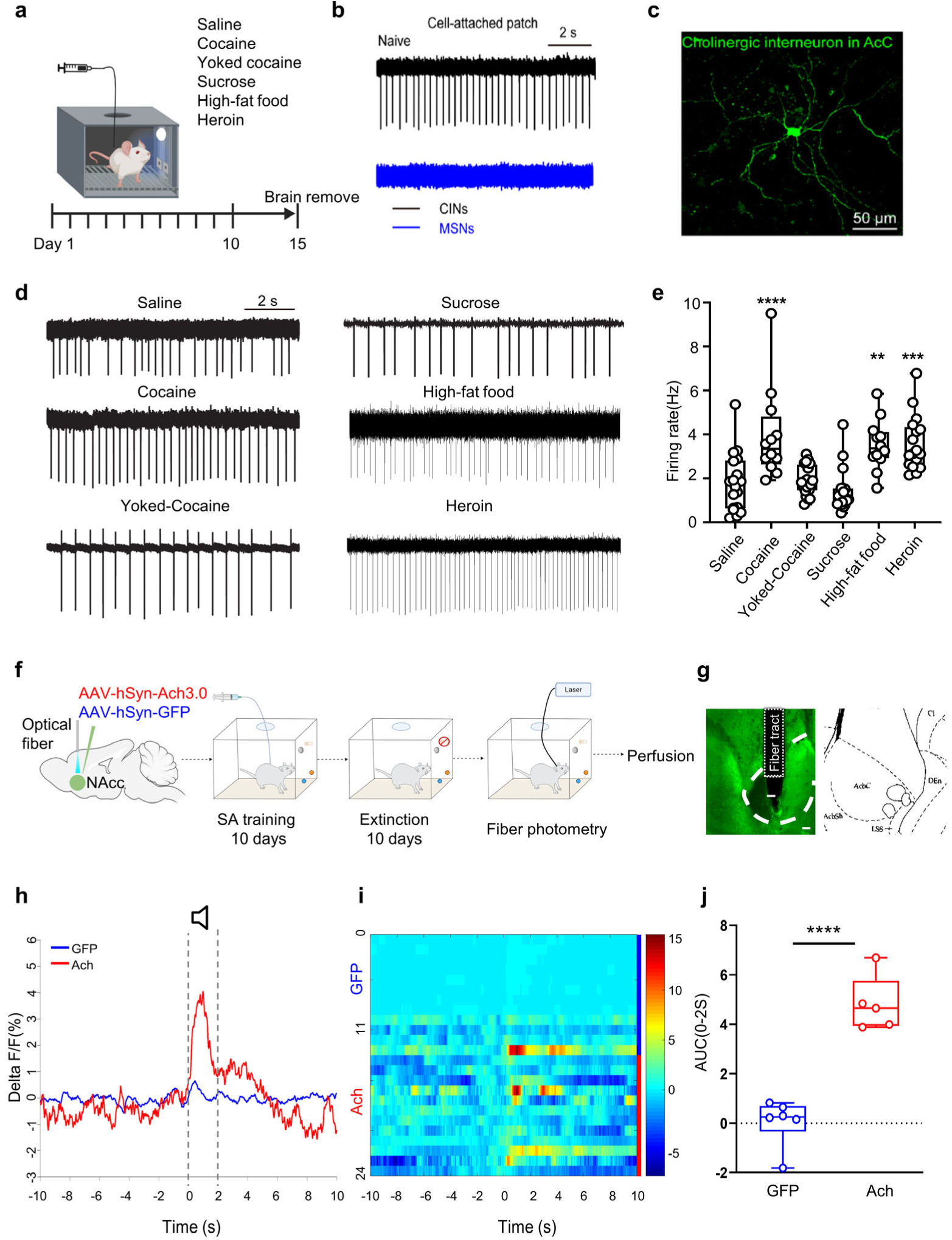
Addictive drug and high-fat self-administration increase CIN firing and cue-evoked ACh release in the NAcC. **(a)** Experimental timeline for self-administration (SA) training and ex vivo recordings across different reinforcers. **(b)** Representative voltage-clamp traces recorded at 0 mV in gap-free mode from neurons (CINs, top; MSNs, bottom). **(c)** Neurobiotin-labeled cholinergic interneurons in the NAcC, showing characteristic morphological features. **(d)** Representative firing traces of CINs recorded under different training conditions (Saline SA; Cocaine SA; Yoked cocaine; Sucrose SA; High-fat food SA; Heroin SA). **(e)** Firing rates of CINs in different groups. n = 18, 12, 20, 21, 12 and 17 cells for saline SA, cocaine SA, yoked cocaine, sucrose SA, high-fat food SA and heroin SA, respectively. **(f)** Schematic of *in vivo* recording of acetylcholine release from cholinergic interneurons in the NAcC. **(g)** Illustration of fiber photometry implant sites and viral expression. Scale bar = 200 µm. **(h and i)** Fiber photometry recordings of ACh3.0/GFP signals **(h)** and corresponding heatmap **(i)**, with Time 0 indicating drug-associated cue presentation. **(j)** AUC of z-scored ACh3.0 signals in NAcC CINs during 0-2 s after cue onset. n = 6 rats for GFP and 5 rats for ACh3.0. Data are presented as mean ± s.e.m. \*\**P* < 0.01, \*\*\**P* < 0.001, \*\*\*\**P* < 0.0001 by one-way ANOVA with Dunnett’s multiple-comparison test (e) or unpaired *t* test (j). For statistical details, see Supplementary Table 1.

To test whether this aberrant CIN firing generalizes across reinforcers, we next examined CIN firing under different reinforcement conditions. Rats were trained on heroin SA, sucrose SA, or high-fat food SA under procedures analogous to the cocaine paradigm, followed by the same withdrawal interval before recordings. CINs from heroin SA and high-fat SA rats also displayed elevated pacemaker firing compared with saline SA animals, whereas sucrose SA did not alter CIN excitability (**Fig. 1d,e**). These data indicate that self-administration of addictive drugs and high-fat reward, but not passive drug exposure or sucrose reward, selectively induces a persistent high-firing CIN state in the NAc.

### Cocaine-associated cues evoke acetylcholine transients selectively in NAc core

Because CINs provide the primary source of ACh in the NAc, we next asked how ACh release is modulated during cue-induced drug seeking. To monitor ACh dynamics *in vivo*, we expressed the genetically encoded ACh sensor AAV-hSyn-ACh3.0 or a GFP control in the NAc core (NAcC) or shell (NAcSh). Rats underwent 10 days of cocaine SA followed by extinction training, during which nose pokes no longer produced cocaine infusion but associated cues were still presented, and were subsequently tested in a cue-induced reinstatement session in which cocaine-paired cues were presented in the absence of drug delivery (**Fig. 1f and Extended Data Fig. 1a**). We also confirmed localized fluorescence within the injection site (**Fig. 1g and Extended Data Fig. 1b**). In NAcC but not NAcSh, cocaine-paired cue presentation elicited a rapid increase in ACh fluorescence (**Fig. 1h,i and Extended Data Fig. 1c,d**). Quantification of fluorescence changes in the 0-2 s window after cue onset revealed robust cue-evoked ACh transients in the ACh3.0 group relative to GFP controls in the NAcC (**Fig. 1j**) but not NAcSh (**Extended Data Fig. 1e**). Thus, drug-associated cues evoke rapid ACh release preferentially in the NAcC during cocaine seeking.

### NAc core CINs are required for cue-induced cocaine seeking and associated MSN synaptic potentiation

To determine whether NAc CINs are required for cue-induced cocaine seeking, we depleted these neurons using an intra-NAcC injection of an anti-ChAT IgG-saporin immunotoxin^37^ after extinction training (**Fig. 2a**). Two weeks after infusion, rats were tested in a cue-induced reinstatement session. Vehicle-treated rats showed robust recovery of active nose poking in response to cocaine-paired cues, whereas saporin-treated rats displayed a pronounced attenuation of cue-induced cocaine seeking; inactive nose pokes were unchanged in both groups (**Fig. 2b**). Immunostaining confirmed a marked loss of ChAT-positive neurons in the NAcC of saporin-treated rats relative to vehicle controls (**Fig. 2c**). These findings indicate that NAcC CINs are necessary for the expression of cue-induced cocaine seeking.

**Fig. 2.**
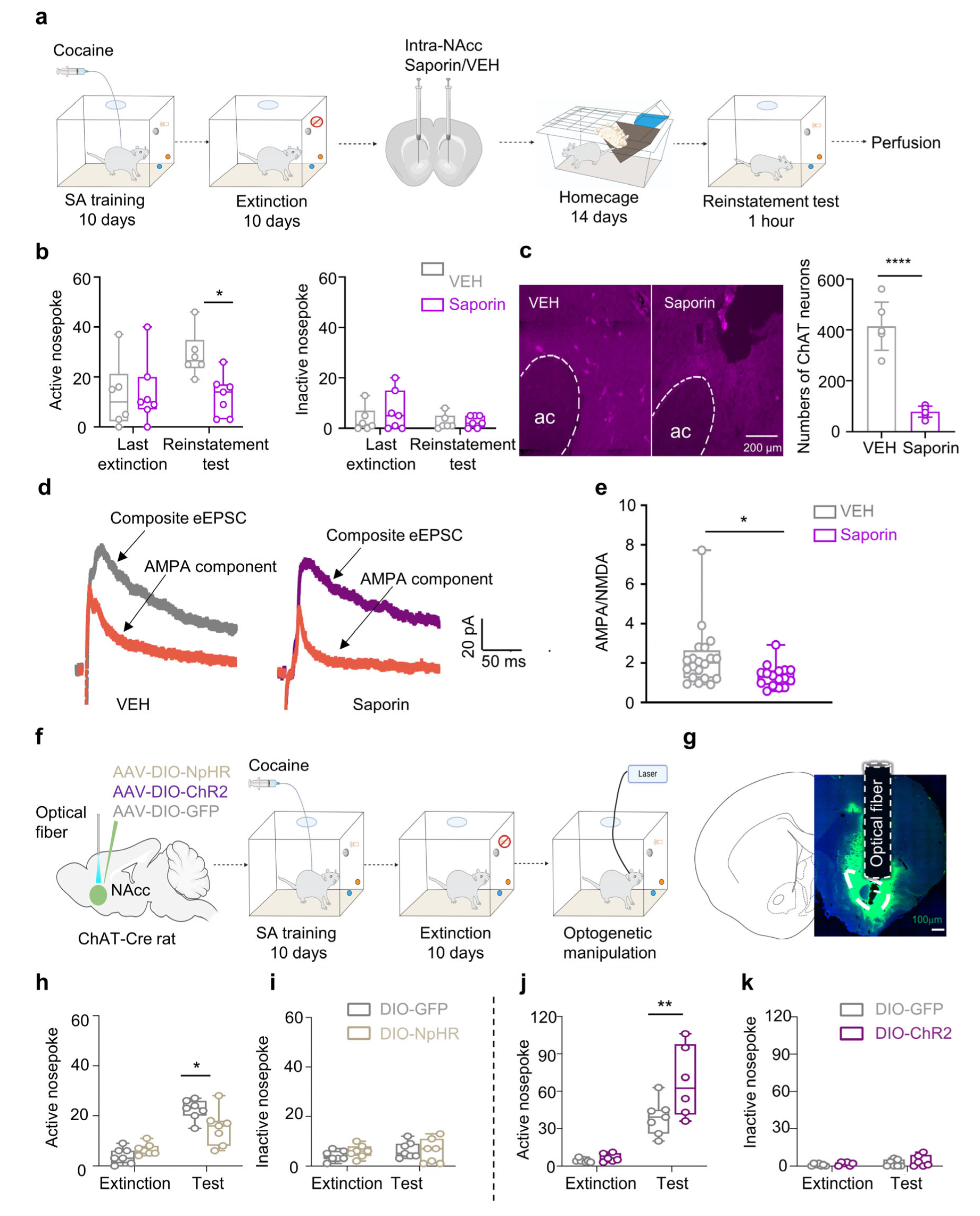
NAcC CINs are required for cue-induced cocaine seeking and associated MSN synaptic potentiation. **(a)** Experimental timeline for CIN depletion in the NAcC and subsequent behavioral and electrophysiological testing. **(b)** Number of active and inactive nose poke responses on the last day of extinction (Last extinction) and during the reinstatemen*t* test. n = 6 rats for VEH and n = 7 rats for Saporin. **(c)** Immunofluorescence and quantification of CINs in the NAc. Scale bar = 200 µm. n = 6 rats for each group. **(d)** Sample total and AMPA current traces from each group following cue-induced reinstatemen*t* test. **(e)** AMPA/NMDA ratios in MSNs in different groups. n = 20 cells for vehicle and n = 16 cells for saporin. **(f)** Timeline of optogenetic activation or inhibition of CINs during cue-induced reinstatement. **(g)** Viral injection sites and fiber implant locations. Scale bar = 100 µm. **(h-k)** Number of active and inactive nose poke responses during reinstatement with optogenetic inhibition (h and i) or activation of CINs (j and k). n = 7 rats for each group **(h and i)**. n = 7 rats for DIO-GFP and n = 6 rats for DIO-ChR2 (j and k). Data are presented as mean ± s.e.m. \**P* < 0.05, \*\**P* < 0.01, \*\*\*\**P* < 0.0001 by two-way repeated-measures ANOVA with Sidak’s multiple-comparison test (b and h-k) or unpaired *t* test (c and e). For statistical details, see Supplementary Table 2.

Cue-induced relapse across several drugs is associated with rapid potentiation of excitatory transmission onto accumbal MSNs, often reflected by an increased AMPA/NMDA (A/N) ratio^38–40^. We therefore asked whether CIN depletion also affects this synaptic signature. Following the reinstatemen*t* test, we performed whole-cell recordings from NAcC MSNs to measure A/N ratios in vehicle- and saporin-treated rats. Consistent with previous work, vehicle-treated rats exhibited an elevated A/N ratio after cue-induced cocaine seeking, whereas CIN depletion markedly reduced this enhancement (**Fig. 2d,e**). Thus, loss of NAcC CINs not only suppresses cue-induced cocaine seeking, but also reverses the associated increase in MSN A/N ratio, suggesting that CIN activity is required to maintain both the behavioral and synaptic manifestations of cocaine craving.

### Bidirectional manipulation of NAc core CINs gates cue-induced cocaine seeking

Given that CIN firing was increased after cocaine SA and that CIN ablation reduced responding to cocaine-paired cues, we next asked whether optogenetic manipulation of CINs is sufficient to bidirectionally modulate cue-induced cocaine seeking. To this end, we injected Cre-dependent AAVs encoding ChR2-, NpHR-eYFP, or GFP into the NAcC of ChAT-Cre rats (**Fig. 2f,g**). After establishing stable cocaine SA followed by extinction, rats were tested in a cue-induced reinstatement session. During the test, we delivered 473-nm blue light to activate, or 580-nm yellow light to inhibit, NAcC CINs. Optogenetic inhibition of CINs decreased active nose pokes in response to cocaine-paired cues (**Fig. 2h**), whereas optogenetic activation of CINs increased active responding (**Fig. 2j**). Inactive nose pokes remained unchanged across groups during optogenetic manipulation (**Fig. 2i,k**). Furthermore, optogenetic inhibition of CINs in NAcSh had no effect on cue-induced cocaine seeking behavior (**Extended Data Fig. 2**). These findings demonstrate that NAcC CIN activity is sufficient to bidirectionally gate cue-induced cocaine seeking.

### CIN-specific ISR signaling sustains cocaine-induced hyperexcitability and seeking

The integrated stress response (ISR), indexed by phosphorylation of eIF2α, regulates translational programs that support long-term memory, synaptic plasticity and neuronal excitability ^41–43^. Because ISR activity in striatal CINs has also been linked to learning-related changes in excitability ^34^, we asked whether cocaine recruits ISR signaling in NAc CINs. Cocaine SA, but not sucrose SA, increased p-eIF2α and ATF4 in CINs and also elevated p-eIF2α in D1- and D2-expressing neurons (**Extended Data Fig. 3a-h**), indicating that cocaine engages ISR signaling across multiple accumbal cell populations.

To test whether ISR activation is required for cocaine-induced CIN remodeling, rats received intra-NAc AAV-hSyn-ACh3.0 and daily treatment with ISRIB or vehicle during cocaine SA, followed by extinction and cue-induced reinstatement with simultaneous fiber photometry (**Fig. 3a**). ISRIB reduced active nose pokes during cue-induced reinstatement (**Fig. 3b**), lowered CIN firing ex vivo (**Fig. 3c,d**), and markedly attenuated cocaine cue-evoked ACh transients (**Fig. 3e-g**). Thus, ISR activity during cocaine experience is required for the emergence of CIN hyperexcitability and abnormal cue-evoked cholinergic signaling.

**Fig. 3.**
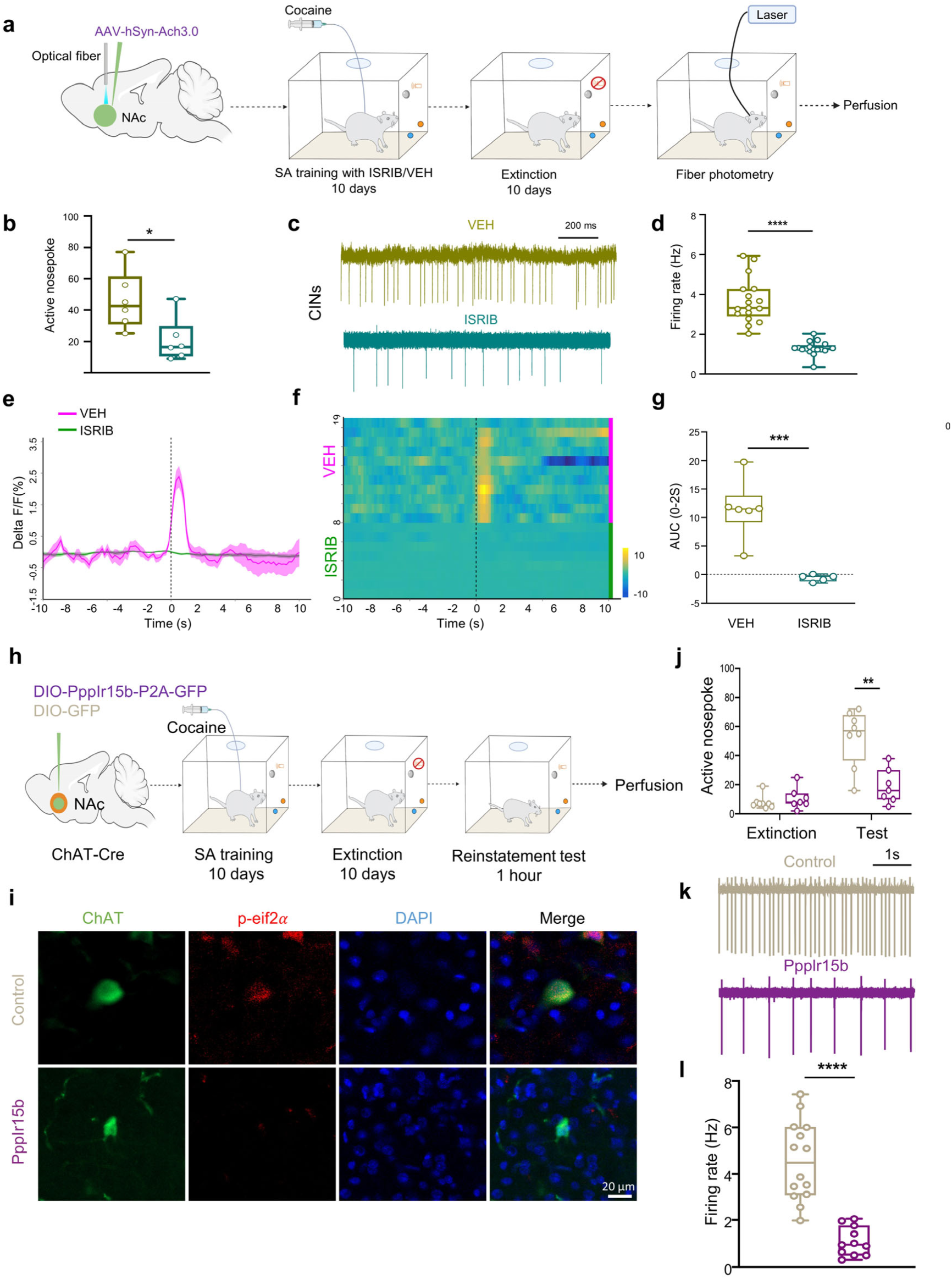
CIN-specific ISR signaling sustains cocaine-induced hyperexcitability and seeking. **(a)** Experimental timeline showing ISRIB or vehicle treatment during cocaine SA training, followed by extinction and cue-induced reinstatement. **(b)** Number of active nose poke responses during the cue-induced reinstatemen*t* test. N = 6 rats for each group. **(c and d)** Representative spontaneous firing traces of CINs (c) and quantification of CIN firing rates (d). n = 17 cells from 3 rats per group. **(e-g)** ACh3.0 photometric responses aligned to cocaine-cue onset: temporal dynamics of ΔF/F signals (e), heatmaps of trial-aligned signals (f), and AUC values over the 0-2 s post-cue interval (g). n = 6 rats for VEH and n = 5 rats for ISRIB. **(h)** Experimental timeline for CIN-specific ISR suppression using Cre-dependent Ppp1r15b expression in ChAT-Cre rats. **(i)** Immunofluorescence validation of CIN-restricted reduction of p-eIF2α after Ppp1r15b expression. Green, ChAT; red, p-eIF2α; blue, DAPI. Scale bar = 20 µm. **(j)** Number of active nose poke responses on the last day of extinction and during cue-induced reinstatement. n = 8 rats for DIO-GFP and n = 7 rats for DIO-Ppp1r15b-P2A-GFP. **(k and l)** Representative CIN firing traces (k) and quantification of firing rates (l). n = 14 cells from 4 rats for control and n = 11 cells from 3 rats for Ppp1r15b. Data are presented as mean ± s.e.m. \**P* < 0.05, \*\**P* < 0.01, \*\*\**P* < 0.001, \*\*\*\**P* < 0.0001. Statistical tests are as specified in Supplementary Table 3.

We next tested whether elevating ISR tone is sufficient to transform ordinary natural reward seeking into maladaptive cue-driven behavior. Sal003 administration during sucrose SA did not increase cue-induced sucrose seeking, but increased CIN firing and disrupted the normal sucrose cue-associated ACh pause (**Extended Data Fig. 4a-g**). Immunofluorescence confirmed that ISRIB and Sal003 bidirectionally regulated p-eIF2α levels in NAc CINs (**Extended Data Fig. 5**). These findings indicate that ISR elevation is sufficient to reshape CIN physiology and cue-evoked ACh dynamics, but is not by itself sufficient to enhance sucrose seeking.

To establish cell-type-specific causality, we selectively suppressed ISR signaling in CINs using ChAT-Cre rats and a Cre-dependent AAV expressing the eIF2α phosphatase regulatory subunit Ppp1r15b ^46,47^ (**Fig. 3h**). Ppp1r15b expression reduced p-eIF2α selectively in NAc ChAT-positive neurons (**Fig. 3i**) and markedly attenuated cue-induced cocaine seeking (**Fig. 3j**). CIN-specific ISR suppression did not alter cocaine acquisition or extinction, and inactive responding remained unchanged (**Extended Data Fig. 7a-c**), indicating that the reduction in cue-induced seeking was not explained by impaired drug intake, extinction learning or generalized behavioral suppression. To assess cell-type specificity further, we performed parallel Ppp1r15b manipulations in D1- and D2-expressing neurons. In contrast to CINs, ISR suppression in either D1R- or D2R-expressing neurons did not alter cocaine self-administration, extinction or cue-induced reinstatement (**Extended Data Fig. 6a-l**).

Finally, CIN-specific Ppp1r15b expression markedly reduced spontaneous CIN firing compared with controls (**Fig. 3k,l**), indicating that ISR suppression reverses the cocaine-associated hyperexcitable CIN state. Together, these results identify ISR signaling in NAc CINs as a functionally selective mechanism that sustains cocaine-associated CIN hyperexcitability and cue-driven seeking.

### Cocaine engages an ISR-sensitive HCN program in CINs

To identify the intrinsic conductances that support NAc CIN pacemaker activity, we recorded spontaneous firing while pharmacologically blocking distinct ion-channel classes (**Fig. 4a**). ZD7288, an HCN channel antagonist, markedly reduced spontaneous CIN firing, whereas blockade of T-type or L-type calcium channels, BK, SK or KCNQ potassium channels, or fast excitatory and inhibitory synaptic transmission did not produce comparable effects; TTX abolished firing as expected (**Fig. 4a-c**). In cocaine-trained rats, ZD7288 reduced the elevated firing rate toward saline-control levels (**Fig. 4d-f**), indicating that HCN conductance is required to maintain cocaine-associated CIN hyperexcitability.

**Fig. 4.**
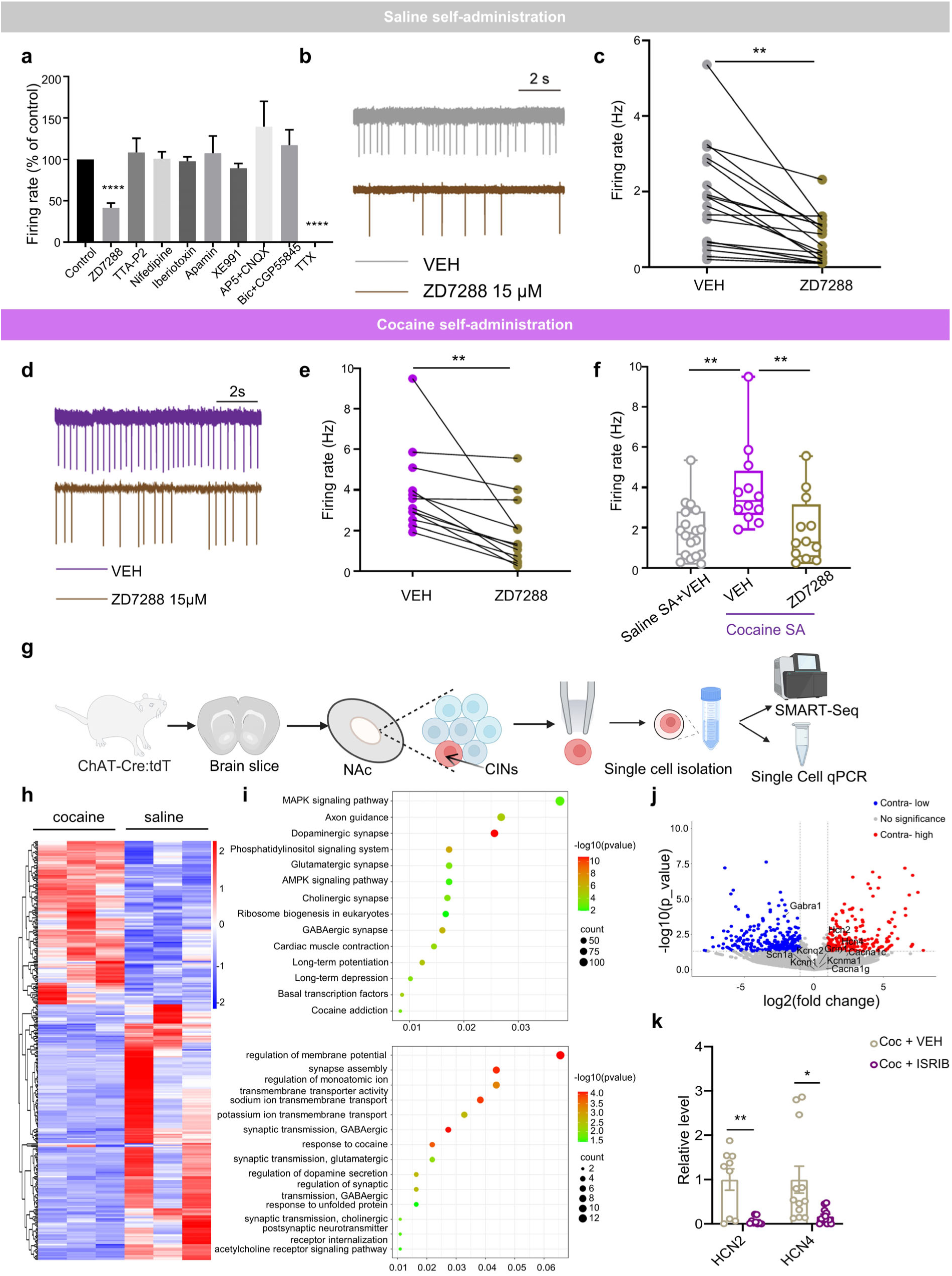
Cocaine engages an ISR-sensitive HCN program in CINs. **(a)** Ex vivo CIN firing recorded in NAc slices from saline SA rats following drug application. n = 19, 18, 7, 7, 8, 8, 7, 7, 7 and 5 cells for control, ZD7288, TTA-P2, nifedipine, iberiotoxin, apamin, XE991, AP5 + CNQX, bicuculline + CGP55845 and TTX, respectively. **(b)** Representative CIN firing traces before (top) and after (bottom) ZD7288. **(c)** Quantification of CIN firing rates before and after ZD7288. n = 18 cells for each group. **(d)** Representative CIN firing traces from cocaine SA rats before and after ZD7288. **(e)** Quantification of CIN firing rates before and after ZD7288. n = 12 cells for each group. **(f)** ZD7288-induced changes in CIN firing in saline SA versus cocaine SA groups. n = 18 cells for Saline SA + VEH, n = 12 cells for Cocaine SA + VEH and n = 12 cells for Cocaine SA + ZD7288. **(g)** SMART-Seq and qPCR timeline schematic. **(h-j)** Single-cell transcriptomic profiling of NAc CINs: Heatmaps of DEGs between cocaine and saline groups, *P* < 0.05; n = 3 per group **(h)**; KEGG (top) and GO (bottom) enrichment of DEGs **(i)**; and volcano plot highlighting DEGs and ion channel-related genes **(j)**. **(k)** Relative mRNA levels of HCN2 and HCN4 in ISRIB- and vehicle-treated groups. n = 9 cells for each group (left) and n = 12 cells for each group (right). Data are presented as mean ± s.e.m. \**P* < 0.05, \*\**P* < 0.01 by one-way ANOVA with Dunnett’s multiple-comparison test (a and f) or paired *t* test (c and e) or unpaired *t* test (k). For statistical details, see Supplementary Table 4.

To identify molecular effectors underlying the cocaine-induced CIN hyperexcitable state, we individually collected NAcC ChAT-positive neurons from ChAT-Cre:tdTomato rats after saline or cocaine SA and performed single-cell RNA sequencing (**Fig. 4g**). Cocaine SA produced 292 upregulated and 258 downregulated genes, with enrichment of pathways related to membrane potential regulation and ion transport (**Fig. 4h,i**). Among ion-channel-related genes, HCN2 and HCN4 were increased after cocaine SA (**Fig. 4j**), and single-cell qPCR showed that ISRIB treatment reduced HCN2 and HCN4 mRNA levels (**Fig. 4k**). Pharmacological manipulation of ISR signaling also bidirectionally altered HCN2 protein expression in CINs (**Extended Data Fig. 8a-c**), supporting an ISR-sensitive HCN expression program.

We next measured hyperpolarization-activated currents (*I*_h_), the characteristic current mediated by HCN channels^48^. Consistent with these molecular findings, cocaine training increased hyperpolarization-activated current (*I*_h_) in CINs. Cocaine-trained CINs showed larger maximal HCN currents and higher current density at strongly hyperpolarized potentials, as well as increased available HCN current closer to the physiological voltage range (**Extended Data Fig. 9a-i**). Together, these data indicate that cocaine experience recruits an ISR-sensitive HCN program that strengthens HCN-dependent intrinsic excitability in NAc CINs.

### HCN2 is the CIN-specific causal effector of cue-induced cocaine seeking

We next asked which HCN subtype mediates the cocaine-associated CIN state. HCN channels, encoded by four genes (HCN1-4), are widely expressed in the heart and central nervous system, but HCN3 is sparsely distributed at very low levels in the central nervous system ^49^. HCN1 expression did not differ across reinforcement conditions, whereas HCN2 was increased in cocaine SA and yoked-cocaine groups and HCN4 was selectively increased after cocaine SA (**Extended Data Fig. 10a-d**). These findings suggested that HCN2 and HCN4 are recruited by cocaine exposure, but did not establish which isoform is functionally required.

To test isoform specificity, we injected lentiviral RNAi constructs targeting HCN1, HCN2 or HCN4 into the NAc after cocaine training (**Fig. 5a**). HCN2 knockdown produced the clearest reduction in HCN-dependent current across the activation protocols (**Fig. 5b-g**), decreased CIN firing (**Fig. 5h**), and reduced active responding during cue-induced cocaine seeking (**Fig. 5i**). In contrast, HCN1 or HCN4 knockdown had little effect on CIN firing or cocaine seeking. Western blotting confirmed selective knockdown of HCN1, HCN2 and HCN4, respectively (**Extended Data Fig. 11a-c**).

**Fig. 5.**
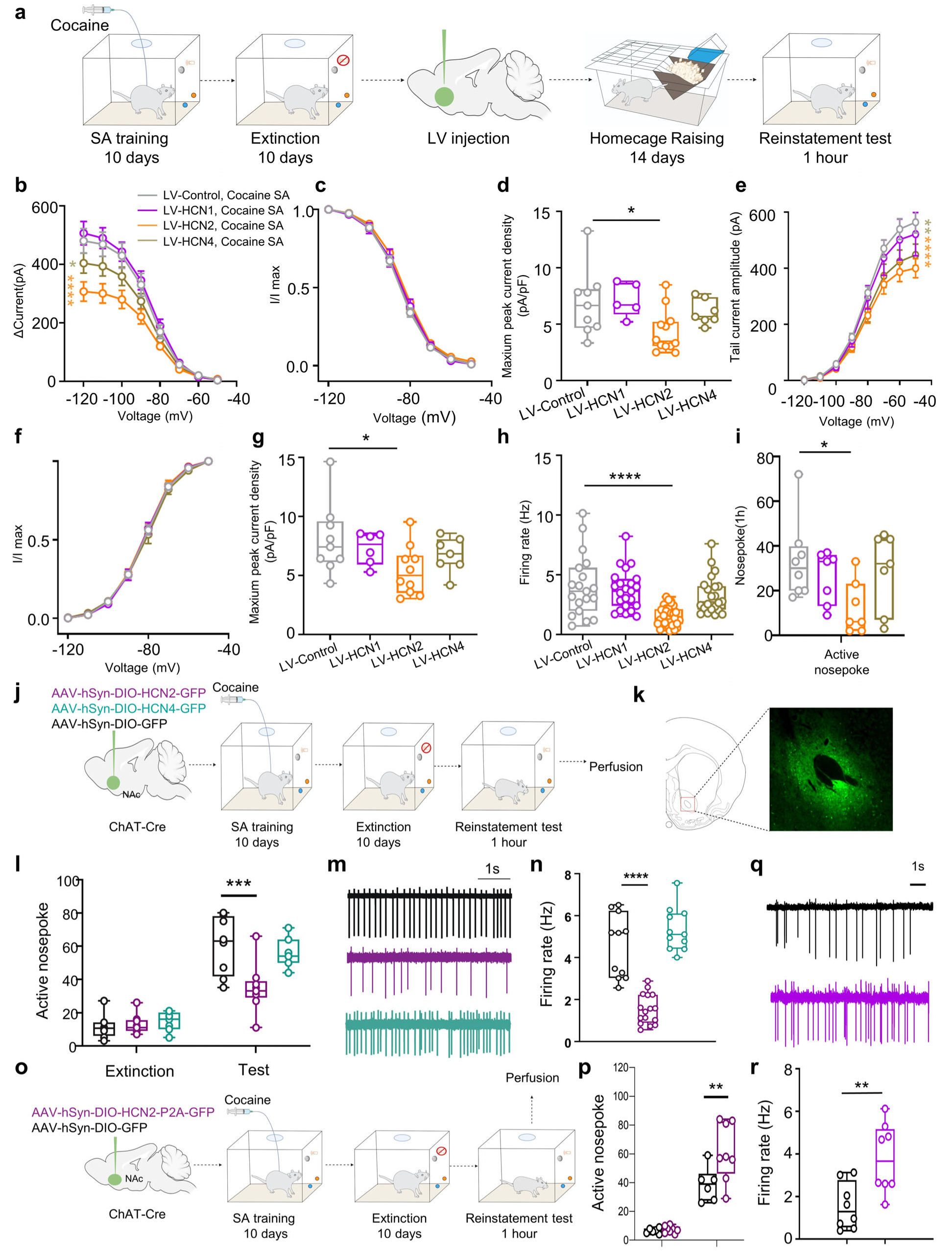
HCN2 is the CIN-specific causal effector of cue-induced cocaine seeking. **(a)** Schematic timeline for post-training knockdown of HCN1, HCN2 or HCN4 in the NAc. **(b-g)** HCN current analyses following HCN1, HCN2 or HCN4 knockdown: steady-state activation curves (b), normalized activation curves (c), peak current-density quantification (d), tail-current activation curves (e), normalized curves corresponding to (e) (f), and peak current-density summary (g). **(h)** Quantification of CIN firing rates after HCN isoform knockdown. n = 19, 23, 33 and 22 cells for LV-Control, LV-HCN1, LV-HCN2 and LV-HCN4 groups, respectively. **(i)** Number of active nose poke responses during cue-induced reinstatement. n = 8, 7, 8 and 7 rats for LV-Control, LV-HCN1, LV-HCN2 and LV-HCN4 groups, respectively. **(j and k)** Experimental timeline for CIN-specific HCN2 or HCN4 knockdown (j) and validation of viral expression in NAc CINs (k). **(l)** Number of active nose poke responses on the last day of extinction and during cue-induced reinstatement after CIN-specific HCN2 or HCN4 knockdown. n = 8 rats for DIO-GFP, n = 9 rats for DIO-HCN2-shRNA and n = 7 rats for DIO-HCN4-shRNA. **(m and n)** Representative spontaneous CIN firing traces (m) and quantification of firing rates (n). n = 11, 16 and 11 cells for DIO-GFP, DIO-HCN2-shRNA and DIO-HCN4-shRNA groups, respectively. **(o)** Experimental timeline for CIN-specific HCN2 overexpression. **(p)** Number of active nose poke responses on the last day of extinction and during cue-induced reinstatement after HCN2 overexpression. n = 6 rats for DIO-GFP and n = 8 rats for DIO-HCN2-P2A-GFP. **(q and r)** Representative CIN firing traces (q) and quantification of firing rates (r). n = 8 cells for each group. Data are presented as mean ± s.e.m. \**P* < 0.05, \*\**P* < 0.01, \*\*\**P* < 0.001, \*\*\*\**P* < 0.0001. Statistical tests are as specified in Supplementary Table 5.

We then tested whether HCN2 specifically within CINs is necessary and sufficient for this behavioral state. CIN-specific HCN2 knockdown, but not HCN4 knockdown, reduced cue-induced cocaine seeking (**Fig. 5j-l**) and decreased CIN firing (**Fig. 5m,n**). Conversely, CIN-restricted HCN2 overexpression increased cue-induced cocaine seeking (**Fig. 5o,p**) and increased CIN firing (**Fig. 5q,r**). Thus, HCN2 in CINs is both necessary and sufficient to regulate the hyperexcitable CIN state that promotes cue-induced cocaine seeking.

### Local and systemic HCN blockade suppress maladaptive cue-driven seeking

Finally, we tested whether HCN channels provide a pharmacologically tractable target across reinforcers. After cocaine SA and extinction, bilateral intra-NAcC infusion of ZD7288 markedly reduced cue-induced cocaine seeking (**Fig. 6a,b**). In the same cocaine paradigm, local HCN blockade prevented the cue-associated increase in AMPA/NMDA ratio in NAcC MSNs (**Fig. 6c,d**). In heroin-trained rats, ZD7288 reduced cue-induced heroin seeking (**Fig. 6e**) and normalized the elevated firing of CINs recorded ex vivo (**Fig. 6f,g**). Local HCN blockade also reduced cue-induced high-fat reward seeking (**Fig. 6h**).

**Fig. 6.**
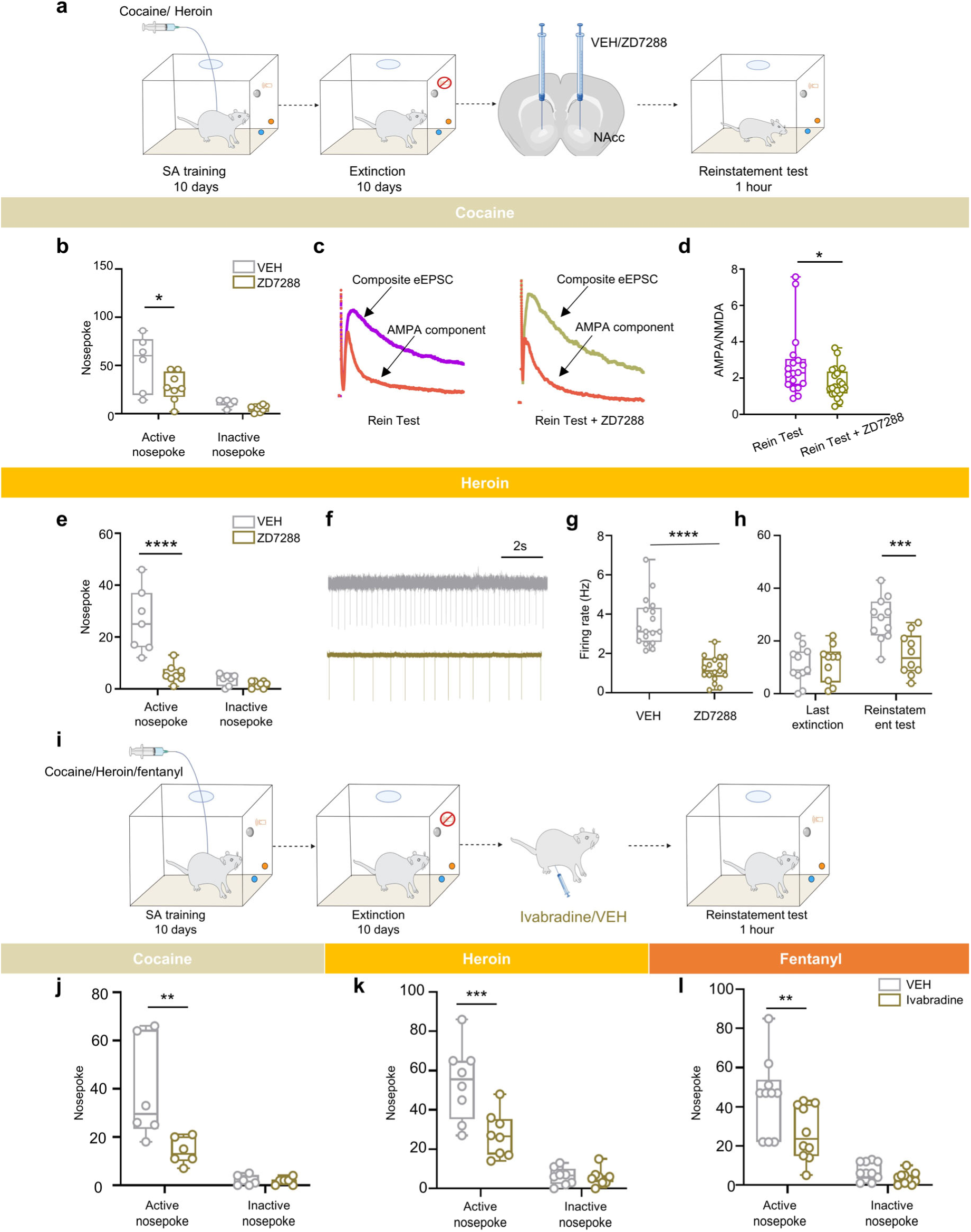
Local and systemic HCN blockade suppress maladaptive cue-driven seeking and associated accumbal plasticity. **(a)** Schematic timeline for bilateral intra-NAcC infusion of ZD7288 before cue-induced reinstatement. **(b)** Active and inactive nose poke responses during cue-induced cocaine reinstatement. n = 6 rats for VEH and n = 8 rats for ZD7288. **(c and d)** Representative total and AMPA current traces from NAcC MSNs after cue-induced cocaine reinstatement (c) and quantification of the AMPA/NMDA ratio (d). n = 18 cells for VEH and n = 21 cells for ZD7288. **(e)** Active and inactive nose poke responses during cue-induced heroin reinstatement. N = 7 rats for VEH and n = 8 rats for ZD7288. **(f and g)** Representative spontaneous CIN firing traces from heroin-trained rats following VEH or ZD7288 treatment (f) and quantification of CIN firing frequency (g). n = 17 cells for each group. **(h)** Active nose poke responses in high-fat-food trained rats on the last extinction session and during cue-induced reinstatement. n = 11 rats for VEH and n = 10 rats for ZD7288. **(i)** Experimental timeline for systemic ivabradine treatment before cue-induced reinstatement. **(j-l)** Active and inactive nose poke responses following systemic ivabradine treatment in cocaine-trained (j), heroin-trained (k), and fentanyl-trained (l) rats. For cocaine, n = 6 VEH and n = 7 ivabradine rats; for heroin, n = 8 rats per group; for fentanyl, n = 10 rats per group. Data are presented as mean ± s.e.m. \**P* < 0.05, \*\**P* < 0.01, \*\*\**P* < 0.001, \*\*\*\**P* < 0.0001. Statistical tests are as specified in Supplementary Table 6.

The local effect generalized further to fentanyl: intra-NAcC ZD7288 suppressed cue-induced fentanyl seeking (**Extended Data Fig. 12a-c**). In high-fat trained rats, ZD7288 normalized elevated CIN firing (**Extended Data Fig. 12e,f**), whereas sucrose seeking was unaffected (**Extended Data Fig. 12d,g**). Intra-NAcSh ZD7288 did not alter cue-induced cocaine or sucrose seeking (**Extended Data Fig. 13a-f**), supporting both reinforcer and anatomical specificity of the NAcC HCN-dependent effect.

Because direct intra-NAc infusion is not clinically practical, we next tested systemic ivabradine, a clinically approved HCN channel blocker ^50,51^ (**Fig. 6i**). Ivabradine reduced cue-induced cocaine-, heroin- and fentanyl-seeking without altering inactive responding (**Fig. 6j-l**). Together with the local infusion and genetic data, these findings identify HCN-dependent excitability as a tractable intervention point for reducing cue-induced relapse-like behavior across multiple addictive drugs.

## Discussion

This study identifies nucleus accumbens CINs as key regulators of maladaptive cue-driven motivation and defines an ISR-HCN2 signaling axis that sustains this state. Using complementary optogenetic, fiber photometry, electrophysiological, genetic, and pharmacological approaches, we show that self-administration of addictive drugs and high-fat reward, but not sucrose self-administration, induces persistent CIN hyperexcitability. Bidirectional manipulation of NAc core CINs suppresses or enhances cue-induced cocaine seeking, respectively. We further found that this state depends on ISR signaling and on HCN channels, particularly HCN2, in NAc CINs. Finally, CIN-specific HCN2 knockdown or pharmacological HCN blockade, including systemic treatment with the FDA-approved HCN blocker ivabradine, reduces cue-induced seeking across multiple addictive drugs and high-fat reward while sparing sucrose seeking. These findings are summarized in the proposed model shown in **Extended Data Fig. 14**. Together, these findings reveal a cholinergic mechanism by which addictive and highly palatable reward experience can be converted into persistent maladaptive cue motivation.

### CINs sustain maladaptive cue-driven motivation

Previous work has established NAc CINs as powerful modulators of cue-guided behavior. Under physiological conditions, CINs shape conditioned cue-response relationships, regulate striatal output, and oppose excessive cue-motivated responding ^29,52^. This suppressive role is thought to prevent excessive cue-driven responding and maintain behavioral flexibility ^52^. More recent studies further suggest that CIN-derived acetylcholine can modulate dopamine release in a task- and state-dependent manner, supporting effortful reward-directed actions^30^. Our findings extend this framework by showing that CINs are not only adaptive regulators of cue-guided behavior but can also become substrates of maladaptive cue motivation after addictive experience. Rather than simply responding to cues, NAc core CINs enter a persistent high-firing state after cocaine or heroin self-administration and after high-fat reward self-administration, but not after passive cocaine exposure or sucrose self-administration. The causal data further indicate that this CIN state has functional significance. Depletion or optogenetic inhibition of NAc core CINs reduces cue-induced cocaine seeking, whereas optogenetic activation enhances it. CIN depletion also prevents the cue-associated increase in MSN AMPA/NMDA ratio, linking CIN activity to a synaptic signature of relapse-like behavior. This finding is consistent with previous work showing that reinstatement of drug seeking is associated with rapid glutamatergic potentiation in the NAc^38,39^. Thus, NAc core CINs act as state-setting regulators that sustain both the behavioral expression of cue-induced seeking and the local synaptic potentiation associated with relapse-like behavior. Future studies should determine how CIN-derived acetylcholine acts on specific downstream receptors and synapses, including dopamine terminals, glutamatergic inputs, GABAergic microcircuits, and MSN subtypes, to transform a high-firing CIN state into persistent cue-driven action.

Dopamine has long been central to theories of reward prediction, reinforcement learning, and incentive motivation^9,53,54^. These models have provided a powerful framework for understanding how drug-associated cues acquire motivational value. However, phasic dopamine transients alone may not fully account for the persistence, cue selectivity, and state dependence of relapse vulnerability after prolonged abstinence. Our study adds a complementary cholinergic mechanism by showing that voluntary drug-taking experience produces lasting changes in NAc CIN excitability and cue-evoked acetylcholine release, and that acute manipulation of CIN activity bidirectionally controls cue-induced drug seeking. This does not argue against dopamine-based models. Recent work showing that dopamine and acetylcholine signals can be partly independent yet temporally coordinated across task states fits well with this interpretation^55^. A key future direction is to define when CINs act independently of dopamine and when their effects depend on dopamine-acetylcholine coupling.

### ISR signaling links addictive experience to persistent CIN hyperexcitability

The ISR is increasingly recognized as a translational control pathway that links cellular stress to synaptic plasticity, memory, and brain disease^41,56^. Previous work showed that striatal cholinergic neurons constitutively engage ISR signaling and that this pathway can influence dopamine modulation and learning-related adaptation^34^. Our data extend these findings by placing ISR signaling within NAc CINs as a mechanism for persistent maladaptive motivation. We found that pharmacological or genetic inhibition of ISR signaling reduces CIN excitability and cue-induced cocaine seeking, indicating that ISR activity is required to maintain the cocaine-associated hyperexcitable CIN state. Notably, cocaine, but not sucrose, increased ISR markers in CINs, suggesting that ISR functions as a molecular switch that sustains the hyperactive state of CINs in addiction. Stress is a significant risk factor for both the initiation of addiction and the propensity for relapse^2,57^. Diverse stressors can converge on ISR kinases that phosphorylate eIF2α and reprogram translation, thereby altering neuronal excitability and plasticity. In this context, our findings suggest that ISR activation in CINs is a candidate mechanistic link between stress vulnerability and addiction relapse.

Our findings also refine how ISR signaling should be interpreted in reward learning. Elevating ISR tone during sucrose self-administration increased CIN firing and altered sucrose cue-evoked acetylcholine pause dynamics, but did not enhance sucrose seeking. This suggests that ISR elevation can reshape CIN physiology but is not by itself sufficient to transform ordinary natural reward seeking into maladaptive motivation. Addictive experience likely recruits additional neuromodulatory, transcriptional, or synaptic adaptations that couple CIN hyperexcitability to persistent cue-driven behavior.

### HCN2 acts as a downstream effector of the CIN ISR program

We identify HCN2 as a critical downstream effector through which ISR signaling stabilizes CIN hyperexcitability. We show that cocaine self-administration upregulates HCN2 and HCN4 expression in NAc CINs, enhances *I*_h_, and produces a persistently elevated pacemaker state. Pharmacological HCN blockade normalizes CIN firing and suppresses cue-induced drug seeking, and convergent viral manipulations demonstrate isoform and cell-type specificity: knockdown of HCN2, but not HCN4, reduces CIN excitability and cocaine seeking, whereas HCN2 overexpression has the opposite effect. Together, these findings indicate that HCN2 functions as a principal effector through which cocaine experience stabilizes a high-gain CIN state that amplifies drug-cue-evoked ACh release.

More broadly, converging evidence indicates that HCN isoforms, especially HCN1, HCN2 and HCN4, contribute to higher cognition, affect regulation, and several neuropsychiatric conditions. In cortical and hippocampal circuits, cAMP-HCN signaling constrains persistent firing and synaptic integration, and reducing HCN function can improve stress-related cognitive deficits^58,59^. Within reward- and mood-related circuits, HCN2 in ventral tegmental area dopamine neurons and in NAc CINs has been shown to regulate depressive- and anxiety-like behaviors as well as reward-evoked dopamine release ^60,61^. HCN channels have also been implicated in addiction-related behaviors ^62,63^, including HCN-dependent rhythmicity in medial habenula cholinergic neurons during nicotine withdrawal ^63^. Extending these observations, our data provide cell-type- and isoform-specific evidence that an ISR-sensitive program upregulates HCN2 in NAc CINs, strengthens *I*_h_, and stabilizes a hyperexcitable CIN state that amplifies drug-cue-evoked ACh release, while CIN-restricted HCN2 manipulations bidirectionally control cue-induced drug seeking. These findings suggest a broader principle by which neuromodulatory systems maintain persistent motivational states. The ISR-HCN pathway in CINs may represent a generalizable mechanism for sustained maladaptive motivation.

This study identifies accumbal CINs, together with their ISR-HCN2 signaling axis, as key regulators of maladaptive cue-driven motivation. By showing that cocaine experience recruits ISR-dependent upregulation of HCN2 in CINs to sustain a hyperexcitable, cue-responsive state, our findings extend dopamine-focused models of addiction to include a complementary, cholinergic and cell-type-specific mechanism for persistent cue reactivity. These results highlight the ISR-HCN2 pathway as a promising target for relapse prevention and demonstrate that pharmacological HCN blockade with an approved agent can attenuate cue-induced drug seeking, suggesting a concrete translational strategy for modulating pathological cue-driven motivation while sparing basic reward processing in our paradigms.

## Methods

### Animals

All procedures were approved by the Biomedical Ethics Committee of animal use and protection of Peking University and were performed in accordance with the National Institutes of Health Guide for the Care and Use of Laboratory Animals. Rats were housed in a reverse light cycle room (light period: 8 pm to 8 am) under controlled temperature (23 ± 2 °C) and humidity (50 ± 5%). After surgery, rats were separated alone with access to food and water ad libitum. Male Sprague-Dawley (SD) rats purchased from the Vital River Company, D1-Cre, D2-Cre, acetyltransferase (ChAT): Cre+ transgenic and ChAT/tdTomato rats were used for study.

### Viruses

The following AAVs and LVs were used in this study: For optogenetic inhibition of CINs, AAV-pEF1α-DIO-NpHR-GFP was used, whereas AAV-pEF1α-DIO-ChR2-eYFP was used for optogenetic activation of CINs (1 × 10^13^ vg/ml). For fiber photometry recordings of acetylcholine dynamics, AAV9-hSyn-ACh3.0 was employed. To selectively reduce eIF2α phosphorylation in CINs within the NAc, pAV-hSyn-DIO-Ppp1r15b-P2A-GFP or the corresponding control virus pAV-hSyn-DIO-P2A-GFP was used (1 × 10^13^ vg/ml). In addition, to selectively reduce Hcn2 and Hcn4 mRNA levels in CINs, AAV9-CAG-DIO-GFP-3in1mir30shRNA (HCN2), AAV9-CAG-DIO-GFP-3in1mir30shRNA (HCN4), or pAV-CAG-DIO-GFP-mir30shRNA were used (1 × 10^13^ vg/ml). All viral vectors were purchased from Shandong Weizhen Biotechnology Co., Ltd., or were provided by the Yulong Li Laboratory, Peking University.

### Chemicals

Cocaine hydrochloride and heroin hydrochloride were purchased from Qinghai Pharmaceutical Factory. Fentanyl was purchased from China National Pharmaceutical Industry Corporation Ltd. Saporin (Advanced Targeting Systems Inc., ChAT antibody catalog no. AB-N34ap) was dissolved in 0.01 M PBS at a final concentration of 0.5 µg/µl. ISRIB (1.25 mg/kg, i.p.), Sal003 (2.5 mg/kg, i.p.), ZD7288 (0.625 µg/µl for intra-NAc) and ivabradine (5 mg/kg, i.p.) were purchased from MedChemExpress (USA).

The 0.2 mol/L phosphate buffer (PB, pH 7.4) was prepared by dissolving 5.93 g of NaH2PO4·2H2O and 58.02 g of Na2HPO4·12H2O in deionized water to a final volume of 1000 mL. The 30% sucrose solution was prepared with 300 g of sucrose, 500 mL of deionized water, and 500 mL of 0.2 mol/L PB. The viruses used in this experiment, including AAV-hSyn-GFP, AAV-DIO-NpHR-GFP, AAV-DIO-ChR2-GFP, AAV-DIO-EYFP, AAV9-CMV, DIO-HCN2-P2A-GFP, and AAV-CMV-DIO-GFP, were purchased from Shandong Weizhen Biotechnology Co., Ltd.

### Intravenous surgery

Based on our previous studies ^64,65^, rats were anesthetized with isoflurane (3% induction and 2% maintenance). The tip termination of the silastic catheter was placed into the jugular vein, and the other termination connecting to cannula reached above the skull through the neck. Following surgery, rats received daily injections of penicillin and heparin sodium to prevent inflammation and infection. After a 5-day recovery period, rats entered the self-administration training phase.

### Intracranial surgery

Rats were anesthetized with isoflurane for stereotaxic manipulation in a stereotaxic apparatus. The ear bar fixer was used to secure the rat’s head in a horizontal position. The connective tissue covering the skull was carefully removed, and arterial forceps were employed to expose the bony landmarks of the skull. The anterior and posterior fontanelles were adjusted to align along the same horizontal plane. Subsequently, four shallow holes were drilled into the skull surface, and small stainless-steel screws were securely implanted ^66^. For fiber photometry ^67^, AAV9-hSyn-ACh3.0 or AAV9-hSyn-EGFP were unilaterally injected into the NAc of SD rats, followed by implantation of optical fiber. For optogenetic manipulations, AAV-pEF1α-DIO-ChR2-eYFP, AAV-pEF1α-DIO-eNpHR-eYFP, or the control vector were bilaterally injected into NAc core and shell in both SD and ChAT-Cre rats at the following coordinates: NAc core (AP: +1.9 mm; ML: ±4.2 mm; DV: -6.4 mm; 16° angle) and NAc shell (AP: +1.8 mm; ML: ±0.6 mm; DV: -8.2 mm). Optical cannulae were implanted at the following coordinates: NAc core (AP: +1.9 mm; ML: ±4.2 mm; DV: -5.9 mm; 16° angle) and NAc shell (AP: +1.8 mm; ML: ±2.5 mm; DV: -7.3 mm; 10° angle). A glass pipette was attached to a syringe filled with mineral oil for virus withdrawal and infusion, and viruses were injected using a microinjection syringe pump. The injection speed was 50 nl/min and injection volumes were 500 nl. After the injection, the needle remained for additional 10 min and was lifted slowly. For cannulation, we firstly adjusted the length of the needle by 1 mm more than the cannula. We drilled two holes into the skull and guided cannula bilaterally based on coordinates of NAc core (AP: +1.9 mm; ML: ±4.2 mm; DV: -6.2 mm; 16° angle) and shell (AP: +1.8 mm, ML: ±3.2 mm, DV: -6.6 mm, 16° angle). Intracranial drug injections, including ZD7288, were performed via implanted cannulae (0.1 µg in 0.5 µl per side, infused over 2 min).

### Apparatus

Self-administration (SA) experiments were conducted in standard operant conditioning chambers (Ningbo AniLab Experimental Instruments and Software). Each chamber was equipped with two nose poke ports positioned 9 cm above the chamber floor. The left port served as the active nose poke, and the right port served as the inactive nose poke. Responses at the active nose poke triggered drug infusion or reward delivery paired with a 5-s light-tone conditioned cue. Responses at the inactive nose poke resulted in no programmed consequences. Active nose poke responses, inactive nose poke responses, and infusion numbers were automatically recorded.

Rats were implanted with an intravenous catheter secured to the skull and connected via flexible tubing to an infusion pump. The tubing was enclosed within a metal spring and connected to a swivel mounted above the chamber, allowing free movement during behavioral testing.

### Self-administration

Cocaine SA training was performed during the dark phase. Rats underwent daily 3-h training sessions, divided into three 1-h sessions separated by 5-min intervals. A fixed-ratio 1 (FR1) reinforcement schedule was used. At the start of each 1-h session, the house light was illuminated. A single active nose poke resulted in immediate intravenous cocaine infusion (0.75 mg/kg/infusion), simultaneous extinction of the house light, presentation of a 5-s auditory cue, and initiation of a 40-s timeout period during which responses had no programmed consequences. The maximum number of cocaine infusions was capped at 20 per hour ^64,65^.

To dissociate pharmacological drug effects from operant drug-seeking behavior, yoked-cocaine design was implemented. Whenever a cocaine SA rat received a cocaine infusion, yoked rats simultaneously received an identical infusion independent of their own behavior. Nose poke responses in yoked rats did not trigger drug delivery. All other experimental parameters were identical to those used for cocaine SA rats.

For heroin SA, each active nose poke triggered an intravenous heroin infusion (0.05 mg/kg per infusion). All other training parameters and behavioral contingencies were identical to those used in the cocaine SA paradigm.

For fentanyl SA, each active nose poke resulted in fentanyl infusion (0.0032 mg/kg per infusion) ^68^. All other training parameters and behavioral contingencies were identical to those used in the cocaine SA paradigm.

In the saline SA group, procedures were identical to those described above, except that cocaine, heroin, or fentanyl was replaced with an equivalent volume of sterile saline ^69^.

For sucrose SA, active nose poke triggered delivery of a sucrose solution via a circular receptacle located between the two nose poke ports. The maximum number of sucrose deliveries was limited to 40 per hour.

Rats in the high-fat food diet group were maintained on either a high-fat or standard chow diet for 42 consecutive days, followed by self-administration training identical to that used for the sucrose SA group ^70,71^.

### Extinction training

During the extinction phase, active nose poke no longer resulted in drug or sucrose infusion and was paired only with a 5-s light-tone cue, under an FR1 schedule. All other training parameters were identical to those used during self-administration. Extinction was considered successful when the number of active nose poke decreased to ≤20% of baseline (mean responses over the last three days of SA) for at least two consecutive days ^72,73^.

### Cue-induced reinstatement

Rats received the assigned pharmacological treatment or vehicle injection before the test. During the 1-h test, all conditions were identical to training except that drug was no longer available. Active nose poke triggered only a 5-s light-tone cue under a FR1 schedule with a 40-s timeout period, but did not result in drug infusion ^74,75^.

### Fiber photometry

Real-time monitoring of acetylcholine (ACh) dynamics during cue-induced reinstatement was performed using a fiber photometry system combined with an ACh fluorescent probe, as previously described ^67,76^. A 488 nm excitation light was delivered through a dichroic mirror and focused by an objective lens, coupled to an optical converter, and connected via an external patch cable to the implanted fiber in the rat’s head. Fluorescence signals were filtered, collected by a photomultiplier tube, and recorded with data acquisition software. Signal analysis was conducted in MATLAB.

On the day prior to testing, patch cables were connected to the implanted ferrule and rats were allowed to habituate in the self-administration chamber for 30 min. On the test day, the system was pre-warmed for 15 min. The fiber tip was adjusted to 30 µW to minimize photobleaching, and the software gain was set to 2.5 V at baseline. Baseline recordings (∼15 s) were obtained with only the patch cable connected. Subsequently, cables were attached to the ferrule and rats were recorded in the home cage for 10 min before being placed in the self-administration chamber for 1 h of test recording. After the session, rats were returned to the home cage, and an additional 15 s of recording was obtained before ending the experiment.

During analysis, each active nose poke during the response period was automatically marked as an event (0 s), and the area under the curve (AUC) of the fluorescence signal from 0-2 s following each event was quantified as an indicator of ACh release.

### Slice preparation

Rats were deeply anesthetized with isoflurane before transcardial perfusion with 100 mL ice-cold choline-based slicing solution containing (in mM): 110 choline chloride, 2.5 KCl, 1.25 NaH_2_PO_4_, 25 NaHCO_3_, 0.5 CaCl_2_, 7 MgCl_2_, 25 glucose, 1 sodium ascorbate, and 3.1 sodium pyruvate (pH 7.4, 305-315 mOsm, bubbled with 95% O_2_/5% CO_2_). Brains were rapidly removed and immersed in ice-cold oxygenated slicing solution. Coronal brain slices containing the nucleus accumbens were prepared at 300 µm thickness using a vibratome (Leica VT1200 S, Germany) ^77^. Slices were incubated in oxygenated slicing solution at 33-37 °C for 10-20 min and then transferred to artificial cerebrospinal fluid (aCSF) containing (in mM): 125 NaCl, 2.5 KCl, 1.25 NaH_2_PO_4_, 25 NaHCO_3_, 2.0 CaCl_2_, 2.0 MgCl_2_, and 10 glucose (pH 7.4, 305-315 mOsm, bubbled with 95% O_2_/5% CO_2_). Slices were allowed to recover at room temperature for at least 30 min before recording.

### Cell-attached patch and whole-cell recordings

Slices were transferred to a submerged recording chamber and continuously perfused with oxygenated aCSF at 32-33 °C at a flow rate of 1-2 mL/min. Neurons in the nucleus accumbens (NAc) were visualized using an IR-DIC microscope (Olympus BX-51WI) equipped with an IR-1000 camera (Dage-MTI). Patch pipettes were pulled from thin-walled borosilicate glass capillaries (BF150-110-10, Sutter Instruments) and had an open-tip resistance of 4-6 MΩ when filled with the appropriate internal solution. Recordings were obtained in either cell-attached or whole-cell patch-clamp configuration^78^ using a

MultiClamp 700B amplifier (Molecular Devices). Data were acquired with pClamp 10.6 software, filtered at 2 kHz, and sampled at 33 kHz with an Axon Digidata 1440A digitizer (Molecular Devices). For whole-cell recordings, data acquisition was initiated after approximately 10 min of stabilization following break-in. Recordings with unstable access resistance or changes in series resistance greater than 20% were excluded from analysis.

### CIN rhythmic firing and *I*_h_ measurement

CINs were targeted in the NAc core of the ventral striatum. CINs were identified based on their large somata, which were substantially larger than neighboring neurons, characteristic multipolar or spearhead-like morphology, lack of dendritic spines, spontaneous rhythmic firing, and prominent hyperpolarization-activated current, *I*_h_ ^14,37,79^.

Spontaneous rhythmic firing of CINs was recorded in gap-free mode using the cell-attached voltage-clamp configuration with the command potential set to 0 mV. This configuration was used to monitor spontaneous action potential firing while minimizing perturbation of the intracellular environment. *I*_h_ was measured using whole-cell voltage-clamp recordings ^80^. Patch pipettes were filled with a potassium-based internal solution containing (in mM): 118 KMeSO_4_, 15 KCl, 2 MgCl_2_, 0.2 EGTA, 10 HEPES, 4 Na_2_ATP, 0.3 Tris-GTP, and 14 Tris-phosphocreatine, adjusted to pH 7.25 with KOH and to 295-305 mOsm. Cells were held at -50 mV and subjected to hyperpolarizing voltage steps from -50 to -120 mV in 10-mV increments. Steady-state currents were measured at each voltage step to generate HCN channel activation curves. In some recordings, HCN channels were maximally activated by stepping the membrane potential from -50 to -130 mV, followed by return steps to test potentials in 10-mV increments. Tail currents were measured to construct current-voltage relationships and estimate relative conductance. Activation kinetics were obtained by fitting the *I*_h_ activation phase with a monoexponential function ^79^.

### AMPA/NMDA ratio recordings

AMPA/NMDA ratios were measured using whole-cell voltage-clamp recordings ^38,39^. Medium spiny neurons (MSNs) in the NAc were targeted based on their medium-sized somata and characteristic morphology. Patch pipettes were filled with a cesium-based internal solution containing (in mM): 122 CsCl, 1 CaCl_2_, 5 MgCl_2_, 10 EGTA, 10 HEPES, 4 Na_2_ATP, 0.3 Tris3-GTP, and 14 Tris2-phosphocreatine, adjusted to pH 7.25 with CsOH and to 295-305 mOsm. A bipolar stimulating electrode was placed approximately 300-350 µm dorsal to the recorded MSNs in the NAc core. During recordings, slices were perfused with aCSF containing 10 µM bicuculline and 2 µM CGP55845 to block GABA_A_ and GABA_B_ receptor-mediated inhibitory transmission, respectively. Evoked excitatory postsynaptic currents (EPSCs) were elicited every 20 s using brief bipolar electrical stimulation. Stimulation intensity was adjusted at -70 mV to evoke stable EPSCs at approximately half of the maximal response amplitude. Cells were then gradually voltage-clamped to +40 mV and allowed to stabilize for 5 min before ten consecutive evoked EPSCs were recorded and averaged to obtain *I*_total_. The perfusion solution was then switched to aCSF containing 10 µM bicuculline, 2 µM CGP55845, and 50 µM D-AP5 to block NMDA receptors. After at least 2 min of stabilization, ten additional evoked EPSCs were recorded and averaged to obtain *I*_AMPA_. The NMDA receptor-mediated component was calculated as *I*_total_-*I*_AMPA_, and the AMPA/NMDA ratio was calculated as:

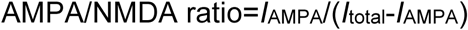

where *I*_total_ represents the averaged evoked current recorded at +40 mV before D-AP5 application, and *I*_AMPA_ represents the averaged evoked current at +40 mV after NMDA receptor blockade.

### Optogenetic manipulation

Optogenetics allows precise control of specific neurons in the brain of freely moving animals using light-sensitive proteins ^81^. In this experiment, ChAT-Cre rats were injected with optogenetic activation (AAV-DIO-ChR2) or inhibition (AAV-DIO-NpHR) viruses, and optical fibers were implanted. When exposed to 473 nm blue light, ChR2 channels open, allowing the influx of cations, including Na^+^ and Ca^2+^, thereby depolarizing and activating neurons. With 580 nm yellow light, NpHR pumps Cl^-^ into neurons, resulting in membrane hyperpolarization and neuronal inhibition.

The day before testing, rats were habituated in a self-administration chamber for 30 min, with jumper cables connected to their head implants. On the test day, equipment was preheated for 15 min. After a 5-min exploration period, rats were placed in the self-administration chamber for behavioral testing. Blue or yellow light was applied to modulate CINs, and behavioral changes were recorded. Stimulation parameters were set at 10 Hz frequency, 10 mW power, 5 ms pulse width, and 1-h duration. The number of active nose pokes was used as the measure of relapse.

### Western blotting

The specimen head of the cryostat was set to -21 °C, and the chamber temperature was set to -24 °C. Rat brains were embedded in optimal cutting temperature compound and equilibrated in the cryostat for 30 min. Coronal sections were then cut at a thickness of 20 µm. When sections corresponding to the target region were reached, bilateral NAc tissue was collected using a sampling needle inserted vertically to a depth of 1 mm.

Collected tissues were lysed in RIPA buffer supplemented with phosphatase and protease inhibitors at a ratio of 100:1:2. A total of 130 µL lysis buffer was added to each sample, followed by thorough homogenization and incubation on ice for 30 min. Lysates were centrifuged, protein concentrations were determined, and samples were normalized with lysis buffer and loading buffer. Samples were heated at 100 °C for 5 min and stored at -80 °C until analysis ^72^.

Equal amounts of protein were separated by SDS-PAGE at 100 V and transferred onto PVDF membranes at a constant current of 250 mA for 2.5 h in ice-cold transfer buffer. PVDF membranes were activated in methanol for 15 s and equilibrated in transfer buffer prior to transfer. After transfer, membranes were blocked at 4 °C for 1 h and incubated with primary antibodies (1:500) overnight at 4 °C. Membranes were washed with 1× TBST and incubated with secondary antibodies (1:2000) for 1 h at room temperature. Protein bands were visualized using enhanced chemiluminescence.

Membranes were stripped and re-incubated when necessary. Band intensities were quantified using Quantity One software, background-subtracted, and normalized to corresponding loading controls for subsequent analysis.

### BaseScope in situ hybridization

To determine the cell-type-specific expression of phospho-eIF2α in dopamine receptor-defined neurons, RNA probe sets targeting Drd1 or Drd2 mRNA were used to identify D1- or D2-expressing neurons, respectively (Cat No. 895071-C1 and 315641-C2, Advanced Cell Diagnostics). BaseScope in situ hybridization was performed according to the manufacturer’s instructions provided by Advanced Cell Diagnostics. Briefly, sections were washed three times in PBS (5 min each) and treated with protease for 10 min at room temperature. After rinsing, sections were hybridized with target-specific probe sets against Drd1 or Drd2 mRNA (separate sections) for 2 h at 40 °C using the HybEZ™ oven. Signal amplification was carried out through sequential incubation with preamplifier, amplifier, and fluorescently labeled probes according to the BaseScope protocol.

Following in situ hybridization, sections were processed for immunofluorescence staining. Sections were incubated with a primary antibody against phospho-EIF2S1 (Ser51) overnight at 4 °C, followed by incubation with appropriate fluorescent secondary antibodies. Nuclei were counterstained with DAPI.

Fluorescence images were acquired using a fluorescence microscope (Olympus, Tokyo, Japan) equipped with a 20× objective lens. Drd1- or Drd2-positive neurons were identified by red fluorescent puncta, and phospho-eIF2α immunoreactivity was visualized in green. Quantification was performed by counting fluorescent puncta and measuring phospho-eIF2α signal intensity within identified neurons. Drd1 and Drd2 signals were analyzed in separate sections.

### Immunofluorescence

Rats were anesthetized with isoflurane and placed in a supine position. A midline incision was made at the xiphoid process and extended bilaterally along the costal margins to open the skin, abdominal cavity, and diaphragm. The rib cage was cut bilaterally to fully expose the heart. A perfusion needle was inserted into the aorta, and PBS was perfused. The right atrium was incised to allow blood drainage. When the liver appeared pale and no further blood outflow was observed, perfusion was switched to 4% PFA until the limbs and neck became rigid.

Animals were decapitated, and brains were rapidly removed and fixed in 4% PFA for 12 h, followed by cryoprotection in 30% sucrose. Brains were then rapidly frozen by immersion in n-hexane precooled in a dry ice-ethanol bath for 16 s, wrapped in aluminum foil, and stored at -80 °C. For sectioning, the specimen head of the cryostat was set to - 21 °C and the chamber temperature to -24 °C. Brain tissues were equilibrated in the cryostat for 30 min, embedded in OCT compound, and sectioned at a thickness of 20 µm. Based on a rat brain stereotaxic atlas, sections containing the target brain region were collected.

Six to eight sections were selected and washed three times in 1× PBS (5 min each), followed by blocking at 37 °C for 1 h. Sections were then incubated with primary antibodies diluted 1:500 at 4 °C for 24 h with gentle agitation. After three PBS washes, sections were incubated with secondary antibodies diluted 1:1000 for 3 h at room temperature. Antibody details are provided in the Key Resources Table. After final PBS washes, sections were mounted onto glass slides, dried at 37 °C, rinsed twice with deionized water, air-dried, and coverslipped with DAPI. Images were acquired using a VS120 virtual slide microscope with a 20× objective.

### Image acquisition and processing

Fluorescence images were acquired using an Olympus fluorescence microscope (Tokyo, Japan) with a 20× objective to detect ChAT-positive neurons. For Extended Data Fig. 2 and Extended Data Fig. 3, images were captured on a Nikon AXR high-resolution confocal system at 400× magnification, acquiring Z-stack images along the vertical axis. All cell counting was performed by an experimenter blinded to group assignments. For each animal, at least three sections were analyzed using FIJI (ImageJ, https://imagej.nih.gov/).

Neuronal somata were selected as regions of interest (ROIs) using the Magic Wand tool in ImageJ, excluding nuclei to avoid non-specific signal. For each ROI, the corrected total cell fluorescence (CTCF) was calculated as: CTCF = Integrated Density - (ROI × Mean background fluorescence) ^34^. Background fluorescence was measured from randomly selected regions in three separate sections, and the mean value was used as the mean background fluorescence for CTCF calculation. Fluorescence measurements were performed on all selected cells within each experimental group. For group comparisons, the average CTCF of the control group was set as the reference value, and CTCF values of individual cells, including those in the control group, were normalized to this reference. Normalized values were used for statistical analyses among groups, with detailed statistical methods provided in the Supplementary Information. An investigator blinded to the experimental conditions performed the image analyses.

### Single-cell RNA sequencing and data analysis

ChAT/tdTomato rats were subjected to either cocaine SA or saline SA training. ChAT-positive neurons were identified by red fluorescence and individually collected via whole-cell patch-clamp ^82,83^. Collected neurons were immediately placed in lysis buffer and lysed using a thermocycler. First-strand cDNA was synthesized using the SMART-Seq II amplification system, followed by PCR pre-amplification. Amplified cDNA products were quality-checked, and qualified samples were used for library preparation via a transposase-based method, followed by library quality assessment. PCR products were denatured into single-stranded DNA, circularized to generate single-stranded circular DNA, and uncircularized linear DNA molecules were digested. Single-stranded circular DNA molecules were then amplified by rolling-circle replication to generate DNA nanoballs (DNBs) containing multiple copies of each template. DNBs were loaded onto high-density DNA nanoarrays and sequenced using combinatorial probe-anchor synthesis (cPAS) technology.

All raw sequencing data generated in this study have been deposited in the NCBI Sequence Read Archive (SRA) under BioProject number PRJNA1392453.

### Data analysis

Raw sequencing reads were processed using SOAPnuke (v2.2.1) to remove low-quality sequences ^84^. Reads were excluded if they: (1) contained adapter sequences, (2) contained more than 5% unknown nucleotides (N), or (3) had more than 20% of bases with a quality score below 15. The remaining high-quality reads were defined as clean data. Clean data were subsequently analyzed, visualized, and mined using the Dr. Tom multi-omics data analysis platform (https://biosys.bgi.com).

### Variant detection

Clean reads were aligned to the reference genome using HISAT2 (v2.2.1) ^85^. Gene fusion events were detected using EricScript (v0.5.5). Alternative splicing events and differential splicing were analyzed using rMATS (v4.1.2) ^86^.

### Differential gene expression analysis

Clean reads were mapped to the reference gene set using Bowtie2 (v2.4.5). Gene expression levels were quantified using RSEM (v1.3.1). Heatmaps depicting gene expression across samples were generated using pheatmap (v1.0.12). Differentially expressed genes (DEGs) were identified using DESeq2 (v1.34.0) ^87^, using a significance threshold of *P* ≤ 0.05 ^88,89^.

### GO and KEGG enrichment analysis

To explore the functional relevance of DEGs associated with phenotypic changes, Gene Ontology (GO; http://www.geneontology.org/) and KEGG pathway (https://www.kegg.jp/) enrichment analyses were performed using a hypergeometric test implemented in Phyper. Terms with Q value ≤ 0.05 were considered significantly enriched among candidate genes.

### Single-cell RT-qPCR

Ex vivo living brain slices were prepared from ChAT/tdTomato rats, and ChAT-positive neurons were aspirated using borosilicate glass pipettes. Target cells were identified based on tdTomato fluorescence. Collected single cells were processed according to the manufacturer’s instructions using the Single Cell Sequence Specific Amplification Kit (Vazyme, Nanjing, China). Real-time reverse transcription quantitative PCR (RT-qPCR) was performed using ChamQ Universal SYBR qPCR Master Mix (Vazyme) on a Mastercycler® nexus gradient thermal cycler (Eppendorf, Germany). Relative mRNA expression levels were normalized to GAPDH as an internal reference and calculated using the 2^-ΔΔCt^ method in GraphPad Prism. All experiments were performed with three independent biological replicates ^90,91^.

### Statistical analyses

Statistical analyses were performed with GraphPad Prism 8. Data were analyzed using paired or unpaired t tests, one-way ANOVA, two-way ANOVA, or two-way repeated-measures ANOVA, followed by Dunnett’s or Sidak’s multiple-comparison tests as specified in Supplementary Tables 1-19. Data are presented as mean ± s.e.m. Exact sample sizes, test statistics, degrees of freedom and P values are provided in Supplementary Tables 1-19. Significance was defined as \**P* < 0.05, \*\**P* < 0.01, \*\*\**P* < 0.001 and \*\*\*\**P* < 0.0001. Rats were randomly assigned to each group. Behavioral experiments were replicated in at least two batches of animals.

## Acknowledgements

We thank Ji Hu, Peter Kalivas and all members from the Xue laboratory for discussions and comments.

## Author contributions

Conceptualization: Y.X.X. and Z.H.

Behavioral, functional imaging, and electrophysiological experiments: S.H.H., J.L.Z., Y.L., X.Y.L., Z.H.S., Z.B.D., Z.H.L. and Y.T.

Formal analysis: S.H.H., J.L.Z., X.Y.Z., H.Y.L., and Y.L.

Visualization: S.H.H., J.L.Z., and Y.L.

Funding acquisition: Y.X.X. and Z.H.

Project administration and supervision: Y.X.X., Z.H., L.L., and J.S.

Writing - original draft: S.H.H., J.L.Z., and Y.L.

Writing - review and editing: S.H.H., J.L.Z., Y.L., Y.C.H., Y.X.L., H.Y.L., Y.X.X., L.L., and J.S.

## Data availability

All raw sequencing data generated in this study have been deposited in the NCBI Sequence Read Archive (SRA) under BioProject accession number PRJNA1392453. All other data reported in this paper are available from the corresponding authors upon reasonable request. No custom code was generated or used for the central analyses in this study. Commercially available chemicals, pharmacological reagents, and viral vectors used in this study are described in the Supplementary Information. Information on custom viral constructs, shRNA sequences, and other key reagents is available from the corresponding authors upon reasonable request.

Extended Data Figs. 1-14

Supplementary Tables 1-19. Detailed statistics associated with Figs. 1-6 and Extended Data Figs. 1-13.

---

**Extended Data Fig. 1.**
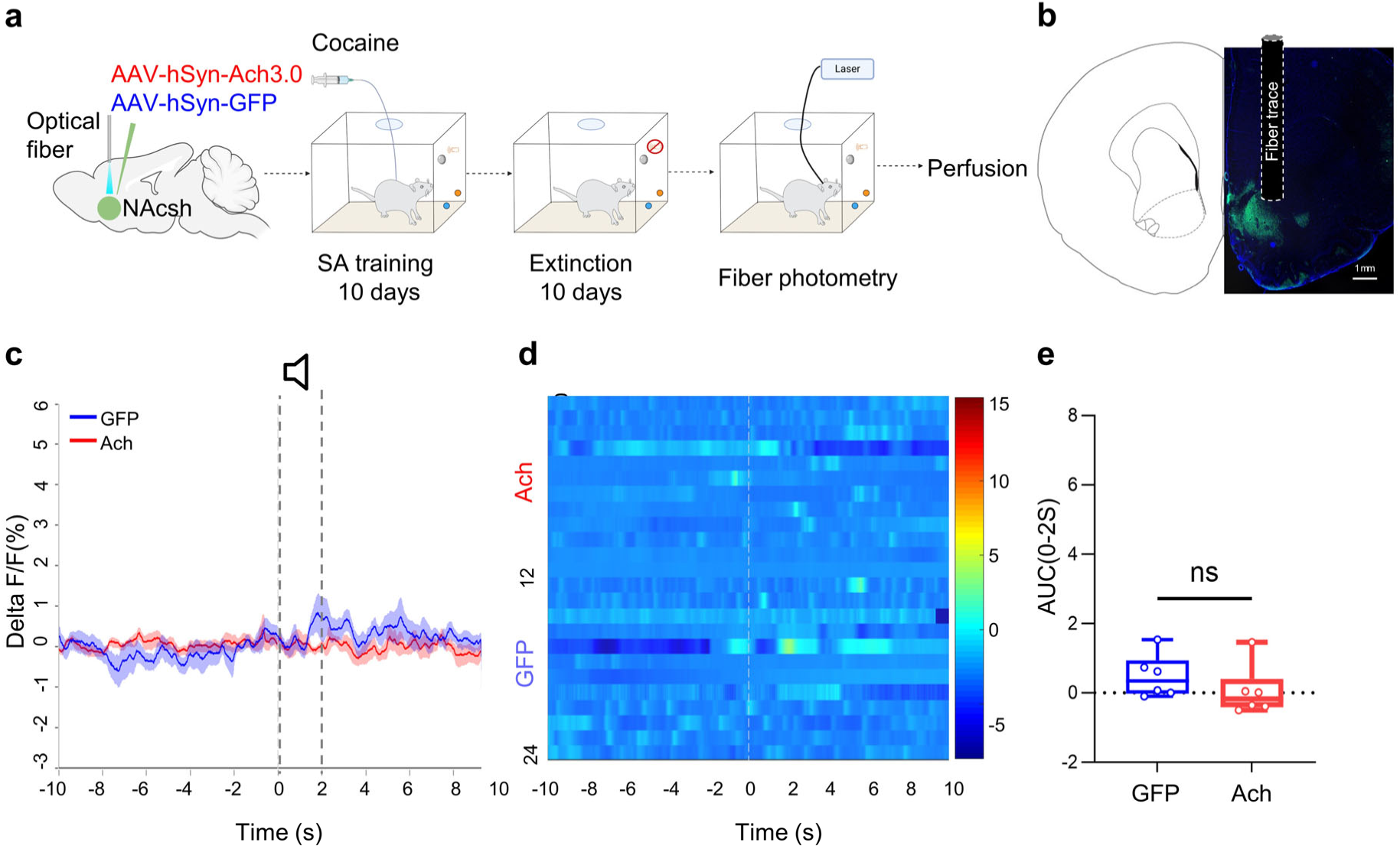
Cocaine-paired cues do not evoke significant ACh transients in the NAcSh. **(a)** Schematic of *in vivo* recording of acetylcholine release from cholinergic interneurons in the NAcSh. **(b)** Illustration of fiber photometry implant sites and viral expression. Scale bar = 1 mm. **(c and d)** Fiber photometry recordings of ACh3.0/GFP signals **(c)** and corresponding heatmap **(d)**, with Time 0 indicating drug-associated cue presentation. **(e)** AUC of z-scored ACh3.0 signals in NAcSh CINs during 0-2 s after cue onset. n = 6 rats for GFP and n = 7 rats for ACh. Data are presented as mean ± s.e.m. For statistical details, see Supplementary Table 7.

**Extended Data Fig. 2.**
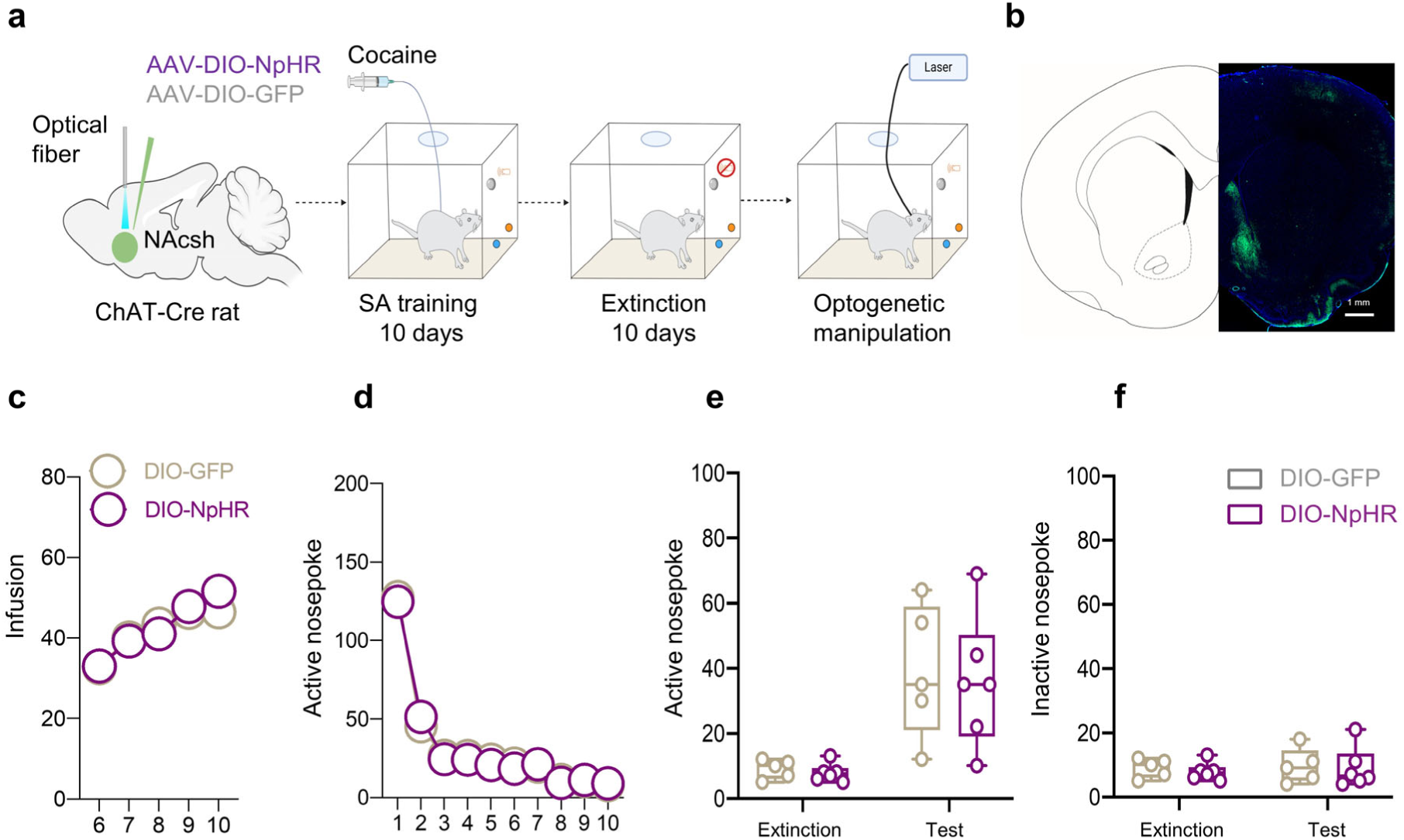
Optogenetic inhibition of NAcSh CINs has no effect on cue-induced cocaine seeking. **(a)** Timeline of optogenetic inhibition of NAcSh CINs during cue-induced reinstatement. **(b)** Viral injection sites and fiber implant locations. Scale bar = 1 mm. **(c)** Number of cocaine infusions during the acquisition phase. **(d)** Number of active nose poke responses during the extinction sessions. **(e-f)** Number of active and inactive nose poke responses during reinstatement with optogenetic inhibition of CINs. n = 5 rats for DIO-GFP and n = 6 rats for DIO-NpHR. Data are presented as mean ± s.e.m. For statistical details, see Supplementary Table 8.

**Extended Data Fig. 3.**
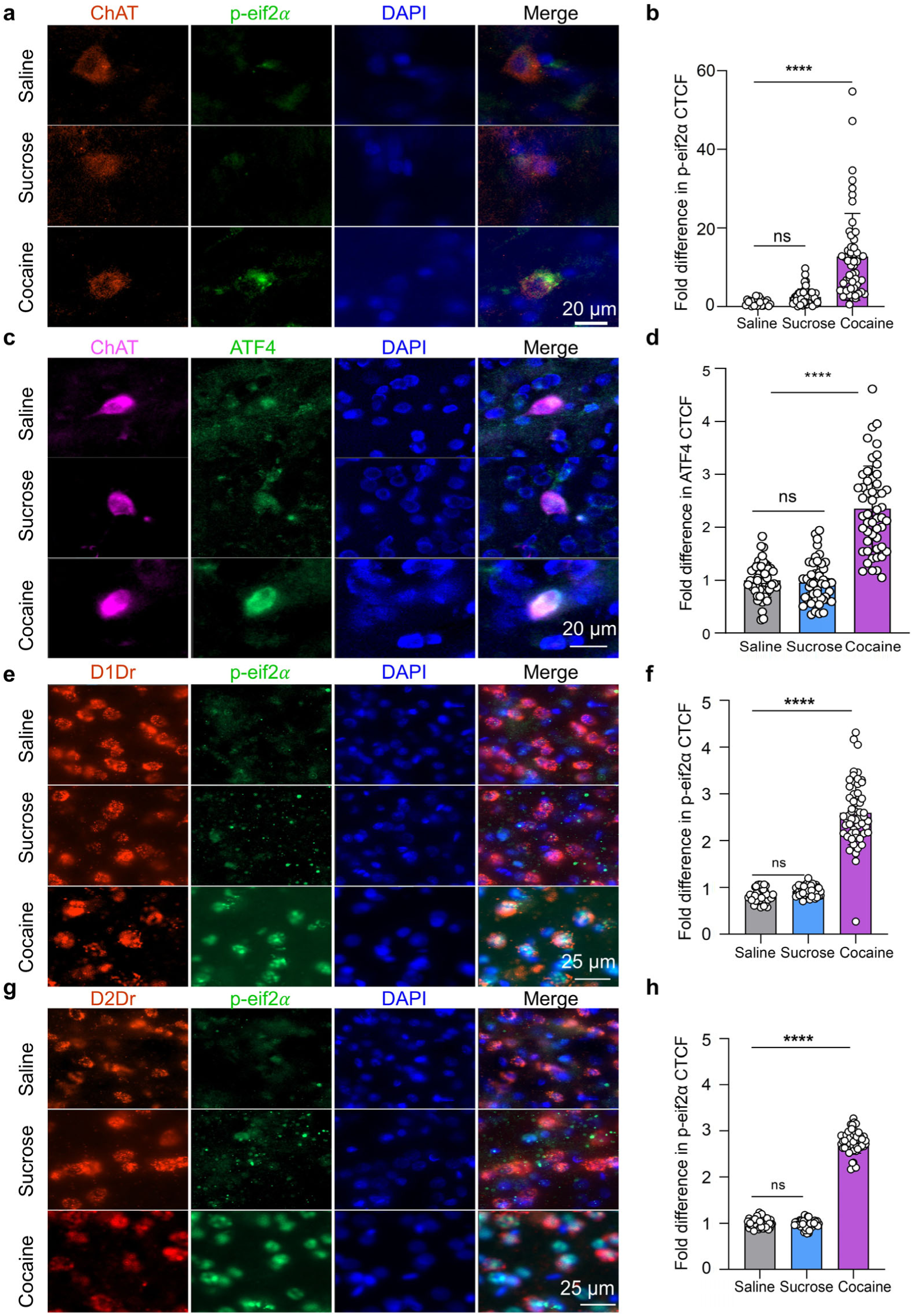
Cocaine SA, but not sucrose SA, increases ISR markers in CINs and D1- and D2-expressing neurons. **(a)** Representative photomicrographs of immunofluorescence staining of ChAT (red), phospho-eIF2α (green), and DAPI (blue) in the NAc from the saline, sucrose and cocaine groups, scale bars = 20 µm. **(b)** Quantification of p-eIF2α CTCF from (a). n = 22, 45 and 55 cells from 3 rats for saline, sucrose and cocaine groups, respectively. Data are presented as mean ± s.e.m. \*\*\*\**P* < 0.0001 by one-way ANOVA with Dunnett’s multiple-comparison test. For statistical details, see Supplementary Table 9. **(c)** Representative photomicrographs of immunofluorescence staining of ChAT (purple), ATF4 (green), and DAPI (blue) in the NAc from the saline, sucrose and cocaine groups, scale bars = 20 µm. **(d)** Quantification of ATF4 CTCF in ChAT-positive CINs from (c). n = 51, 47 and 52 cells from 3 rats for saline, sucrose and cocaine groups, respectively. Data are presented as mean ± s.e.m. \*\*\*\**P* < 0.0001 by one-way ANOVA with Dunnett’s multiple-comparison test. For statistical details, see Supplementary Table 9. **(e)** Representative photomicrographs of immunofluorescence staining of D1 neurons (red), phospho-eIF2α (green), and DAPI (blue) in the NAc from the saline, sucrose and cocaine groups, scale bars = 25 µm. D1Dr, dopamine receptor D1. **(f)** Quantification of p-eIF2α CTCF from (e). n = 40, 43 and 57 cells from 3 rats for saline, sucrose and cocaine groups, respectively. Data are presented as mean ± s.e.m. \*\*\*\**P* < 0.0001 by one-way ANOVA with Dunnett’s multiple-comparison test. For statistical details, see Supplementary Table 9. **(g)** Representative photomicrographs of immunofluorescence staining of D2 neurons (red), phospho-eIF2α (green), and DAPI (blue) in the NAc from the saline, sucrose and cocaine groups, scale bars = 25 µm. D2Dr, dopamine receptor D2. **(h)** Quantification of p-eIF2α CTCF from (g). n = 46, 49 and 53 cells from 3 rats for saline, sucrose and cocaine groups, respectively. Data are presented as mean ± s.e.m. \*\*\*\**P* < 0.0001 by one-way ANOVA with Dunnett’s multiple-comparison test. For statistical details, see Supplementary Table 9.

**Extended Data Fig. 4.**
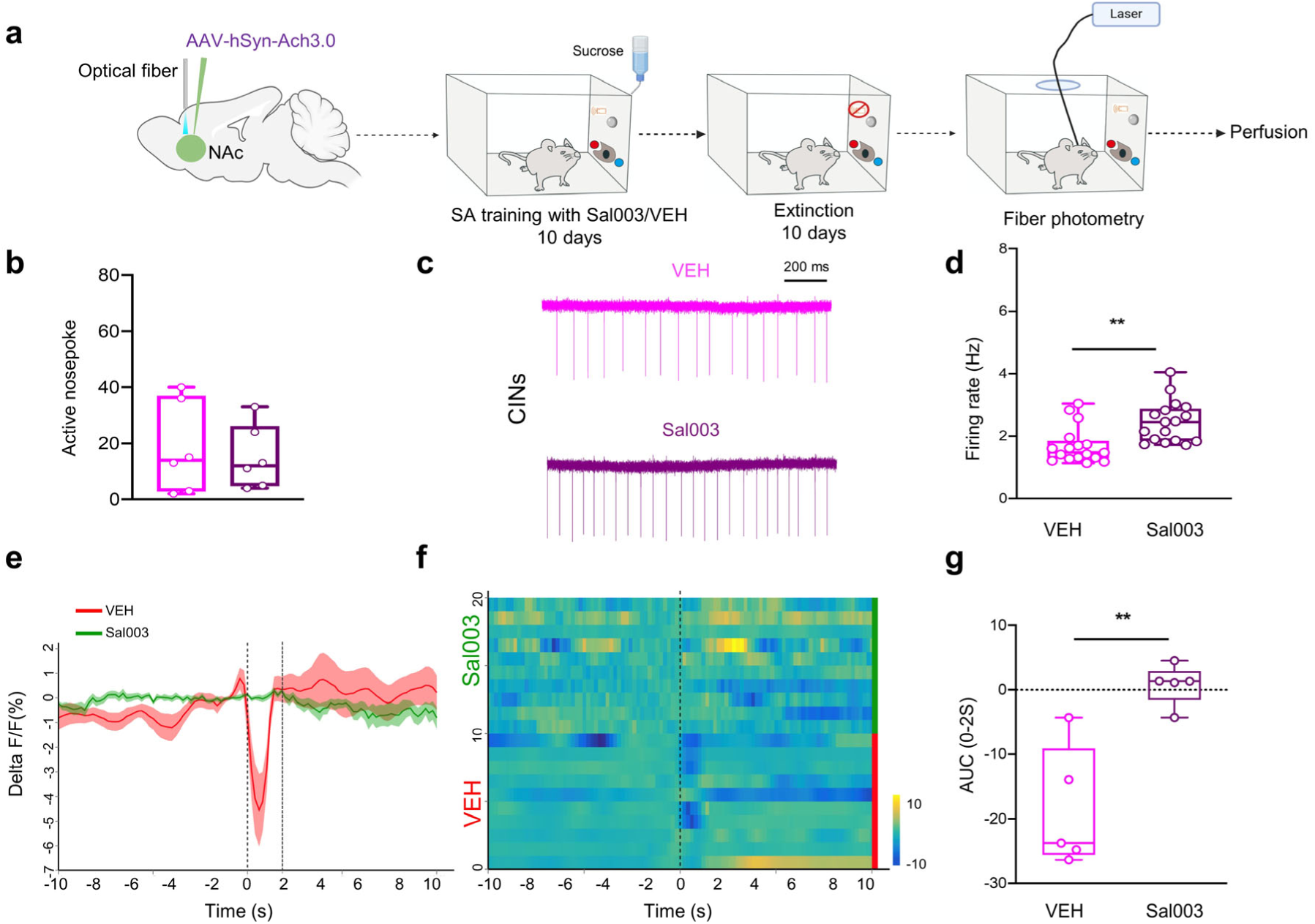
Elevating ISR signaling during sucrose training alters CIN excitability and cue-evoked ACh dynamics without enhancing sucrose seeking. **(a)** Experimental timeline showing Sal003 or vehicle treatment during sucrose SA training, followed by extinction and cue-induced reinstatement. **(b)** Number of active nose poke responses during cue-induced reinstatement. n = 6 rats for each group. **(c and d)** Representative CIN firing traces (c) and quantification of firing rates (d). n = 17 cells from 3 rats per group. **(e-g)** ACh3.0 photometric responses aligned to sucrose-cue onset: temporal dynamics of ΔF/F signals (e), heatmaps of trial-aligned signals (f), and AUC values over the 0-2 s post-cue interval (g). n = 5 rats for each group. Data are presented as mean ± s.e.m. \**P* < 0.05, \*\**P* < 0.01, \*\*\**P* < 0.001, \*\*\*\**P* < 0.0001 by unpaired *t* test (b, d and g). For statistical details, see Supplementary Table 10.

**Extended Data Fig. 5.**
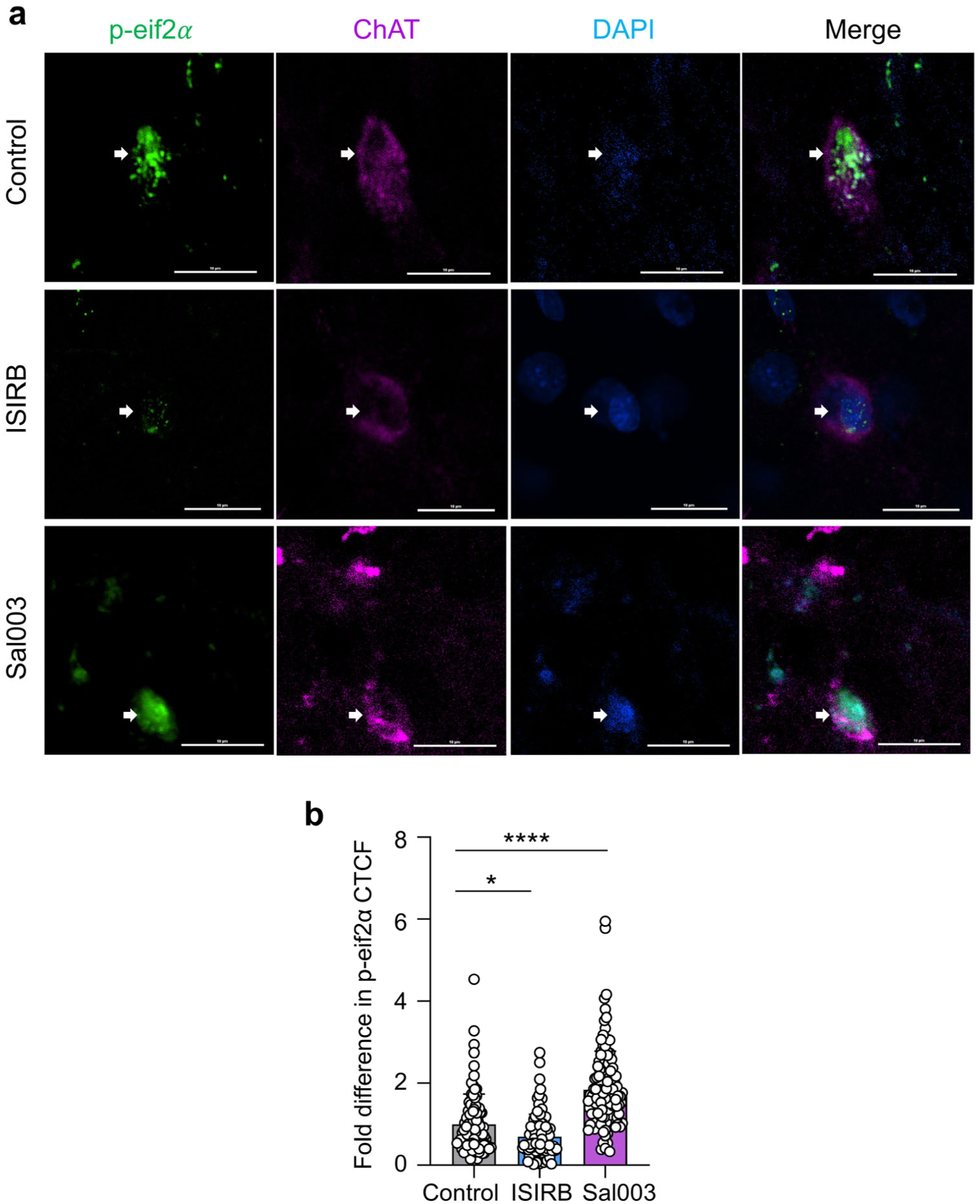
Pharmacological manipulation of ISR signaling bidirectionally regulates p-eIF2α levels in CINs. **(a)** Representative photomicrographs of immunofluorescence staining of phospho-eIF2α (green), ChAT (purple) and DAPI (blue) in the NAc from the control, ISRIB and Sal003 treatment groups, scale bars = 10 µm. **(b)** Quantification of p-eIF2α CTCF from (a). n = 93, 93 and 128 cells from 3 rats for control, ISRIB and Sal003 groups, respectively. Data are presented as mean ± s.e.m. \**P* < 0.05, \*\*\*\**P* < 0.0001 by one-way ANOVA with Dunnett’s multiple-comparison test. For statistical details, see Supplementary Table 11.

**Extended Data Fig. 6.**
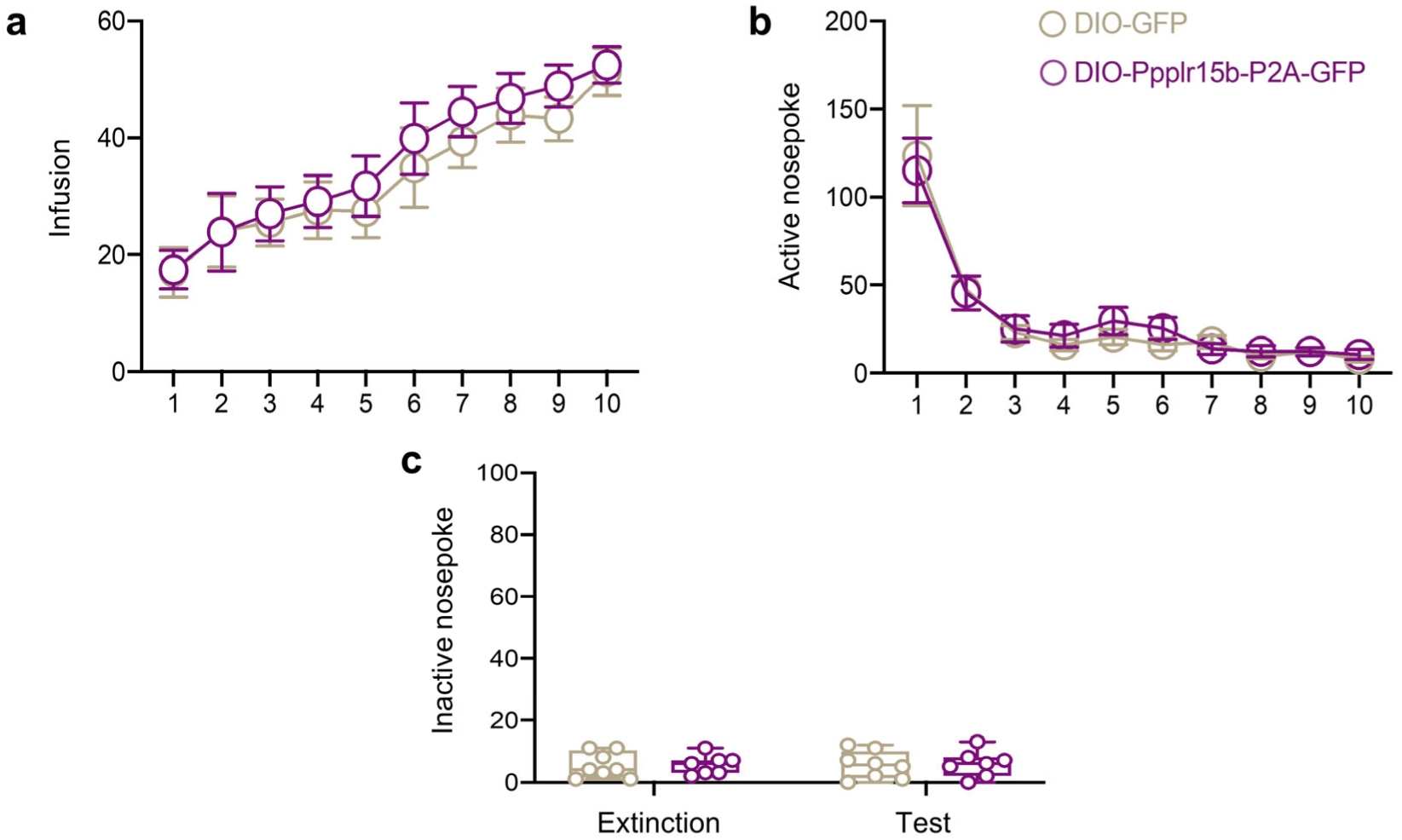
CIN-specific ISR suppression does not alter cocaine acquisition, extinction, or inactive responding. **(a)** Number of cocaine infusions during the acquisition phase after CIN-specific Ppp1r15b expression. **(b)** Number of active nose poke responses during extinction. **(c)** Number of inactive nose poke responses on the last day of extinction and during cue-induced reinstatement. n = 8 rats for DIO-GFP and n = 7 rats for DIO-Ppp1r15b-P2A-GFP. Data are presented as mean ± s.e.m. For statistical details, see Supplementary Table 12.

**Extended Data Fig. 7.**
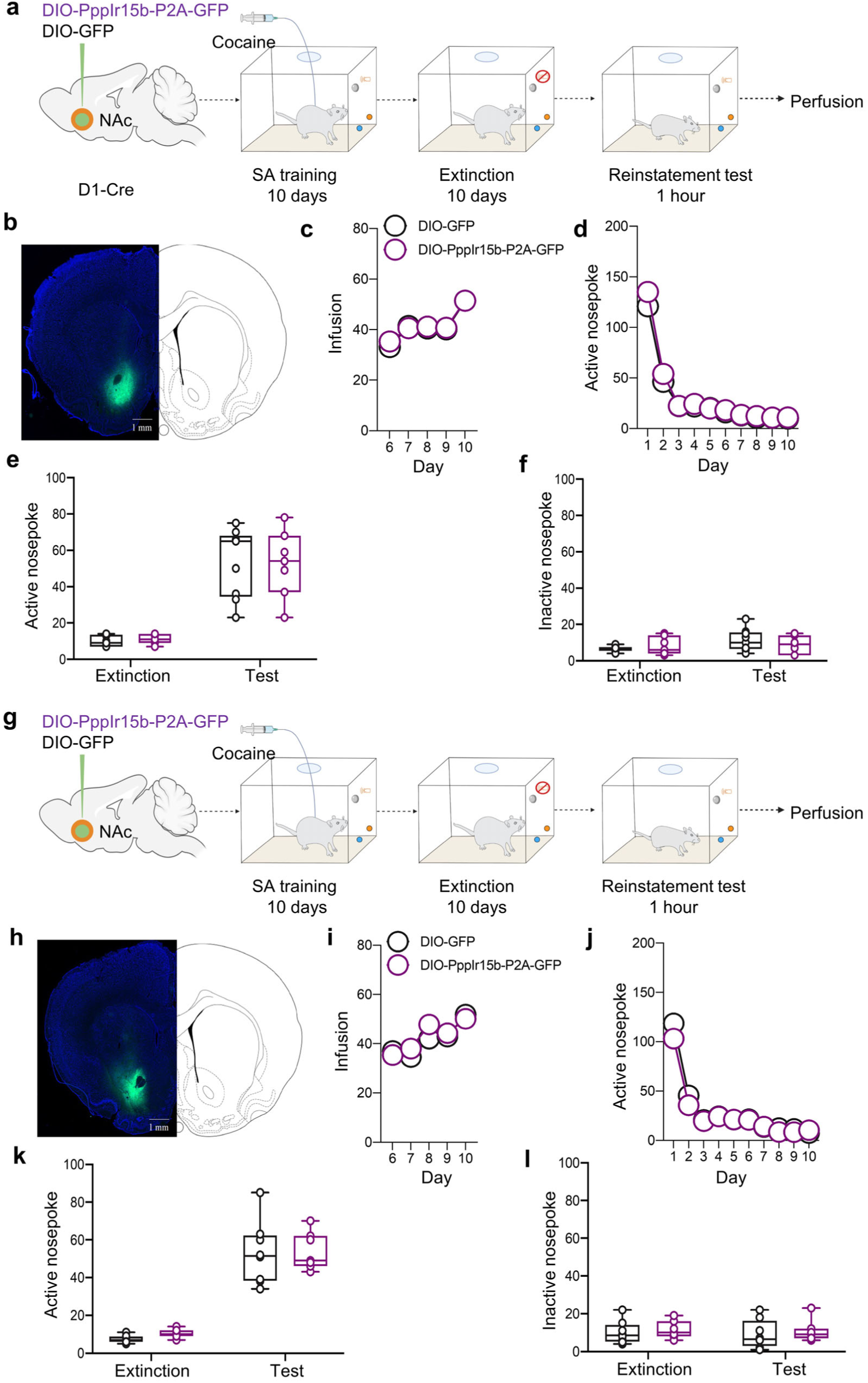
Genetic suppression of ISR signaling in NAc D1R- or D2R-expressing neurons does not affect cue-induced cocaine seeking. **(a-f)** D1R-neuron-specific ISR suppression: experimental timeline (a), validation of viral expression in the NAc (b), cocaine infusions during acquisition (c), active nose poke responses during extinction (d), active responses on the last extinction day and reinstatemen*t* test (e), and inactive responses on the last extinction day and reinstatemen*t* test (f). n = 9 rats for DIO-GFP and n = 7 rats for DIO-Ppp1r15b-P2A-GFP in (e,f). **(g-l)** D2R-neuron-specific ISR suppression: experimental timeline (g), Validation of viral expression in the NAc (h), cocaine infusions during acquisition (i), active nose poke responses during extinction (j), active responses on the last extinction day and reinstatemen*t* test (k), and inactive responses on the last extinction day and reinstatemen*t* test (l). n = 8 rats for DIO-GFP and n = 7 rats for DIO-Ppp1r15b-P2A-GFP in (k,l). Data are presented as mean ± s.e.m. For statistical details, see Supplementary Table 13.

**Extended Data Fig. 8.**
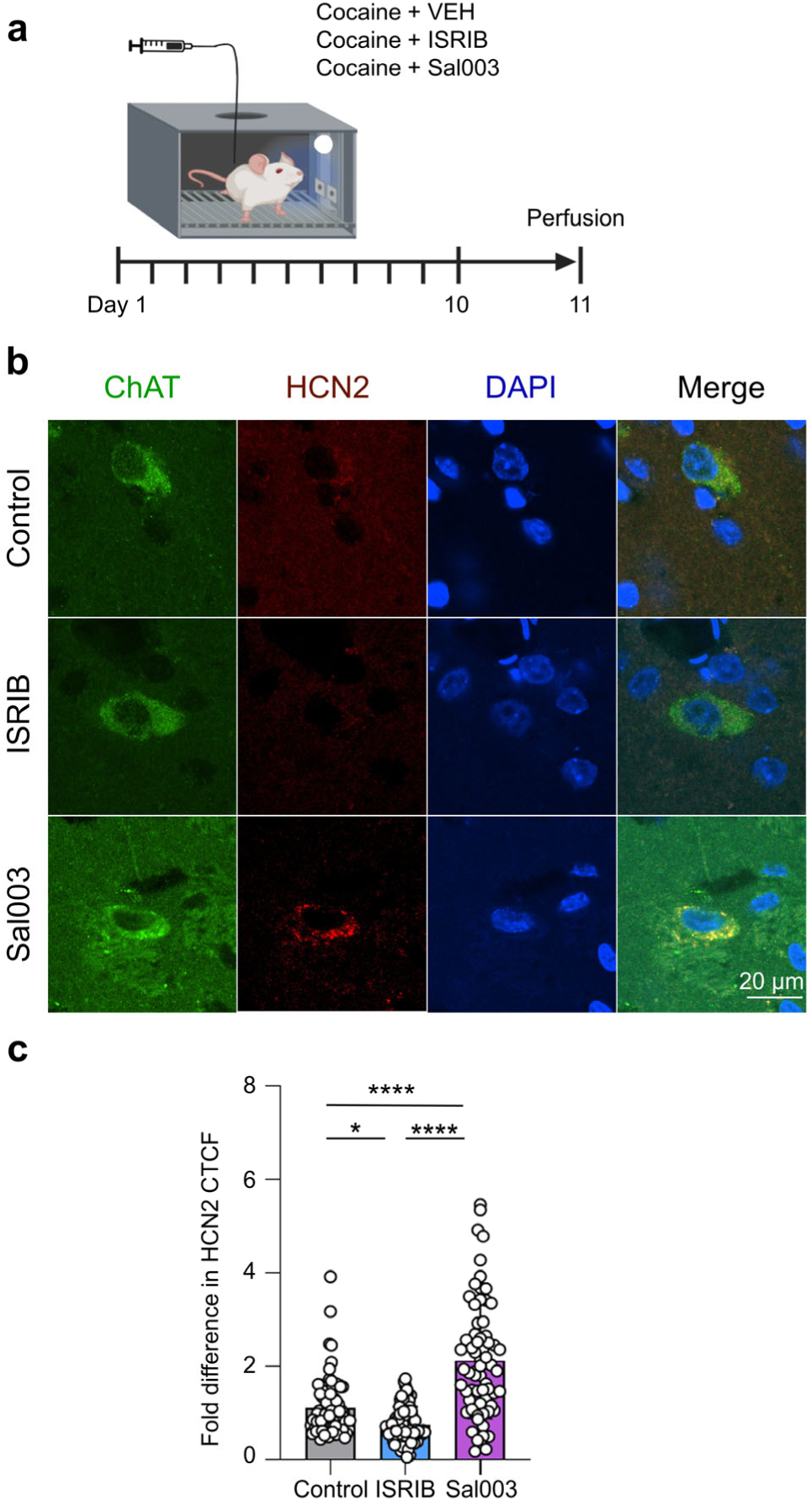
Pharmacological manipulation of ISR signaling bidirectionally regulates HCN2 protein expression in CINs. **(a)** Schematic timeline for pharmacological manipulation of eIF2α phosphorylation during SA training. **(b)** Representative immunofluorescence images showing HCN2 (red), ChAT (green), and DAPI (blue) staining in the NAc of control, ISRIB-, and Sal003-treated groups, scale bar = 20 µm. n = 83, 86 and 69 cells from 3 rats for control, ISRIB and Sal003 groups, respectively. **(c)** Quantification of HCN2 CTCF from **(b)**. Data are presented as mean ± s.e.m. \**P* < 0.05, \*\*\*\**P* < 0.0001 by one-way ANOVA with Dunnett’s multiple-comparison test. For statistical details, see Supplementary Table 14.

**Extended Data Fig. 9.**
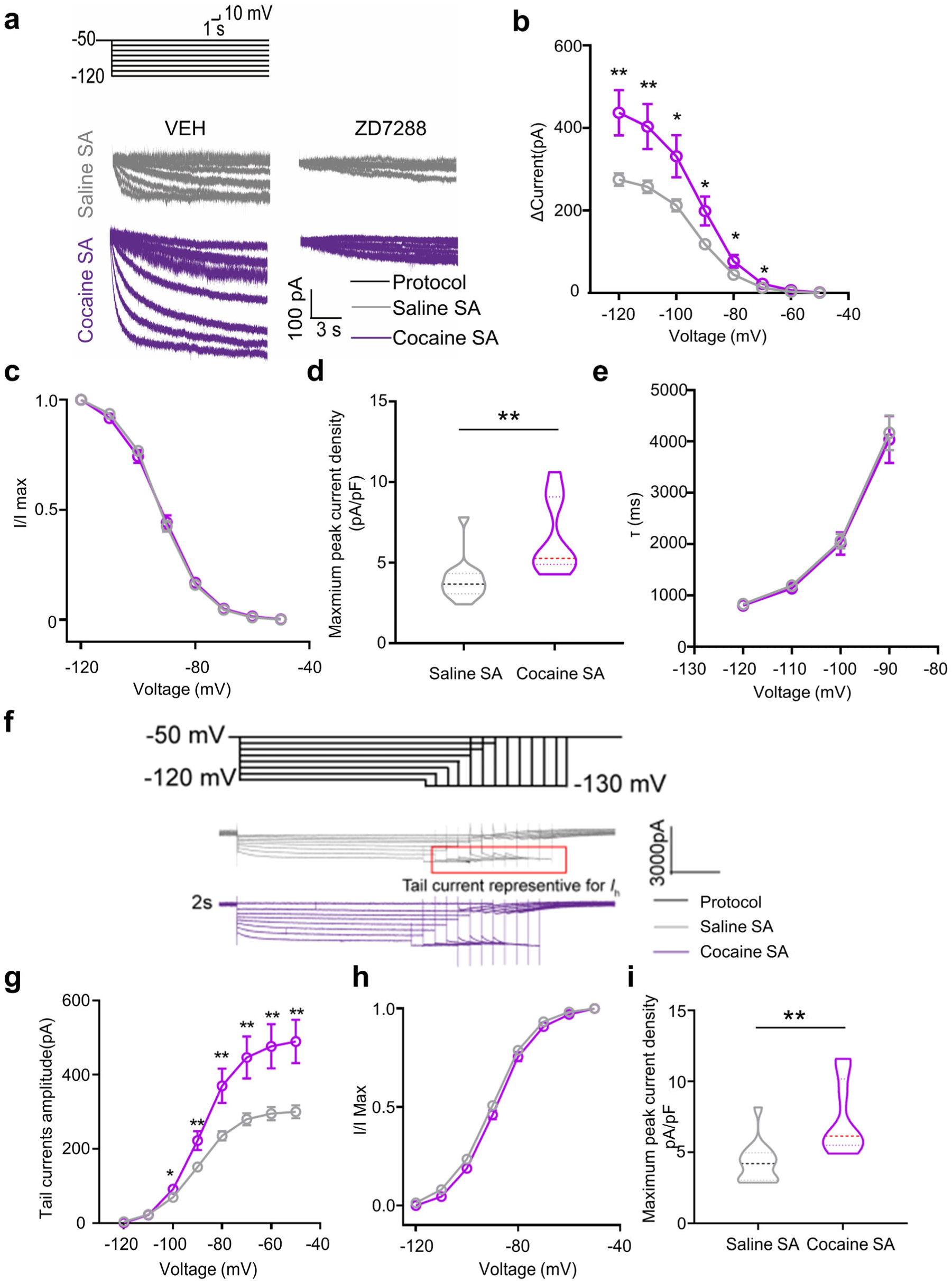
Cocaine training increases maximal and available functional *I*_h_ in CINs. **(a-e)** Maximal functional *I*_h_: voltage-step protocol and representative HCN current traces before and during 15 µM ZD7288 in saline- and cocaine-trained rats (a), steady-state activation curves (b), normalized activation curves (c), maximal peak current density (d), and activation time constants across voltage steps (e). n = 14 cells for saline SA and n = 9 cells for cocaine SA. **(f-i)** Available functional *I*_h_: tail-current protocol and representative traces before and during 15 µM ZD7288 (f), steady-state tail-current activation curves (g), normalized activation curves (h), and maximal peak current density (i). n = 14 cells for saline SA and n = 9 cells for cocaine SA. Data are presented as mean ± s.e.m. \**P* < 0.05, \*\**P* < 0.01 by unpaired *t* test. For statistical details, see Supplementary Table 15.

**Extended Data Fig. 10.**
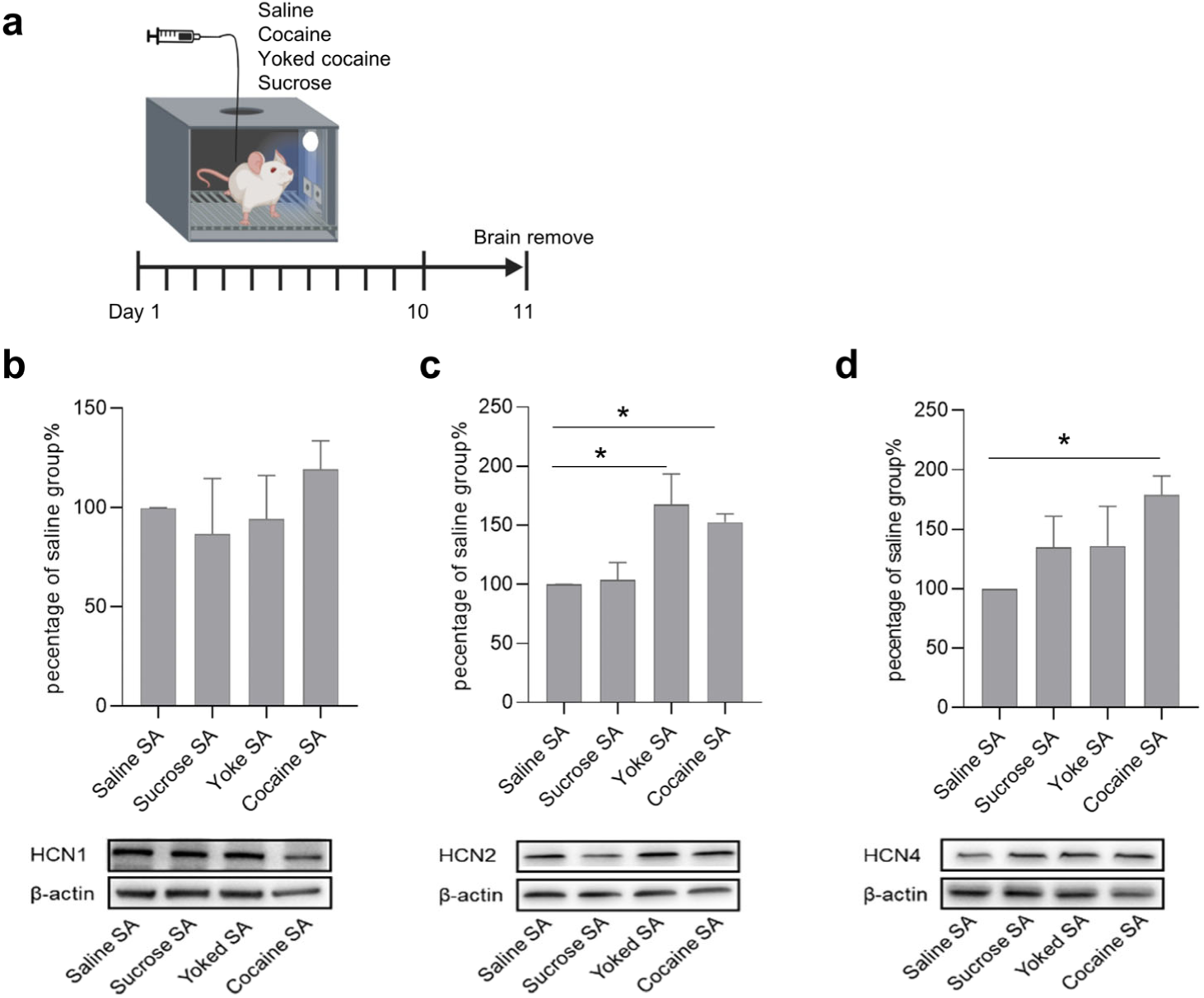
Cocaine training increases HCN2 and HCN4 expression in the NAc. **(a)** Timeline of the different drugs training experiments. **(b-d)** Expression levels of HCN channel subtypes (HCN1, HCN2 and HCN4) and representative immunoblot images. For HCN1 and HCN2 (b,c), n = 5, 5, 5 and 7 rats for saline SA, sucrose SA, yoked cocaine and cocaine SA groups, respectively; for HCN4 (d), n = 6, 6, 6 and 8 rats, respectively. Data are presented as mean ± s.e.m. \**P* < 0.05 by one-way ANOVA with Dunnett’s multiple-comparison test (b, c and d). For statistical details, see Supplementary Table 16.

**Extended Data Fig. 11.**
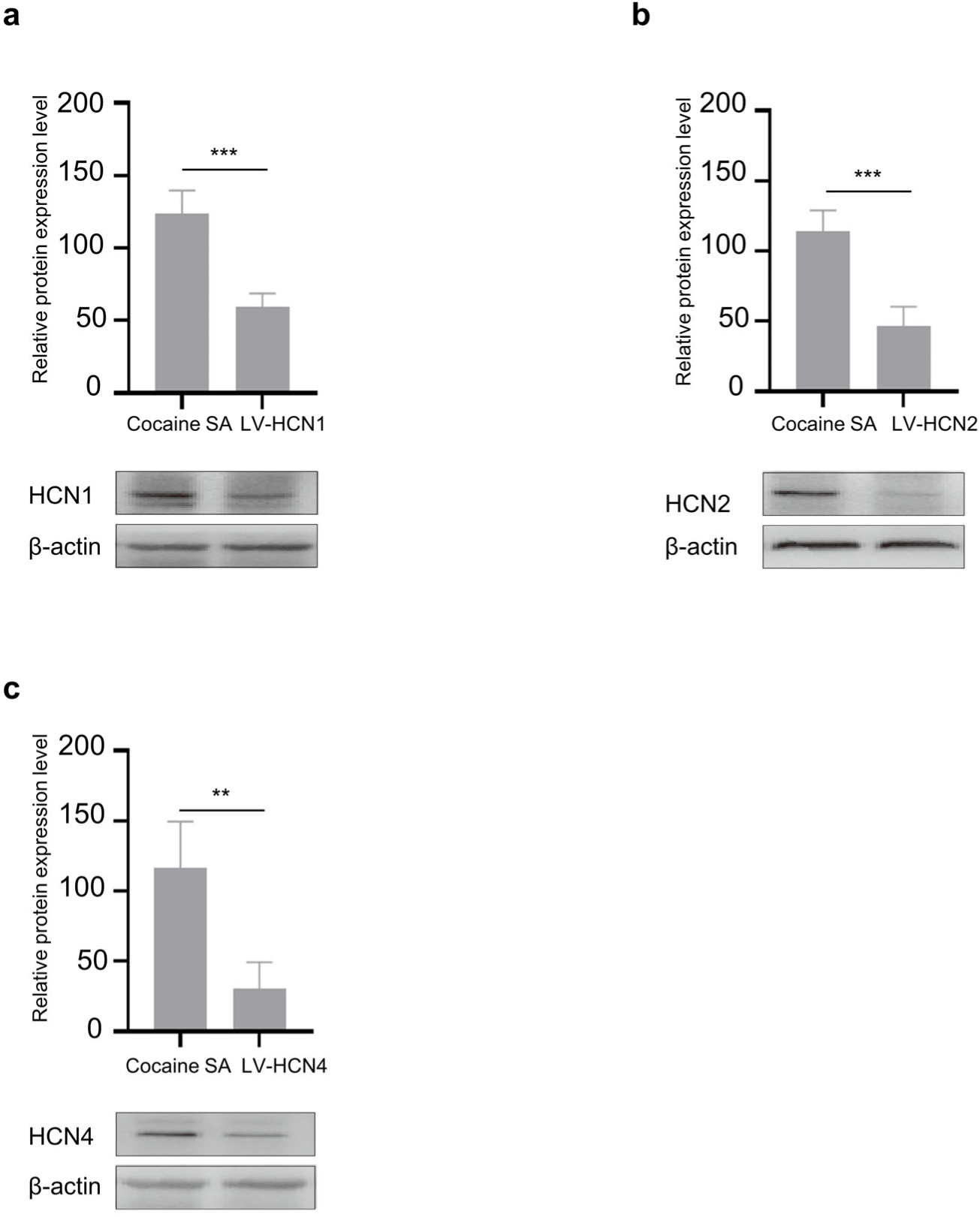
Validation of lentiviral-mediated knockdown efficiency for different HCN isoforms. **(a)** Expression levels of HCN1 and representative immunoblot images. n = 3 rats for Cocaine SA and n = 6 rats for LV-HCN1. **(b)** Expression levels of HCN2 and representative immunoblot images. n = 3 rats for Cocaine SA and n = 6 rats for LV-HCN2. **(c)** Expression levels of HCN4 and representative immunoblot images. n = 3 rats for Cocaine SA and n = 4 rats for LV-HCN4. Data are presented as mean ± s.e.m. \*\**P* < 0.01, \*\*\**P* < 0.001. Unpaired *t* test (a, b and c). For statistical details, see Supplementary Table 17.

**Extended Data Fig. 12.**
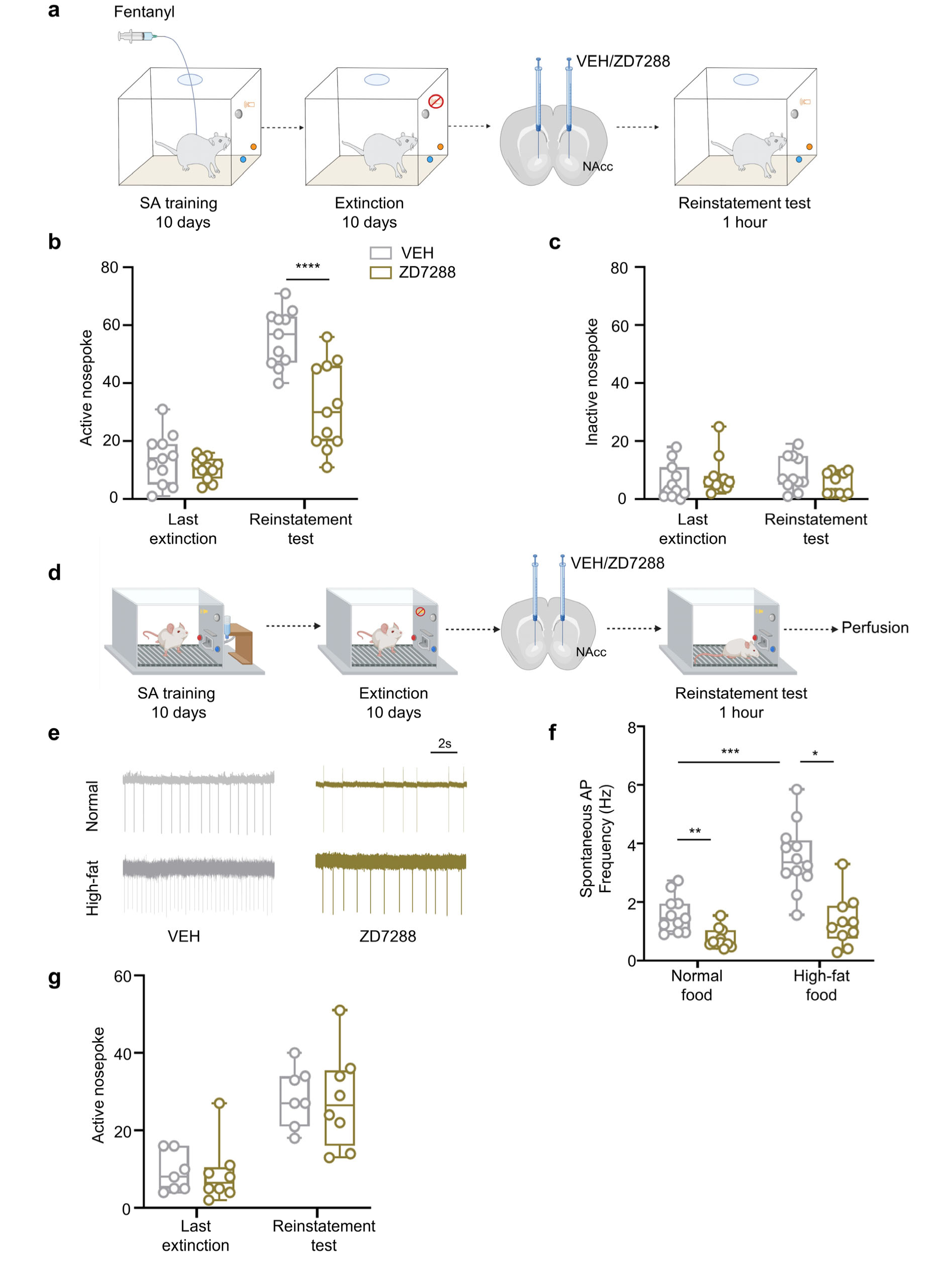
NAcC HCN blockade suppresses fentanyl seeking, normalizes high-fat CIN firing, and spares sucrose seeking. **(a)** Schematic timeline for intra-NAcC infusion of ZD7288 in fentanyl SA-trained rats. **(b and c)** Active (b) and inactive (c) nose poke responses during cue-induced fentanyl reinstatement. n = 11 rats for each group. **(d)** Schematic timeline for intra-NAcC infusion of ZD7288 in high-fat-food- and sucrose-trained rats. **(e and f)** Representative spontaneous CIN firing traces from normally fed and high-fat diet-fed rats following VEH or ZD7288 treatment (e) and quantification of firing rates (f). For normal food, n = 11 cells per group; for high-fat food, n = 12 VEH and n = 11 ZD7288 cells. **(g)** Active nose poke responses in sucrose SA-trained rats on the last extinction session and during cue-induced reinstatement. n = 7 rats for VEH and n = 8 rats for ZD7288. Data are presented as mean ± s.e.m. \**P* < 0.05, \*\**P* < 0.01, \*\*\**P* < 0.001, \*\*\*\**P* < 0.0001. Statistical tests are as specified in Supplementary Table 18.

**Extended Data Fig. 13.**
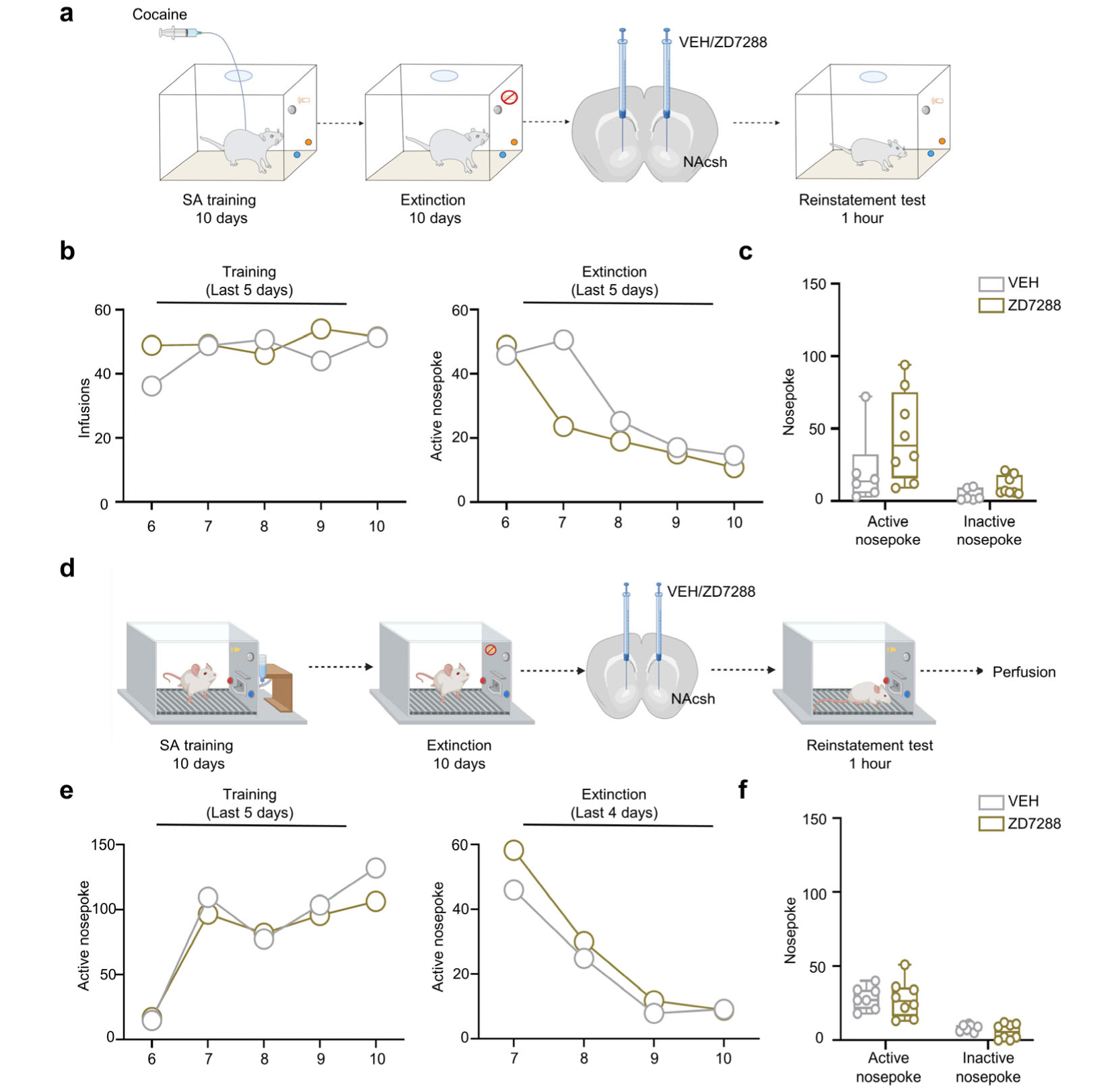
NAcSh HCN blockade does not affect cue-induced cocaine or sucrose seeking. **(a)** Schematic timeline for intra-NAc shell infusion of ZD7288 in cocaine SA-trained rats. **(b)** Number of cocaine infusions during the last five days of acquisition session (left) and number of active nose poke responses during the last five days of extinction sessions (right). **(c)** The number of active and inactive nose poke responses during the cue-induced reinstatemen*t* test. n = 6 rats for VEH and n = 8 rats for ZD7288. **(d)** Schematic timeline for intra-NAc shell infusion of ZD7288 in sucrose SA-trained rats. **(e)** Number of active nose poke responses during the last five days of sucrose acquisition (left) and during the last four days of extinction (right). **(f)** The number of active and inactive nose poke responses during the cue-induced reinstatemen*t* test. n = 7 rats for VEH and n = 8 rats for ZD7288. Data are presented as mean ± s.e.m. ns: *P* > 0.05 by two-way ANOVA with Sidak’s multiple-comparison test (c and f). For statistical details, see Supplementary Table 19.

**Extended Data Fig. 14.**
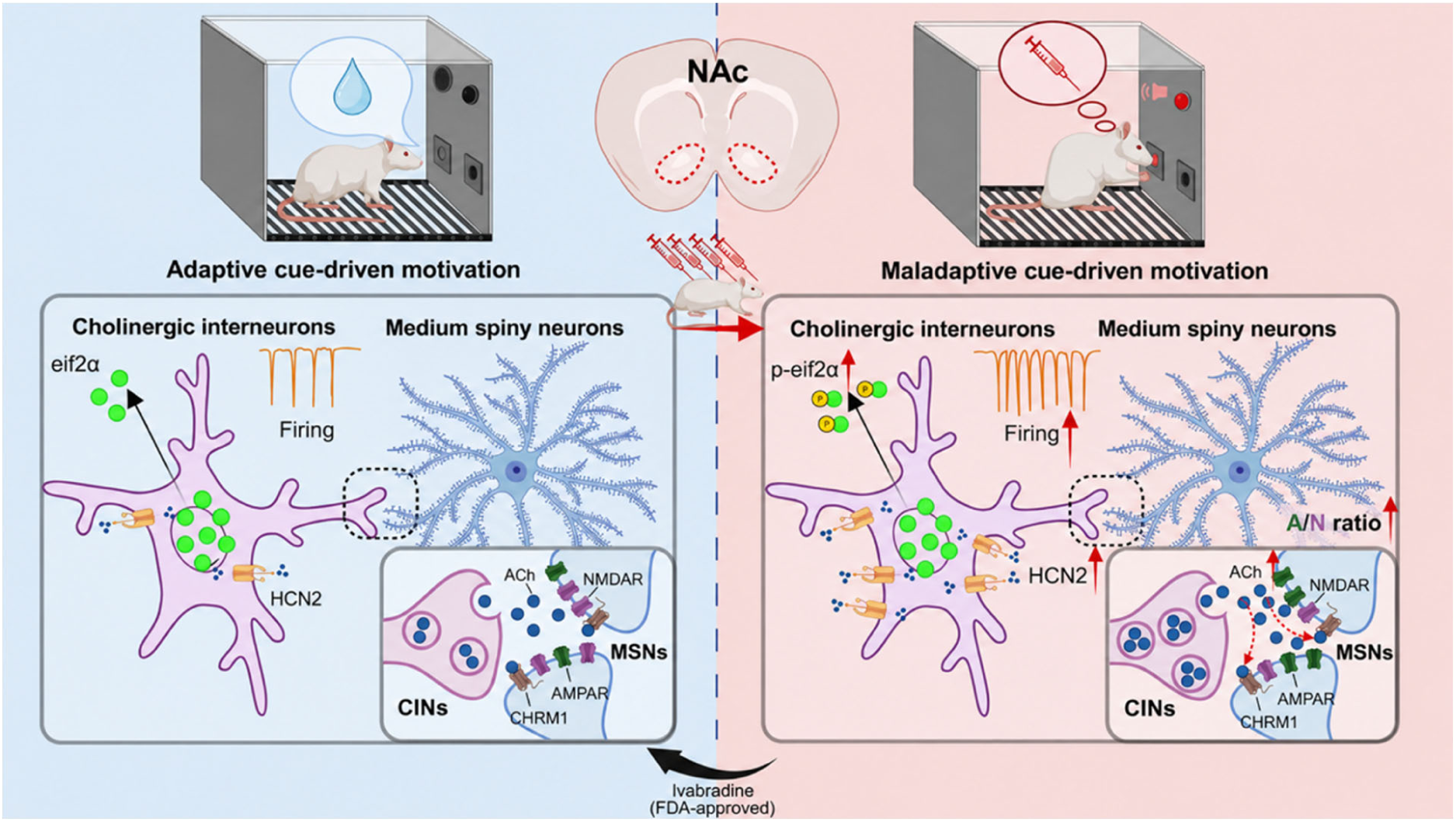
Proposed model of the accumbal CIN ISR-HCN2 axis in maladaptive cue-driven motivation. Under natural-reward conditions, basal ISR and HCN2 signaling support physiological pacemaker activity of nucleus accumbens CINs and limited cue-evoked ACh signaling. Following voluntary drug-taking experience, increased eIF2α phosphorylation upregulates HCN2 and enhances HCN-dependent CIN excitability. This persistent high-firing state promotes cue-evoked ACh release and cholinergic modulation of MSNs, potentially through muscarinic receptors, together with increased AMPA/NMDA current ratios. These cellular and synaptic adaptations strengthen maladaptive cue-driven motivation and facilitate cue-induced drug seeking. CIN-specific HCN2 knockdown or pharmacological HCN blockade, including systemic ivabradine treatment, attenuates cue-induced seeking. Red arrows indicate increases relative to the natural-reward condition. NAc, nucleus accumbens; ISR, integrated stress response; HCN2, hyperpolarization-activated cyclic nucleotide-gated channel 2; CHRM1, muscarinic acetylcholine receptor M1.

