## Supplementary Tables 1-16 for "An accumbal cholinergic ISR-HCN2 axis sustains maladaptive cue motivation"

**Supplementary Table 1.** Addictive drug and high-fat self-administration increase CIN firing and cue-evoked ACh release in the NAcC.

| Figure | Panel | Sample size | Normality test | Normality test results | Test | Comparison | *F* | *t* | df | *P* | Post hoc test |
| --- | --- | --- | --- | --- | --- | --- | --- | --- | --- | --- | --- |
| Fig. 1 | e | Saline: n = 18;<br>Cocaine: n = 12;<br>Yoked cocaine: n = 20;<br>Sucrose: n = 21;<br>High-fat food: n = 12;<br>Heroin: n = 17 | Shapiro-Wilk test | Pass | One-way ANOVA | Drug | 13.3 | / | 5, 94 | <0.0001 | Saline vs. Cocaine: <0.0001; Saline vs. Yoked cocaine: 0.9998;<br>Saline vs. Sucrose: 0.8049;<br>Saline vs. High-fat food: 0.0016;<br>Saline vs. Heroin: 0.0002 |
| Fig. 1 | j | GFP: n = 6;<br>ACh: n = 5 | Shapiro-Wilk test | Pass | Unpaired t test | Intervention | / | 7.601 | 9 | <0.0001 | / |

**Supplementary Table 2.** NAcC CINs are required for cue-induced cocaine seeking and associated MSN synaptic potentiation.

| Figure | Panel | Sample size | Normality test | Normality test results | Test | Comparison | F | t | df | P | Post hoc test |
| --- | --- | --- | --- | --- | --- | --- | --- | --- | --- | --- | --- |
| Fig. 2 | b | Last extinction: VEH: n = 6;<br>Saporin: n = 7;<br>Reinstatement test: VEH: n = 6;<br>Saporin: n = 7 | Shapiro-Wilk test | Pass | Two-way repeated-measures ANOVA | Time × Treatment | 3.884 | / | 1, 11 | 0.0744 | Last extinction (Vehicle vs. Saporin): 0.9776;<br>Reinstatement test (Vehicle vs. Saporin): 0.03 |
| Fig. 2 | b | Last extinction: VEH: n = 6;<br>Saporin: n = 7;<br>Reinstatement test: VEH: n = 6;<br>Saporin: n = 7 | Shapiro-Wilk test | Pass | Two-way repeated-measures ANOVA | Time | 3.071 | / | 1, 11 | 0.1075 | Last extinction (Vehicle vs. Saporin): 0.9776;<br>Reinstatement test (Vehicle vs. Saporin): 0.03 |
| Fig. 2 | b | Last extinction: VEH: n = 6;<br>Saporin: n = 7;<br>Reinstatement test: VEH: n = 6;<br>Saporin: n = 7 | Shapiro-Wilk test | Pass | Two-way repeated-measures ANOVA | Treatment | 2.883 | / | 1, 11 | 0.1205 | Last extinction (Vehicle vs. Saporin): 0.9776;<br>Reinstatement test (Vehicle vs. Saporin): 0.03 |
| Fig. 2 | c | VEH: n = 6;<br>Saporin: n = 6 | Shapiro-Wilk test | Pass | Unpaired t test | Treatment | / | 8.469 | 10 | <0.0001 | / |
| Fig. 2 | e | VEH: n = 20;<br>Saporin: n = 16 | Shapiro-Wilk test | Pass | Unpaired t test | Treatment | / | 2.225 | 34 | 0.0307 | / |
| Fig. 2 | h | DIO-GFP: n = 7;<br>DIO-NpHR: n = 7 | Shapiro-Wilk test | Pass | Two-way repeated-measures ANOVA | Time × Intervention | 6.877 | / | 1, 12 | 0.0223 | Extinction (DIO-GFP vs. DIO-NpHR): 0.6126;<br>Test (DIO-GFP vs. DIO-NpHR): 0.0136 |
| Fig. 2 | h | DIO-GFP: n = 7;<br>DIO-NpHR: n = 7 | Shapiro-Wilk test | Pass | Two-way repeated-measures ANOVA | Time | 51.6 | / | 1, 12 | <0.0001 | Extinction (DIO-GFP vs. DIO-NpHR): 0.6126;<br>Test (DIO-GFP vs. DIO-NpHR): 0.0136 |
| Fig. 2 | h | DIO-GFP: n = 7;<br>DIO-NpHR: n = 7 | Shapiro-Wilk test | Pass | Two-way repeated-measures ANOVA | Intervention | 2.206 | / | 1, 12 | 0.1633 | Extinction (DIO-GFP vs. DIO-NpHR): 0.6126;<br>Test (DIO-GFP vs. DIO-NpHR): 0.0136 |
| Fig. 2 | i | DIO-GFP: n = 7;<br>DIO-NpHR: n = 7 | Shapiro-Wilk test | Pass | Two-way repeated-measures ANOVA | Time × Intervention | 0.9771 | / | 1, 12 | 0.3424 | Extinction (DIO-GFP vs. DIO-NpHR): 0.6291;<br>Test (DIO-GFP vs. DIO-NpHR): > 0.9999 |
| Fig. 2 | i | DIO-GFP: n = 7;<br>DIO-NpHR: n = 7 | Shapiro-Wilk test | Pass | Two-way repeated-measures ANOVA | Time | 1.237 | / | 1, 12 | 0.2879 | Extinction (DIO-GFP vs. DIO-NpHR): 0.6291;<br>Test (DIO-GFP vs. DIO-NpHR): > 0.9999 |
| Fig. 2 | i | DIO-GFP: n = 7;<br>DIO-NpHR: n = 7 | Shapiro-Wilk test | Pass | Two-way repeated-measures ANOVA | Intervention | 0.2641 | / | 1, 12 | 0.6167 | Extinction (DIO-GFP vs. DIO-NpHR): 0.6291;<br>Test (DIO-GFP vs. DIO-NpHR): > 0.9999 |
| Fig. 2 | j | DIO-GFP: n = 7;<br>DIO-ChR2: n = 6 | Shapiro-Wilk test | Pass | Two-way repeated-measures ANOVA | Time × Intervention | 4.789 | / | 1, 11 | 0.0511 | Extinction (DIO-GFP vs. DIO-ChR2): > 0.9999;<br>Test (DIO-GFP vs. DIO-ChR2): 0.0061 |
| Fig. 2 | j | DIO-GFP: n = 7;<br>DIO-ChR2: n = 6 | Shapiro-Wilk test | Pass | Two-way repeated-measures ANOVA | Time | 62.62 | / | 1, 11 | <0.0001 | Extinction (DIO-GFP vs. DIO-ChR2): > 0.9999;<br>Test (DIO-GFP vs. DIO-ChR2): 0.0061 |
| Fig. 2 | j | DIO-GFP: n = 7;<br>DIO-ChR2: n = 6 | Shapiro-Wilk test | Pass | Two-way repeated-measures ANOVA | Intervention | 6.31 | / | 1, 11 | 0.0289 | Extinction (DIO-GFP vs. DIO-ChR2): > 0.9999;<br>Test (DIO-GFP vs. DIO-ChR2): 0.0061 |
| Fig. 2 | k | DIO-GFP: n = 7;<br>DIO-ChR2: n = 6 | Shapiro-Wilk test | Pass | Two-way repeated-measures ANOVA | Time × Intervention | 0.2522 | / | 1, 11 | 0.6254 | Extinction (DIO-GFP vs. DIO-ChR2): > 0.9999;<br>Test (DIO-GFP vs. DIO-ChR2): 0.4778 |
| Fig. 2 | k | DIO-GFP: n = 7;<br>DIO-ChR2: n = 6 | Shapiro-Wilk test | Pass | Two-way repeated-measures ANOVA | Time | 2.636 | / | 1, 11 | 0.1328 | Extinction (DIO-GFP vs. DIO-ChR2): > 0.9999;<br>Test (DIO-GFP vs. DIO-ChR2): 0.4778 |
| Fig. 2 | k | DIO-GFP: n = 7;<br>DIO-ChR2: n = 6 | Shapiro-Wilk test | Pass | Two-way repeated-measures ANOVA | Intervention | 1.392 | / | 1, 11 | 0.263 | Extinction (DIO-GFP vs. DIO-ChR2): > 0.9999;<br>Test (DIO-GFP vs. DIO-ChR2): 0.4778 |

**Supplementary Table 3.** CIN-specific ISR signaling sustains cocaine-induced hyperexcitability and seeking.

| Figure | Panel | Sample size | Normality test | Normality test results | Test | Comparison | F | t | df | P | Post hoc test |
| --- | --- | --- | --- | --- | --- | --- | --- | --- | --- | --- | --- |
| Fig. 3 | b | VEH: n = 6;<br>ISRIB: n = 6 | Shapiro-Wilk test | Pass | Unpaired t test | Drug | / | 2.682 | 10 | 0.023 | / |
| Fig. 3 | d | VEH: n = 17 from 3 rats;<br>ISRIB: n = 17 from 3 rats. | Shapiro-Wilk test | Pass | Unpaired t test | Drug | / | 8.352 | 32 | < 0.0001 | / |
| Fig. 3 | g | VEH: n = 6;<br>ISRIB: n = 5 | Shapiro-Wilk test | Pass | Unpaired t test | Drug | / | 5.118 | 9 | 0.0006 | / |
| Fig. 3 | j | DIO-GFP: n = 8;<br>DIO-Ppp1r15b-P2A-GFP: n = 7 | Shapiro-Wilk test | Pass | Two-way repeated-measures ANOVA | Time × Intervention | 19.85 | / | 1, 13 | 0.0006 | Extinction (DIO-GFP vs. DIO-Ppp1r15b-P2A-GFP): 0.8945;<br>Test (DIO-GFP vs. DIO-Ppp1r15b-P2A-GFP): <0.0001 |
| Fig. 3 | j | DIO-GFP: n = 8;<br>DIO-Ppp1r15b-P2A-GFP: n = 7 | Shapiro-Wilk test | Pass | Two-way repeated-measures ANOVA | Time | 42.07 | / | 1, 13 | < 0.0001 | Extinction (DIO-GFP vs. DIO-Ppp1r15b-P2A-GFP): 0.8945;<br>Test (DIO-GFP vs. DIO-Ppp1r15b-P2A-GFP): <0.0001 |
| Fig. 3 | j | DIO-GFP: n = 8;<br>DIO-Ppp1r15b-P2A-GFP: n = 7 | Shapiro-Wilk test | Pass | Two-way repeated-measures ANOVA | Intervention | 10.04 | / | 1, 13 | 0.0074 | Extinction (DIO-GFP vs. DIO-Ppp1r15b-P2A-GFP): 0.8945;<br>Test (DIO-GFP vs. DIO-Ppp1r15b-P2A-GFP): <0.0001 |
| Fig. 3 | l | Control: n = 14 from 4 rats;<br>Ppp1r15b: n = 11 from 3 rats. | Shapiro-Wilk test | Pass | Unpaired t test | Intervention | / | / | 23 | < 0.0001 | / |

**Supplementary Table 4.** Cocaine engages an ISR-sensitive HCN program in CINs.

| Figure | Panel | Sample size | Normality test | Normality test results | Test | Comparison | F | t | df | P | Post hoc test |
| --- | --- | --- | --- | --- | --- | --- | --- | --- | --- | --- | --- |
| Fig. 4 | a | Control: n = 19;<br>ZD7288: n = 18;<br>TTAP2: n = 7;<br>Nifedipine: n = 7;<br>Iberritoxin: n = 8;<br>Apamin: n = 8;<br>XE991: n = 7;<br>AP5 + CNQX: n = 7;<br>Bicuculline + CGP55845: n = 7;<br>TTX: n = 5 | Shapiro-Wilk test | Pass | One-way ANOVA | Intervention | 9.793 | / | 9, 83 | <0.0001 | Control vs. ZD7288: <0.0001;<br>Control vs. TTAP2: 0.9979;<br>Control vs. Nifedipine: >0.9999;<br>Control vs. Iberritoxin: 0.9999;<br>Control vs. Apamin: 0.9994;<br>Control vs. XE991: 0.9943;<br>Control vs. AP5 + CNQX: 0.1106;<br>Control vs. Bicuculline + CGP55845: 0.9159;<br>Control vs. TTX: <0.0001 |
| Fig. 4 | c | VEH: n = 18;<br>ZD7288: n = 18 | Shapiro-Wilk test | Pass | Paired t test | Treatment | / | 4.404 | 17 | 0.0004 | / |
| Fig. 4 | e | VEH: n = 12;<br>ZD7288: n = 12 | Shapiro-Wilk test | Pass | Paired t test | Treatment | / | 3.693 | 11 | 0.0035 | / |
| Fig. 4 | f | Saline SA + VEH: n = 18;<br>Cocaine SA + VEH: n = 12;<br>Cocaine SA + ZD7288: n = 12 | Shapiro-Wilk test | Pass | One-way ANOVA | Treatment | 6.892 | / | 2, 39 | 0.0027 | Saline SA + Vehicle vs. Cocaine SA + Vehicle: 0.0037;<br>Cocaine SA + Vehicle vs. Cocaine SA + ZD7288: 0.0120 |
| Fig. 4 | k (left) | HCN2: VEH: n = 9;<br>ISRIB: n = 9 | Shapiro-Wilk test | Pass | Unpaired t test | Treatment | / | 3.817 | 16 | 0.0015 | / |
| Fig. 4 | k (right) | HCN4: VEH: n = 12;<br>ISRIB: n = 12 | Shapiro-Wilk test | Pass | Unpaired t test | Treatment | / | 2.675 | 22 | 0.0138 | / |

**Supplementary Table 5.** HCN2 is the CIN-specific causal effector of cue-induced cocaine seeking.

| Figure | Panel | Sample size | Normality test | Normality test results | Test | Comparison | F | t | df | P | Post hoc test |
| --- | --- | --- | --- | --- | --- | --- | --- | --- | --- | --- | --- |
| Fig. 5 | b | LV-Control, Cocaine SA: n = 9;<br>LV-HCN1, Cocaine SA: n = 5;<br>LV-HCN2, Cocaine SA: n = 12;<br>LV-HCN4, Cocaine SA: n = 7 | Shapiro-Wilk test | Pass | Two-way ANOVA | Intervention × Voltage | 2.279 | / | 21, 232 | 0.0016 | LV-Control, Cocaine SA vs. LV-HCN1, Cocaine SA: 0.963;<br>LV-Control, Cocaine SA vs. LV-HCN2, Cocaine SA: <0.0001;<br>LV-Control, Cocaine SA vs. LV-HCN4, Cocaine SA: 0.0252 |
| Fig. 5 | b | LV-Control, Cocaine SA: n = 9;<br>LV-HCN1, Cocaine SA: n = 5;<br>LV-HCN2, Cocaine SA: n = 12;<br>LV-HCN4, Cocaine SA: n = 7 | Shapiro-Wilk test | Pass | Two-way ANOVA | Intervention | 22.77 | / | 3, 232 | <0.0001 | LV-Control, Cocaine SA vs. LV-HCN1, Cocaine SA: 0.963;<br>LV-Control, Cocaine SA vs. LV-HCN2, Cocaine SA: <0.0001;<br>LV-Control, Cocaine SA vs. LV-HCN4, Cocaine SA: 0.0252 |
| Fig. 5 | b | LV-Control, Cocaine SA: n = 9;<br>LV-HCN1, Cocaine SA: n = 5;<br>LV-HCN2, Cocaine SA: n = 12;<br>LV-HCN4, Cocaine SA: n = 7 | Shapiro-Wilk test | Pass | Two-way ANOVA | Voltage | 165.4 | / | 7, 232 | <0.0001 | LV-Control, Cocaine SA vs. LV-HCN1, Cocaine SA: 0.963;<br>LV-Control, Cocaine SA vs. LV-HCN2, Cocaine SA: <0.0001;<br>LV-Control, Cocaine SA vs. LV-HCN4, Cocaine SA: 0.0252 |
| Fig. 5 | c | LV-Control, Cocaine SA: n = 9<br>LV-HCN1, Cocaine SA: n = 5<br>LV-HCN2, Cocaine SA: n = 10<br>LV-HCN4, Cocaine SA: n = 7 | Shapiro-Wilk test | Pass | Two-way ANOVA | Intervention × Voltage | 0.3092 | / | 21, 232 | 0.9987 | LV-Control, Cocaine SA vs. LV-HCN1, Cocaine SA: > 0.9999;<br>LV-Control, Cocaine SA vs. LV-HCN2, Cocaine SA: 0.0174;<br>LV-Control, Cocaine SA vs. LV-HCN4, Cocaine SA: > 0.9999 |
| Fig. 5 | c | LV-Control, Cocaine SA: n = 9<br>LV-HCN1, Cocaine SA: n = 5<br>LV-HCN2, Cocaine SA: n = 10<br>LV-HCN4, Cocaine SA: n = 7 | Shapiro-Wilk test | Pass | Two-way ANOVA | Intervention | 2.868 | / | 3, 232 | 0.0373 | LV-Control, Cocaine SA vs. LV-HCN1, Cocaine SA: > 0.9999;<br>LV-Control, Cocaine SA vs. LV-HCN2, Cocaine SA: 0.0174;<br>LV-Control, Cocaine SA vs. LV-HCN4, Cocaine SA: > 0.9999 |
| Fig. 5 | c | LV-Control, Cocaine SA: n = 9<br>LV-HCN1, Cocaine SA: n = 5<br>LV-HCN2, Cocaine SA: n = 10<br>LV-HCN4, Cocaine SA: n = 7 | Shapiro-Wilk test | Pass | Two-way ANOVA | Voltage | 1685 | / | 7, 232 | <0.0001 | LV-Control, Cocaine SA vs. LV-HCN1, Cocaine SA: > 0.9999;<br>LV-Control, Cocaine SA vs. LV-HCN2, Cocaine SA: 0.0174;<br>LV-Control, Cocaine SA vs. LV-HCN4, Cocaine SA: > 0.9999 |
| Fig. 5 | d | LV-Control, Cocaine SA: n = 9;<br>LV-HCN1, Cocaine SA: n = 5;<br>LV-HCN2, Cocaine SA: n = 12; | Shapiro-Wilk test | Pass | One-way ANOVA | Intervention | 3.621 | / | 3, 29 | 0.0246 | LV-Control, Cocaine SA vs. LV-HCN1, Cocaine SA: 0.9956;<br>LV-Control, Cocaine SA vs. LV-HCN2, Cocaine SA: 0.0411;<br>LV-Control, Cocaine SA vs. LV-HCN4, Cocaine SA: 0.8607 |
|  |  | LV-HCN4, Cocaine SA: n = 7 |  |  |  |  |  |  |  |  |  |
| Fig. 5 | e | LV-Control, Cocaine SA: n = 9;<br>LV-HCN1, Cocaine SA: n = 5;<br>LV-HCN2, Cocaine SA: n = 10;<br>LV-HCN4, Cocaine SA: n = 7 | Shapiro-Wilk test | Pass | Two-way ANOVA | Intervention × Voltage | 1.237 | / | 21, 224 | 0.2222 | LV-Control, Cocaine SA vs. LV-HCN1, Cocaine SA: 0.7861;<br>LV-Control, Cocaine SA vs. LV-HCN2, Cocaine SA: <0.0001;<br>LV-Control, Cocaine SA vs. LV-HCN4, Cocaine SA: 0.0026 |
| Fig. 5 | e | LV-Control, Cocaine SA: n = 9;<br>LV-HCN1, Cocaine SA: n = 5;<br>LV-HCN2, Cocaine SA: n = 10;<br>LV-HCN4, Cocaine SA: n = 7 | Shapiro-Wilk test | Pass | Two-way ANOVA | Intervention | 11.03 | / | 3, 224 | <0.0001 | LV-Control, Cocaine SA vs. LV-HCN1, Cocaine SA: 0.7861;<br>LV-Control, Cocaine SA vs. LV-HCN2, Cocaine SA: <0.0001;<br>LV-Control, Cocaine SA vs. LV-HCN4, Cocaine SA: 0.0026 |
| Fig. 5 | e | LV-Control, Cocaine SA: n = 9;<br>LV-HCN1, Cocaine SA: n = 5;<br>LV-HCN2, Cocaine SA: n = 10;<br>LV-HCN4, Cocaine SA: n = 7 | Shapiro-Wilk test | Pass | Two-way ANOVA | Voltage | 183.1 | / | 7, 224 | <0.0001 | LV-Control, Cocaine SA vs. LV-HCN1, Cocaine SA: 0.7861;<br>LV-Control, Cocaine SA vs. LV-HCN2, Cocaine SA: <0.0001;<br>LV-Control, Cocaine SA vs. LV-HCN4, Cocaine SA: 0.0026 |
| Fig. 5 | f | LV-Control, Cocaine SA: n = 9;<br>LV-HCN1, Cocaine SA: n = 5;<br>LV-HCN2, Cocaine SA: n = 10;<br>LV-HCN4, Cocaine SA: n = 7 | Shapiro-Wilk test | Pass | Two-way ANOVA | Intervention × Voltage | 0.1635 | / | 21, 224 | > 0.9999 | LV-Control, Cocaine SA vs. LV-HCN1, Cocaine SA: > 0.9999;<br>LV-Control, Cocaine SA vs. LV-HCN2, Cocaine SA: 0.998;<br>LV-Control, Cocaine SA vs. LV-HCN4, Cocaine SA: 0.9952 |
| Fig. 5 | f | LV-Control, Cocaine SA: n = 9;<br>LV-HCN1, Cocaine SA: n = 5;<br>LV-HCN2, Cocaine SA: n = 10;<br>LV-HCN4, Cocaine SA: n = 7 | Shapiro-Wilk test | Pass | Two-way ANOVA | Intervention | 0.3226 | / | 3, 224 | 0.809 | LV-Control, Cocaine SA vs. LV-HCN1, Cocaine SA: > 0.9999;<br>LV-Control, Cocaine SA vs. LV-HCN2, Cocaine SA: 0.998;<br>LV-Control, Cocaine SA vs. LV-HCN4, Cocaine SA: 0.9952 |
| Fig. 5 | f | LV-Control, Cocaine SA: n = 9;<br>LV-HCN1, Cocaine SA: n = 5;<br>LV-HCN2, Cocaine SA: n = 10;<br>LV-HCN4, Cocaine SA: n = 7 | Shapiro-Wilk test | Pass | Two-way ANOVA | Voltage | 1540 | / | 7, 224 | <0.0001 | LV-Control, Cocaine SA vs. LV-HCN1, Cocaine SA: > 0.9999;<br>LV-Control, Cocaine SA vs. LV-HCN2, Cocaine SA: 0.998;<br>LV-Control, Cocaine SA vs. LV-HCN4, Cocaine SA: 0.9952 |
| Fig. 5 | g | LV-Control, Cocaine SA: n = 9;<br>LV-HCN1, Cocaine SA: n = 5;<br>LV-HCN2, Cocaine SA: n = 10; | Shapiro-Wilk test | Pass | One-way ANOVA | Intervention | 2.397 | / | 3, 28 | 0.0893 | LV-Control, Cocaine SA vs. LV-HCN1, Cocaine SA: 0.8613;<br>LV-Control, Cocaine SA vs. LV-HCN2, Cocaine SA: 0.0387;<br>LV-Control, Cocaine SA vs. LV-HCN4, Cocaine SA: 0.5273 |
|  |  | LV-HCN4, Cocaine SA: n = 7 |  |  |  |  |  |  |  |  |  |
| Fig. 5 | h | LV-Control, Cocaine SA: n = 19;<br>LV-HCN1, Cocaine SA: n = 23;<br>LV-HCN2, Cocaine SA: n = 33;<br>LV-HCN4, Cocaine SA: n = 22 | Shapiro-Wilk test | Pass | One-way ANOVA | Intervention | 13.13 | / | 3, 93 | <0.0001 | LV-Control, Cocaine SA vs. LV-HCN1, Cocaine SA: 0.9151;<br>LV-Control, Cocaine SA vs. LV-HCN2, Cocaine SA: <0.0001;<br>LV-Control, Cocaine SA vs. LV-HCN4, Cocaine SA: 0.4648 |
| Fig. 5 | i | LV-Control, Cocaine SA: n = 8;<br>LV-HCN1, Cocaine SA: n = 7;<br>LV-HCN2, Cocaine SA: n = 8;<br>LV-HCN4, Cocaine SA: n = 7 | Shapiro-Wilk test | Pass | Two-way ANOVA | Intervention × Nose poke type | 1.673 | / | 3, 50 | 0.1846 | LV-Control, Cocaine SA vs. LV-HCN1, Cocaine SA: 0.7676;<br>LV-Control, Cocaine SA vs. LV-HCN2, Cocaine SA: 0.0044;<br>LV-Control, Cocaine SA vs. LV-HCN4, Cocaine SA: 0.9727 |
| Fig. 5 | i | LV-Control, Cocaine SA: n = 8;<br>LV-HCN1, Cocaine SA: n = 7;<br>LV-HCN2, Cocaine SA: n = 8;<br>LV-HCN4, Cocaine SA: n = 7 | Shapiro-Wilk test | Pass | Two-way ANOVA | Intervention | 3.242 | / | 3, 50 | 0.0296 | LV-Control, Cocaine SA vs. LV-HCN1, Cocaine SA: 0.7676;<br>LV-Control, Cocaine SA vs. LV-HCN2, Cocaine SA: 0.0044;<br>LV-Control, Cocaine SA vs. LV-HCN4, Cocaine SA: 0.9727 |
| Fig. 5 | i | LV-Control, Cocaine SA: n = 8;<br>LV-HCN1, Cocaine SA: n = 7;<br>LV-HCN2, Cocaine SA: n = 8;<br>LV-HCN4, Cocaine SA: n = 7 | Shapiro-Wilk test | Pass | Two-way ANOVA | Nose poke type | 52.03 | / | 1, 50 | <0.0001 | LV-Control, Cocaine SA vs. LV-HCN1, Cocaine SA: 0.7676;<br>LV-Control, Cocaine SA vs. LV-HCN2, Cocaine SA: 0.0044;<br>LV-Control, Cocaine SA vs. LV-HCN4, Cocaine SA: 0.9727 |
| Fig. 5 | l | DIO-GFP: n = 8;<br>DIO-HCN2-shRNA: n = 9;<br>DIO-HCN4-shRNA: n = 7 | Shapiro-Wilk test | Pass | Two-way repeated-measures ANOVA | Time × Intervention | 6.453 | / | 2, 42 | 0.0036 | Extinction (DIO-GFP vs. DIO-HCN2-shRNA): >0.9999;<br>Extinction (DIO-GFP vs. DIO-HCN4-shRNA): >0.9999;<br>Test (DIO-GFP vs. DIO-HCN2-shRNA): 0.0004;<br>Test (DIO-GFP vs. DIO-HCN4-shRNA): >0.9999 |
| Fig. 5 | l | DIO-GFP: n = 8;<br>DIO-HCN2-shRNA: n = 9;<br>DIO-HCN4-shRNA: n = 7 | Shapiro-Wilk test | Pass | Two-way repeated-measures ANOVA | Time | 134.9 | / | 1, 42 | < 0.0001 | Extinction (DIO-GFP vs. DIO-HCN2-shRNA): >0.9999;<br>Extinction (DIO-GFP vs. DIO-HCN4-shRNA): >0.9999;<br>Test (DIO-GFP vs. DIO-HCN2-shRNA): 0.0004;<br>Test (DIO-GFP vs. DIO-HCN4-shRNA): >0.9999 |
| Fig. 5 | l | DIO-GFP: n = 8;<br>DIO-HCN2-shRNA: n = 9;<br>DIO-HCN4-shRNA: n = 7 | Shapiro-Wilk test | Pass | Two-way repeated-measures ANOVA | Intervention | 6.443 | / | 2, 42 | 0.0036 | Extinction (DIO-GFP vs. DIO-HCN2-shRNA): >0.9999;<br>Extinction (DIO-GFP vs. DIO-HCN4-shRNA): >0.9999;<br>Test (DIO-GFP vs. DIO-HCN2-shRNA): 0.0004;<br>Test (DIO-GFP vs. DIO-HCN4-shRNA): >0.9999 |
| Fig. 5 | n | DIO-GFP: n = 11;<br>DIO-HCN2-shRNA: n = 16;<br>DIO-HCN4-shRNA: n = 11; | Shapiro-Wilk test | Pass | One-way ANOVA | Intervention | 47.23 | / | 2, 35 | < 0.0001 | DIO-GFP vs. DIO-HCN2-shRNA: < 0.0001;<br>DIO-GFP vs. DIO-HCN4-shRNA: 0.1718 |
| Fig. 5 | p | DIO-GFP: n = 6;<br>DIO-HCN2-P2A-GFP: n = 8 | Shapiro-Wilk test | Pass | Two-way repeated-measures ANOVA | Time × Intervention | 5.215 | / | 1, 12 | 0.0414 | Extinction (DIO-GFP vs. DIO-HCN2-P2A-GFP): >0.9999;<br>Test (DIO-GFP vs. DIO-HCN2-P2A-GFP): 0.0045 |
| Fig. 5 | p | DIO-GFP: n = 6;<br>DIO-HCN2-P2A-GFP: n = 8 | Shapiro-Wilk test | Pass | Two-way repeated-measures ANOVA | Time | 82.81 | / | 1, 12 | < 0.0001 | Extinction (DIO-GFP vs. DIO-HCN2-P2A-GFP): >0.9999;<br>Test (DIO-GFP vs. DIO-HCN2-P2A-GFP): 0.0045 |
| Fig. 5 | p | DIO-GFP: n = 6;<br>DIO-HCN2-P2A-GFP: n = 8 | Shapiro-Wilk test | Pass | Two-way repeated-measures ANOVA | Intervention | 6.511 | / | 1, 12 | 0.0254 | Extinction (DIO-GFP vs. DIO-HCN2-P2A-GFP): >0.9999;<br>Test (DIO-GFP vs. DIO-HCN2-P2A-GFP): 0.0045 |
| Fig. 5 | r | DIO-GFP: n = 8;<br>DIO-HCN2-P2A-GFP: n = 8 | Shapiro-Wilk test | Pass | Unpaired t test | Intervention | / | 3.254 | 14 | 0.0058 | / |

**Supplementary Table 6.** Local and systemic HCN blockade suppress maladaptive cue-driven seeking and associated accumbal plasticity.

| Figure | Panel | Sample size | Normality test | Normality test results | Test | Comparison | F | t | df | P | Post hoc test |
| --- | --- | --- | --- | --- | --- | --- | --- | --- | --- | --- | --- |
| Fig. 6 | b | VEH: n = 6;<br>ZD7288: n = 8 | Shapiro-Wilk test | Pass | Two-way repeated-measures ANOVA | Treatment × Nose poke type | 2.753 | / | 1, 12 | 0.1229 | Active nose poke (Vehicle vs. ZD7288): 0.0152;<br>Inactive nose poke (Vehicle vs. ZD7288): 0.7399 |
| Fig. 6 | b | VEH: n = 6;<br>ZD7288: n = 8 | Shapiro-Wilk test | Pass | Two-way repeated-measures ANOVA | Treatment | 5.868 | / | 1, 12 | 0.0322 | Active nose poke (Vehicle vs. ZD7288): 0.0152;<br>Inactive nose poke (Vehicle vs. ZD7288): 0.7399 |
| Fig. 6 | b | VEH: n = 6;<br>ZD7288: n = 8 | Shapiro-Wilk test | Pass | Two-way repeated-measures ANOVA | Nose poke type | 31.48 | / | 1, 12 | 0.0001 | Active nose poke (Vehicle vs. ZD7288): 0.0152;<br>Inactive nose poke (Vehicle vs. ZD7288): 0.7399 |
| Fig. 6 | d | VEH: n = 18;<br>ZD7288: n = 21 | Shapiro-Wilk test | Pass | Unpaired t test | / | / | 2.299 | 37 | 0.0273 | / |
| Fig. 6 | e | VEH: n = 7;<br>ZD7288: n = 8 | Shapiro-Wilk test | Pass | Two-way ANOVA | Treatment × Nose poke type | 15.8 | / | 1, 26 | 0.0005 | Active nose poke (Vehicle vs. ZD7288): < 0.0001;<br>Inactive nose poke (Vehicle vs. ZD7288): 0.8266 |
| Fig. 6 | e | VEH: n = 7;<br>ZD7288: n = 8 | Shapiro-Wilk test | Pass | Two-way ANOVA | Treatment | 22.66 | / | 1, 26 | < 0.0001 | Active nose poke (Vehicle vs. ZD7288): < 0.0001;<br>Inactive nose poke (Vehicle vs. ZD7288): 0.8266 |
| Fig. 6 | e | VEH: n = 7;<br>ZD7288: n = 8 | Shapiro-Wilk test | Pass | Two-way ANOVA | Nose poke type | 33.1 | / | 1, 26 | < 0.0001 | Active nose poke (Vehicle vs. ZD7288): < 0.0001;<br>Inactive nose poke (Vehicle vs. ZD7288): 0.8266 |
| Fig. 6 | g | VEH: n = 17;<br>ZD7288: n = 17 | Shapiro-Wilk test | Pass | Unpaired t test | / | / | 6.998 | 32 | < 0.0001 | / |
| Fig. 6 | h | VEH: n = 11;<br>ZD7288: n = 10 | Shapiro-Wilk test | Pass | Two-way repeated-measures ANOVA | Treatment × Time | 11.12 | / | 1, 19 | 0.0035 | Last extinction (Vehicle vs. ZD7288): 0.9661;<br>Reinstatement test (Vehicle vs. ZD7288): 0.0005 |
| Fig. 6 | h | VEH: n = 11;<br>ZD7288: n = 10 | Shapiro-Wilk test | Pass | Two-way repeated-measures ANOVA | Treatment | 6.143 | / | 1, 19 | 0.0227 | Last extinction (Vehicle vs. ZD7288): 0.9661;<br>Reinstatement test (Vehicle vs. ZD7288): 0.0005 |
| Fig. 6 | h | VEH: n = 11;<br>ZD7288: n = 10 | Shapiro-Wilk test | Pass | Two-way repeated-measures ANOVA | Time | 23.35 | / | 1, 19 | 0.0001 | Last extinction (Vehicle vs. ZD7288): 0.9661;<br>Reinstatement test (Vehicle vs. ZD7288): 0.0005 |
| Fig. 6 | j | VEH: n = 6;<br>Ivabradine: n = 7 | Shapiro-Wilk test | Pass | Two-way ANOVA | Treatment × Nose poke type | 7.7357 | / | 1, 20 | 0.0134 | Active nose poke (Vehicle vs. Ivabradine): 0.0018;<br>Inactive nose poke (Vehicle vs. Ivabradine): 0.9983 |
| Fig. 6 | j | VEH: n = 6;<br>Ivabradine: n = 7 | Shapiro-Wilk test | Pass | Two-way ANOVA | Treatment | 7.769 | / | 1, 20 | 0.0144 | Active nose poke (Vehicle vs. Ivabradine): 0.0018;<br>Inactive nose poke (Vehicle vs. Ivabradine): 0.9983 |
| Fig. 6 | j | VEH: n = 6;<br>Ivabradine: n = 7 | Shapiro-Wilk test | Pass | Two-way ANOVA | Nose poke type | 30.45 | / | 1, 20 | <0.0001 | Active nose poke (Vehicle vs. Ivabradine): 0.0018;<br>Inactive nose poke (Vehicle vs. Ivabradine): 0.9983 |
| Fig. 6 | k | VEH: n = 8;<br>Ivabradine: n = 8 | Shapiro-Wilk test | Pass | Two-way ANOVA | Treatment × Nose poke type | 10.04 | / | 1, 28 | 0.0037 | Active nose poke (Vehicle vs. Ivabradine): 0.0002;<br>Inactive nose poke (Vehicle vs. Ivabradine): 0.9954 |
| Fig. 6 | k | VEH: n = 8;<br>Ivabradine: n = 8 | Shapiro-Wilk test | Pass | Two-way ANOVA | Treatment | 10.83 | / | 1, 28 | 0.0027 | Active nose poke (Vehicle vs. Ivabradine): 0.0002;<br>Inactive nose poke (Vehicle vs. Ivabradine): 0.9954 |
| Fig. 6 | k | VEH: n = 8;<br>Ivabradine: n = 8 | Shapiro-Wilk test | Pass | Two-way ANOVA | Nose poke type | 71.76 | / | 1, 28 | <0.0001 | Active nose poke (Vehicle vs. Ivabradine): 0.0002;<br>Inactive nose poke (Vehicle vs. Ivabradine): 0.9954 |
| Fig. 6 | l | VEH: n = 10;<br>Ivabradine: n = 10 | Shapiro-Wilk test | Pass | Two-way ANOVA | Treatment × Nose poke type | 3.657 | / | 1, 36 | 0.0638 | Active nose poke (Vehicle vs. Ivabradine): 0.0036;<br>Inactive nose poke (Vehicle vs. Ivabradine): 0.7589 |
| Fig. 6 | l | VEH: n = 10;<br>Ivabradine: n = 10 | Shapiro-Wilk test | Pass | Two-way ANOVA | Treatment | 8.156 | / | 1, 36 | 0.0071 | Active nose poke (Vehicle vs. Ivabradine): 0.0036;<br>Inactive nose poke (Vehicle vs. Ivabradine): 0.7589 |
| Fig. 6 | l | VEH: n = 10;<br>Ivabradine: n = 10 | Shapiro-Wilk test | Pass | Two-way ANOVA | Nose poke type | 60.09 | / | 1, 36 | <0.0001 | Active nose poke (Vehicle vs. Ivabradine): 0.0036;<br>Inactive nose poke (Vehicle vs. Ivabradine): 0.7589 |

**Supplementary Table 7.** Cocaine-paired cues do not evoke significant ACh transients in the NAcSh.

| Figure | Panel | Sample size | Normality test | Normality test results | Test | Comparison | *F* | *t* | df | *P* | Post hoc test |
| --- | --- | --- | --- | --- | --- | --- | --- | --- | --- | --- | --- |
| Extended Data Fig. 1 | e | GFP: n = 6;<br>ACh: n = 7 | Shapiro-Wilk test | Pass | Unpaired t test | Intervention | / | 0.4643 | 11 | 0.6515 | / |

**Supplementary Table 8.** Optogenetic inhibition of NAcSh CINs has no effect on cue-induced cocaine seeking.

| Figure | Panel | Sample size | Normality test | Normality test results | Test | Comparison | F | t | df | P | Post hoc test |
| --- | --- | --- | --- | --- | --- | --- | --- | --- | --- | --- | --- |
| Extended Data Fig. 2 | c | DIO-GFP: n = 5;<br>DIO-NpHR: n = 6 | Shapiro-Wilk test | Pass | Two-way repeated-measures ANOVA | Time × Intervention | 0.5149 | / | 4, 36 | 0.7252 | / |
| Extended Data Fig. 2 | c | DIO-GFP: n = 5;<br>DIO-NpHR: n = 6 | Shapiro-Wilk test | Pass | Two-way repeated-measures ANOVA | Time | 10.44 | / | 4, 36 | < 0.0001 | / |
| Extended Data Fig. 2 | c | DIO-GFP: n = 5;<br>DIO-NpHR: n = 6 | Shapiro-Wilk test | Pass | Two-way repeated-measures ANOVA | Intervention | 0.1064 | / | 1, 9 | 0.7517 | / |
| Extended Data Fig. 2 | d | DIO-GFP: n = 5;<br>DIO-NpHR: n = 6 | Shapiro-Wilk test | Pass | Two-way repeated-measures ANOVA | Time × Intervention | 0.2892 | / | 9, 81 | 0.9759 | / |
| Extended Data Fig. 2 | d | DIO-GFP: n = 5;<br>DIO-NpHR: n = 6 | Shapiro-Wilk test | Pass | Two-way repeated-measures ANOVA | Time | 157.4 | / | 9, 81 | < 0.0001 | / |
| Extended Data Fig. 2 | d | DIO-GFP: n = 5;<br>DIO-NpHR: n = 6 | Shapiro-Wilk test | Pass | Two-way repeated-measures ANOVA | Intervention | 0.2085 | / | 1, 9 | 0.6588 | / |
| Extended Data Fig. 2 | e | DIO-GFP: n = 5;<br>DIO-NpHR: n = 6 | Shapiro-Wilk test | Pass | Two-way repeated-measures ANOVA | Time × Intervention | 0.028 | / | 1, 9 | 0.8718 | / |
| Extended Data Fig. 2 | e | DIO-GFP: n = 5;<br>DIO-NpHR: n = 6 | Shapiro-Wilk test | Pass | Two-way repeated-measures ANOVA | Time | 23.19 | / | 1, 9 | 0.001 | / |
| Extended Data Fig. 2 | e | DIO-GFP: n = 5;<br>DIO-NpHR: n = 6 | Shapiro-Wilk test | Pass | Two-way repeated-measures ANOVA | Intervention | 0.1155 | / | 1, 9 | 0.7418 | / |
| Extended Data Fig. 2 | f | DIO-GFP: n = 5;<br>DIO-NpHR: n = 6 | Shapiro-Wilk test | Pass | Two-way repeated-measures ANOVA | Time × Intervention | 0.02 | / | 1, 9 | 0.8895 | / |
| Extended Data Fig. 2 | f | DIO-GFP: n = 5;<br>DIO-NpHR: n = 6 | Shapiro-Wilk test | Pass | Two-way repeated-measures ANOVA | Time | 0.1985 | / | 1, 9 | 0.6665 | / |
| Extended Data Fig. 2 | f | DIO-GFP: n = 5;<br>DIO-NpHR: n = 6 | Shapiro-Wilk test | Pass | Two-way repeated-measures ANOVA | Intervention | 0.1925 | / | 1, 9 | 0.6712 | / |

**Supplementary Table 9.** Cocaine SA, but not sucrose SA, increases ISR markers in CINs and D1- and D2-expressing neurons.

| Figure | Panel | Sample size | Normality test | Normality test results | Test | Comparison | *F* | *t* | df | *P* | Post hoc test |
| --- | --- | --- | --- | --- | --- | --- | --- | --- | --- | --- | --- |
| Extended Data Fig. 3 | b | Saline: n = 22 from 3 rats;<br>Sucrose: n = 45 from 3 rats;<br>Cocaine: n = 55 from 3 rats. | Shapiro-Wilk test | Pass | One-way ANOVA | Drug | 30.7 | / | 2, 119 | <0.0001 | Saline vs. Sucrose: 0.6757;<br>Saline vs. Cocaine: <0.0001 |
| Extended Data Fig. 3 | d | Saline: n = 51 from 3 rats;<br>Sucrose: n = 47 from 3 rats;<br>Cocaine: n = 52 from 3 rats. | Shapiro-Wilk test | Pass | One-way ANOVA | Drug | 102.1 | / | 2, 147 | <0.0001 | Saline vs. Sucrose: 0.9059;<br>Saline vs. Cocaine: <0.0001 |
| Extended Data Fig. 3 | f | Saline: n = 40 from 3 rats;<br>Sucrose: n = 43 from 3 rats;<br>Cocaine: n = 57 from 3 rats. | Shapiro-Wilk test | Pass | One-way ANOVA | Drug | 249.6 | / | 2, 137 | <0.0001 | Saline vs. Sucrose: 0.3786;<br>Saline vs. Cocaine: <0.0001 |
| Extended Data Fig. 3 | h | Saline: n = 46 from 3 rats;<br>Sucrose: n = 49 from 3 rats;<br>Cocaine: n = 53 from 3 rats. | Shapiro-Wilk test | Pass | One-way ANOVA | Drug | 1970 | / | 2, 145 | <0.0001 | Saline vs. Sucrose: 0.983;<br>Saline vs. Cocaine: <0.0001 |

**Supplementary Table 10.** Elevating ISR signaling during sucrose training alters CIN excitability and cue-evoked ACh dynamics without enhancing sucrose seeking.

| Figure | Panel | Sample size | Normality test | Normality test results | Test | Comparison | *F* | *t* | df | *P* | Post hoc test |
| --- | --- | --- | --- | --- | --- | --- | --- | --- | --- | --- | --- |
| Extended Data Fig. 4 | b | VEH: n = 6;<br>Sal003: n = 6 | Shapiro-Wilk test | Pass | Unpaired t test | Drug | / | 0.3909 | 10 | 0.7041 | / |
| Extended Data Fig. 4 | d | VEH: n = 17 from 3 rats;<br>Sal003: n = 17 from 3 rats. | Shapiro-Wilk test | Pass | Unpaired t test | Treatment | / | 3.534 | 32 | 0.0013 | / |
| Extended Data Fig. 4 | g | VEH: n = 5;<br>Sal003: n = 5 | Shapiro-Wilk test | Pass | Unpaired t test | Drug | / | 4.396 | 8 | 0.0023 | / |

**Supplementary Table 11.** Pharmacological manipulation of ISR signaling bidirectionally regulates p-eIF2α levels in CINs.

| Figure | Panel | Sample size | Normality test | Normality test results | Test | Comparison | *F* | *t* | df | *P* | Post hoc test |
| --- | --- | --- | --- | --- | --- | --- | --- | --- | --- | --- | --- |
| Extended Data Fig. 5 | b | Control: n = 93 from 3 rats;<br>ISRIB: n = 93 from 3 rats;<br>Sal003: n = 128 from 3 rats. | Shapiro-Wilk test | Pass | One-way ANOVA | Treatment | 65.69 | / | 2, 311 | <0.0001 | Control vs. ISRIB: 0.0156;<br>Control vs. Sal003: <0.0001 |

**Supplementary Table 12.** CIN-specific ISR suppression does not alter cocaine acquisition, extinction, or inactive responding.

| Figure | Panel | Sample size | Normality test | Normality test results | Test | Comparison | F | t | df | P | Post hoc test |
| --- | --- | --- | --- | --- | --- | --- | --- | --- | --- | --- | --- |
| Extended Data Fig. 6 | a | DIO-GFP: n = 8;<br>DIO-Ppp1r15b-P2A-GFP: n = 7 | Shapiro-Wilk test | Pass | Two-way repeated-measures ANOVA | Time × Intervention | 0.4093 | / | 9, 117 | 0.9280 | day 1: >0.9999;<br>day 2: >0.9999;<br>day 3: >0.9999;<br>day 4: >0.9999;<br>day 5: 0.9994;<br>day 6: 0.9980;<br>day 7: 0.9978;<br>day 8: >0.9999;<br>day 9: 0.9984;<br>day 10: >0.9999 |
| Extended Data Fig. 6 | a | DIO-GFP: n = 8;<br>DIO-Ppp1r15b-P2A-GFP: n = 7 | Shapiro-Wilk test | Pass | Two-way repeated-measures ANOVA | Time | 46.87 | / | 9, 117 | < 0.0001 | day 1: >0.9999;<br>day 2: >0.9999;<br>day 3: >0.9999;<br>day 4: >0.9999;<br>day 5: 0.9994;<br>day 6: 0.9980;<br>day 7: 0.9978;<br>day 8: >0.9999;<br>day 9: 0.9984;<br>day 10: >0.9999 |
| Extended Data Fig. 6 | a | DIO-GFP: n = 8;<br>DIO-Ppp1r15b-P2A-GFP: n = 7 | Shapiro-Wilk test | Pass | Two-way repeated-measures ANOVA | Intervention | 0.2073 | / | 1, 13 | 0.6564 | day 1: >0.9999;<br>day 2: >0.9999;<br>day 3: >0.9999;<br>day 4: >0.9999;<br>day 5: 0.9994;<br>day 6: 0.9980;<br>day 7: 0.9978;<br>day 8: >0.9999;<br>day 9: 0.9984;<br>day 10: >0.9999 |
| Extended Data Fig. 6 | b | DIO-GFP: n = 8;<br>DIO-Ppp1r15b-P2A-GFP: n = 7 | Shapiro-Wilk test | Pass | Two-way repeated-measures ANOVA | Time × Intervention | 0.2129 | / | 9, 117 | 0.9921 | day 1: 0.9994;<br>day 2: >0.9999;<br>day 3: >0.9999;<br>day 4: >0.9999;<br>day 5: 0.9986;<br>day 6: 0.9984;<br>day 7: >0.9999;<br>day 8: >0.9999;<br>day 9: >0.9999;<br>day 10: >0.9999 |
| Extended Data Fig. 6 | b | DIO-GFP: n = 8;<br>DIO-Ppp1r15b-P2A-GFP: n = 7 | Shapiro-Wilk test | Pass | Two-way repeated-measures ANOVA | Time | 30.91 | / | 9, 117 | < 0.0001 | day 1: 0.9994;<br>day 2: >0.9999;<br>day 3: >0.9999;<br>day 4: >0.9999;<br>day 5: 0.9986;<br>day 6: 0.9984;<br>day 7: >0.9999;<br>day 8: >0.9999;<br>day 9: >0.9999;<br>day 10: >0.9999 |
| Extended Data Fig. 6 | b | DIO-GFP: n = 8;<br>DIO-Ppp1r15b-P2A-GFP: n = 7 | Shapiro-Wilk test | Pass | Two-way repeated-measures ANOVA | Intervention | 0.0809 | / | 1, 13 | 0.7805 | day 1: 0.9994;<br>day 2: >0.9999;<br>day 3: >0.9999;<br>day 4: >0.9999;<br>day 5: 0.9986;<br>day 6: 0.9984; |
|  |  |  |  |  |  |  |  |  |  |  | day 7: >0.9999;<br>day 8: >0.9999;<br>day 9: >0.9999;<br>day 10: >0.9999 |
| Extended Data Fig. 6 | c | DIO-GFP: n = 8;<br>DIO-Ppp1r15b-P2A-GFP: n = 7 | Shapiro-Wilk test | Pass | Two-way repeated-measures ANOVA | Time × Intervention | 0.006 | / | 1, 13 | 0.941 | Extinction (DIO-GFP vs. DIO-Ppp1r15b-P2A-GFP): 0.9945;<br>Test (DIO-GFP vs. DIO-Ppp1r15b-P2A-GFP): 0.9819 |
| Extended Data Fig. 6 | c | DIO-GFP: n = 8;<br>DIO-Ppp1r15b-P2A-GFP: n = 7 | Shapiro-Wilk test | Pass | Two-way repeated-measures ANOVA | Time | 0.037 | / | 1, 13 | 0.8501 | Extinction (DIO-GFP vs. DIO-Ppp1r15b-P2A-GFP): 0.9945;<br>Test (DIO-GFP vs. DIO-Ppp1r15b-P2A-GFP): 0.9819 |
| Extended Data Fig. 6 | c | DIO-GFP: n = 8;<br>DIO-Ppp1r15b-P2A-GFP: n = 7 | Shapiro-Wilk test | Pass | Two-way repeated-measures ANOVA | Intervention | 0.024 | / | 1, 13 | 0.8797 | Extinction (DIO-GFP vs. DIO-Ppp1r15b-P2A-GFP): 0.9945;<br>Test (DIO-GFP vs. DIO-Ppp1r15b-P2A-GFP): 0.9819 |

**Supplementary Table 13.** Genetic suppression of ISR signaling in NAc D1R- or D2R-expressing neurons does not affect cue-induced cocaine seeking.

| Figure | Panel | Sample size | Normality test | Normality test results | Test | Comparison | F | t | df | P | Post hoc test |
| --- | --- | --- | --- | --- | --- | --- | --- | --- | --- | --- | --- |
| Extended Data Fig. 7 | c | DIO-GFP: n = 9;<br>DIO-Ppp1r15b-P2A-GFP: n = 7 | Shapiro-Wilk test | Pass | Two-way repeated-measures ANOVA | Time × Intervention | 0.0651 | / | 4, 56 | 0.9920 | / |
| Extended Data Fig. 7 | c | DIO-GFP: n = 9;<br>DIO-Ppp1r15b-P2A-GFP: n = 7 | Shapiro-Wilk test | Pass | Two-way repeated-measures ANOVA | Time | 8.011 | / | 4, 56 | < 0.0001 | / |
| Extended Data Fig. 7 | c | DIO-GFP: n = 9;<br>DIO-Ppp1r15b-P2A-GFP: n = 7 | Shapiro-Wilk test | Pass | Two-way repeated-measures ANOVA | Intervention | 0.0407 | / | 1, 14 | 0.843 | / |
| Extended Data Fig. 7 | d | DIO-GFP: n = 9;<br>DIO-Ppp1r15b-P2A-GFP: n = 7 | Shapiro-Wilk test | Pass | Two-way repeated-measures ANOVA | Time × Intervention | 0.7653 | / | 9, 126 | 0.6486 | / |
| Extended Data Fig. 7 | d | DIO-GFP: n = 9;<br>DIO-Ppp1r15b-P2A-GFP: n = 7 | Shapiro-Wilk test | Pass | Two-way repeated-measures ANOVA | Time | 190.1 | / | 9, 126 | < 0.0001 | / |
| Extended Data Fig. 7 | d | DIO-GFP: n = 9;<br>DIO-Ppp1r15b-P2A-GFP: n = 7 | Shapiro-Wilk test | Pass | Two-way repeated-measures ANOVA | Intervention | 3.003 | / | 1, 14 | 0.105 | / |
| Extended Data Fig. 7 | e | DIO-GFP: n = 9;<br>DIO-Ppp1r15b-P2A-GFP: n = 7 | Shapiro-Wilk test | Pass | Two-way repeated-measures ANOVA | Time × Intervention | 0.073 | / | 1, 14 | 0.791 | / |
| Extended Data Fig. 7 | e | DIO-GFP: n = 9;<br>DIO-Ppp1r15b-P2A-GFP: n = 7 | Shapiro-Wilk test | Pass | Two-way repeated-measures ANOVA | Time | 87.79 | / | 1, 14 | <0.0001 | / |
| Extended Data Fig. 7 | e | DIO-GFP: n = 9;<br>DIO-Ppp1r15b-P2A-GFP: n = 7 | Shapiro-Wilk test | Pass | Two-way repeated-measures ANOVA | Intervention | 0.001 | / | 1, 14 | 0.996 | / |
| Extended Data Fig. 7 | f | DIO-GFP: n = 9;<br>DIO-Ppp1r15b-P2A-GFP: n = 7 | Shapiro-Wilk test | Pass | Two-way repeated-measures ANOVA | Time × Intervention | 1.281 | / | 1, 14 | 0.2767 | / |
| Extended Data Fig. 7 | f | DIO-GFP: n = 9;<br>DIO-Ppp1r15b-P2A-GFP: n = 7 | Shapiro-Wilk test | Pass | Two-way repeated-measures ANOVA | Time | 2.096 | / | 1, 14 | 0.1697 | / |
| Extended Data Fig. 7 | f | DIO-GFP: n = 9;<br>DIO-Ppp1r15b-P2A-GFP: n = 7 | Shapiro-Wilk test | Pass | Two-way repeated-measures ANOVA | Intervention | 0.1 | / | 1, 14 | 0.7563 | / |
| Extended Data Fig. 7 | i | DIO-GFP: n = 8;<br>DIO-Ppp1r15b-P2A-GFP: n = 7 | Shapiro-Wilk test | Pass | Two-way repeated-measures ANOVA | Time × Intervention | 0.2778 | / | 4, 52 | 0.8910 | / |
| Extended Data Fig. 7 | i | DIO-GFP: n = 8;<br>DIO-Ppp1r15b-P2A-GFP: n = 7 | Shapiro-Wilk test | Pass | Two-way repeated-measures ANOVA | Time | 4.122 | / | 4, 52 | 0.0056 | / |
| Extended Data Fig. 7 | i | DIO-GFP: n = 8;<br>DIO-Ppp1r15b-P2A-GFP: n = 7 | Shapiro-Wilk test | Pass | Two-way repeated-measures ANOVA | Intervention | 0.2737 | / | 1, 13 | 0.6097 | / |
| Extended Data Fig. 7 | j | DIO-GFP: n = 8;<br>DIO-Ppp1r15b-P2A-GFP: n = 7 | Shapiro-Wilk test | Pass | Two-way repeated-measures ANOVA | Time × Intervention | 1.477 | / | 9, 117 | 0.1646 | / |
| Extended Data Fig. 7 | j | DIO-GFP: n = 8; | Shapiro-Wilk test | Pass | Two-way repeated-measures ANOVA | Time | 178.6 | / | 9, 117 | < 0.0001 | / |
|  |  | DIO-Ppp1r15b-P2A-GFP:<br>n = 7 |  |  |  |  |  |  |  |  |  |
| Extended Data Fig. 7 | j | DIO-GFP: n = 8;<br>DIO-Ppp1r15b-P2A-GFP:<br>n = 7 | Shapiro-Wilk test | Pass | Two-way repeated-measures ANOVA | Intervention | 2.837 | / | 1, 13 | 0.1159 | / |
| Extended Data Fig. 7 | k | DIO-GFP: n = 8;<br>DIO-Ppp1r15b-P2A-GFP:<br>n = 7 | Shapiro-Wilk test | Pass | Two-way repeated-measures ANOVA | Time × Intervention | 0.0614 | / | 1, 13 | 0.8082 | / |
| Extended Data Fig. 6 | k | DIO-GFP: n = 8;<br>DIO-Ppp1r15b-P2A-GFP:<br>n = 7 | Shapiro-Wilk test | Pass | Two-way repeated-measures ANOVA | Time | 149.3 | / | 1, 13 | <0.0001 | / |
| Extended Data Fig. 7 | k | DIO-GFP: n = 8;<br>DIO-Ppp1r15b-P2A-GFP:<br>n = 7 | Shapiro-Wilk test | Pass | Two-way repeated-measures ANOVA | Intervention | 0.3376 | / | 1, 13 | 0.5712 | / |
| Extended Data Fig. 7 | l | DIO-GFP: n = 8;<br>DIO-Ppp1r15b-P2A-GFP:<br>n = 7 | Shapiro-Wilk test | Pass | Two-way repeated-measures ANOVA | Time × Intervention | 0.001 | / | 1, 13 | 0.9968 | / |
| Extended Data Fig. 7 | l | DIO-GFP: n = 8;<br>DIO-Ppp1r15b-P2A-GFP:<br>n = 7 | Shapiro-Wilk test | Pass | Two-way repeated-measures ANOVA | Time | 0.154 | / | 1, 13 | 0.7011 | / |
| Extended Data Fig. 7 | l | DIO-GFP: n = 8;<br>DIO-Ppp1r15b-P2A-GFP:<br>n = 7 | Shapiro-Wilk test | Pass | Two-way repeated-measures ANOVA | Intervention | 0.4 | / | 1, 13 | 0.5381 | / |

**Supplementary Table 14.** Pharmacological manipulation of ISR signaling bidirectionally regulates HCN2 protein expression in CINs.

| Figure | Panel | Sample size | Normality test | Normality test results | Test | Comparison | *F* | *t* | df | *P* | Post hoc test |
| --- | --- | --- | --- | --- | --- | --- | --- | --- | --- | --- | --- |
| Extended Data Fig. 8 | c | Control: n = 83 from 3 rats;<br>ISRIB: n = 86 from 3 rats;<br>Sal003: n = 69 from 3 rats. | Shapiro-Wilk test | Pass | One-way ANOVA | Treatment | / | 60.45 | 2, 234 | <0.0001 | Control vs. ISRIB: 0.0104;<br>Control vs. Sal003: <0.0001 |

**Supplementary Table 15.** Cocaine training increases maximal and available functional $I_h$ in CINs.

| Figure | Panel | Sample size | Normality test | Normality test results | Test | Comparison | *F* | *t* | df | *P* | Post hoc test |
| --- | --- | --- | --- | --- | --- | --- | --- | --- | --- | --- | --- |
| Extended Data Fig. 9 | b | -50 mV Saline SA: n = 14;<br>Cocaine SA: n = 9;<br>-60 mV Saline SA: n = 14;<br>Cocaine SA: n = 9;<br>-70 mV Saline SA: n = 14;<br>Cocaine SA: n = 9;<br>-80 mV Saline SA: n = 14;<br>Cocaine SA: n = 9;<br>-90 mV Saline SA: n = 14;<br>Cocaine SA: n = 9;<br>-100 mV Saline SA: n = 14;<br>Cocaine SA: n = 9;<br>-110 mV Saline SA: n = 14;<br>Cocaine SA: n = 9;<br>-120 mV Saline SA: n = 14;<br>Cocaine SA: n = 9 | Shapiro-Wilk test | Pass | Unpaired t test | Drug | / | 0.511 | 21 | 0.6147 | / |
| Extended Data Fig. 9 | b | -50 mV Saline SA: n = 14;<br>Cocaine SA: n = 9;<br>-60 mV Saline SA: n = 14;<br>Cocaine SA: n = 9;<br>-70 mV Saline SA: n = 14;<br>Cocaine SA: n = 9;<br>-80 mV Saline SA: n = 14;<br>Cocaine SA: n = 9;<br>-90 mV Saline SA: n = 14;<br>Cocaine SA: n = 9;<br>-100 mV Saline SA: n = 14;<br>Cocaine SA: n = 9;<br>-110 mV Saline SA: n = 14;<br>Cocaine SA: n = 9;<br>-120 mV Saline SA: n = 14;<br>Cocaine SA: n = 9 | Shapiro-Wilk test | Pass | Unpaired t test | Drug | / | 2.118 | 21 | 0.0462 | / |
| Extended Data Fig. 9 | b | -50 mV Saline SA: n = 14;<br>Cocaine SA: n = 9;<br>-60 mV Saline SA: n = 14;<br>Cocaine SA: n = 9;<br>-70 mV Saline SA: n = 14;<br>Cocaine SA: n = 9;<br>-80 mV Saline SA: n = 14;<br>Cocaine SA: n = 9;<br>-90 mV Saline SA: n = 14;<br>Cocaine SA: n = 9;<br>-100 mV Saline SA: n = 14;<br>Cocaine SA: n = 9;<br>-110 mV Saline SA: n = 14;<br>Cocaine SA: n = 9;<br>-120 mV Saline SA: n = 14;<br>Cocaine SA: n = 9 | Shapiro-Wilk test | Pass | Unpaired t test | Drug | / | 2.537 | 21 | 0.0192 | / |
| Extended Data Fig. 9 | b | -50 mV Saline SA: n = 14;<br>Cocaine SA: n = 9;<br>-60 mV Saline SA: n = 14;<br>Cocaine SA: n = 9;<br>-70 mV Saline SA: n = 14;<br>Cocaine SA: n = 9;<br>-80 mV Saline SA: n = 14;<br>Cocaine SA: n = 9;<br>-90 mV Saline SA: n = 14;<br>Cocaine SA: n = 9;<br>-100 mV Saline SA: n = 14;<br>Cocaine SA: n = 9;<br>-110 mV Saline SA: n = 14;<br>Cocaine SA: n = 9;<br>-120 mV Saline SA: n = 14;<br>Cocaine SA: n = 9 | Shapiro-Wilk test | Pass | Unpaired t test | Drug | / | 2.408 | 21 | 0.0253 | / |

| Figure | Panel | Sample size | Normality test | Normality test results | Test | Comparison | F | t | df | P | Post hoc test |
| --- | --- | --- | --- | --- | --- | --- | --- | --- | --- | --- | --- |
|  |  | -90 mV Saline SA: n = 14;<br>Cocaine SA: n = 9;<br>-100 mV Saline SA: n = 14; Cocaine SA: n = 9;<br>-110 mV Saline SA: n = 14; Cocaine SA: n = 9;<br>-120 mV Saline SA: n = 14; Cocaine SA: n = 9 |  |  |  |  |  |  |  |  |  |
| Extended Data Fig. 9 | b | -50 mV Saline SA: n = 14; Cocaine SA: n = 9;<br>-60 mV Saline SA: n = 14; Cocaine SA: n = 9;<br>-70 mV Saline SA: n = 14; Cocaine SA: n = 9;<br>-80 mV Saline SA: n = 14; Cocaine SA: n = 9;<br>-90 mV Saline SA: n = 14; Cocaine SA: n = 9;<br>-100 mV Saline SA: n = 14; Cocaine SA: n = 9;<br>-110 mV Saline SA: n = 14; Cocaine SA: n = 9;<br>-120 mV Saline SA: n = 14; Cocaine SA: n = 9 | Shapiro-Wilk test | Pass | Unpaired t test | Drug | / | 2.631 | 21 | 0.0156 | / |
| Extended Data Fig. 9 | b | -50 mV Saline SA: n = 14; Cocaine SA: n = 9;<br>-60 mV Saline SA: n = 14; Cocaine SA: n = 9;<br>-70 mV Saline SA: n = 14; Cocaine SA: n = 9;<br>-80 mV Saline SA: n = 14; Cocaine SA: n = 9;<br>-90 mV Saline SA: n = 14; Cocaine SA: n = 9;<br>-100 mV Saline SA: n = 14; Cocaine SA: n = 9;<br>-110 mV Saline SA: n = 14; Cocaine SA: n = 9;<br>-120 mV Saline SA: n = 14; Cocaine SA: n = 9 | Shapiro-Wilk test | Pass | Unpaired t test | Drug | / | 2.661 | 21 | 0.0146 | / |
| Extended Data Fig. 9 | b | -50 mV Saline SA: n = 14; Cocaine SA: n = 9;<br>-60 mV Saline SA: n = 14; Cocaine SA: n = 9;<br>-70 mV Saline SA: n = 14; Cocaine SA: n = 9;<br>-80 mV Saline SA: n = 14; Cocaine SA: n = 9;<br>-90 mV Saline SA: n = 14; Cocaine SA: n = 9;<br>-100 mV Saline SA: n = 14; Cocaine SA: n = 9;<br>-110 mV Saline SA: n = 14; Cocaine SA: n = 9;<br>-120 mV Saline SA: n = 14; Cocaine SA: n = 9 | Shapiro-Wilk test | Pass | Unpaired t test | Drug | / | 3.101 | 21 | 0.0054 | / |
| Extended Data Fig. 9 | b | -50 mV Saline SA: n = 14; Cocaine SA: n = 9; | Shapiro-Wilk test | Pass | Unpaired t test | Drug | / | 3.4 | 21 | 0.0027 | / |
|  |  | -60 mV Saline SA: n = 14;<br>Cocaine SA: n = 9;<br>-70 mV Saline SA: n = 14;<br>Cocaine SA: n = 9;<br>-80 mV Saline SA: n = 14;<br>Cocaine SA: n = 9;<br>-90 mV Saline SA: n = 14;<br>Cocaine SA: n = 9;<br>-100 mV Saline SA: n = 14;<br>Cocaine SA: n = 9;<br>-110 mV Saline SA: n = 14;<br>Cocaine SA: n = 9;<br>-120 mV Saline SA: n = 14;<br>Cocaine SA: n = 9 |  |  |  |  |  |  |  |  |  |
| Extended Data Fig. 9 | c | -50 mV Saline SA: n = 14;<br>Cocaine SA: n = 9;<br>-60 mV Saline SA: n = 14;<br>Cocaine SA: n = 9;<br>-70 mV Saline SA: n = 14;<br>Cocaine SA: n = 9;<br>-80 mV Saline SA: n = 14;<br>Cocaine SA: n = 9;<br>-90 mV Saline SA: n = 14;<br>Cocaine SA: n = 9;<br>-100 mV Saline SA: n = 14;<br>Cocaine SA: n = 9;<br>-110 mV Saline SA: n = 14;<br>Cocaine SA: n = 9;<br>-120 mV Saline SA: n = 14;<br>Cocaine SA: n = 9 | Shapiro-Wilk test | Pass | Unpaired t test | Drug | / | 0.2768 | 21 | 0.7846 | / |
| Extended Data Fig. 9 | c | -50 mV Saline SA: n = 14;<br>Cocaine SA: n = 9;<br>-60 mV Saline SA: n = 14;<br>Cocaine SA: n = 9;<br>-70 mV Saline SA: n = 14;<br>Cocaine SA: n = 9;<br>-80 mV Saline SA: n = 14;<br>Cocaine SA: n = 9;<br>-90 mV Saline SA: n = 14;<br>Cocaine SA: n = 9;<br>-100 mV Saline SA: n = 14;<br>Cocaine SA: n = 9;<br>-110 mV Saline SA: n = 14;<br>Cocaine SA: n = 9;<br>-120 mV Saline SA: n = 14;<br>Cocaine SA: n = 9 | Shapiro-Wilk test | Pass | Unpaired t test | Drug | / | 1.023 | 21 | 0.3179 | / |
| Extended Data Fig. 9 | c | -50 mV Saline SA: n = 14;<br>Cocaine SA: n = 9;<br>-60 mV Saline SA: n = 14;<br>Cocaine SA: n = 9;<br>-70 mV Saline SA: n = 14;<br>Cocaine SA: n = 9;<br>-80 mV Saline SA: n = 14;<br>Cocaine SA: n = 9;<br>-90 mV Saline SA: n = 14;<br>Cocaine SA: n = 9;<br>-100 mV Saline SA: n = 14;<br>Cocaine SA: n = 9;<br>-110 mV Saline SA: n = 14;<br>Cocaine SA: n = 9;<br>-120 mV Saline SA: n = 14;<br>Cocaine SA: n = 9 | Shapiro-Wilk test | Pass | Unpaired t test | Drug | / | 0.5724 | 21 | 0.5731 | / |
|  |  | -120 mV Saline SA: n = 14; Cocaine SA: n = 9 |  |  |  |  |  |  |  |  |  |
| Extended Data Fig. 9 | c | -50 mV Saline SA: n = 14; Cocaine SA: n = 9;<br>-60 mV Saline SA: n = 14; Cocaine SA: n = 9;<br>-70 mV Saline SA: n = 14; Cocaine SA: n = 9;<br>-80 mV Saline SA: n = 14; Cocaine SA: n = 9;<br>-90 mV Saline SA: n = 14; Cocaine SA: n = 9;<br>-100 mV Saline SA: n = 14; Cocaine SA: n = 9;<br>-110 mV Saline SA: n = 14; Cocaine SA: n = 9;<br>-120 mV Saline SA: n = 14; Cocaine SA: n = 9 | Shapiro-Wilk test | Pass | Unpaired t test | Drug | / | 0.5121 | 21 | 0.614 | / |
| Extended Data Fig. 9 | c | -50 mV Saline SA: n = 14; Cocaine SA: n = 9;<br>-60 mV Saline SA: n = 14; Cocaine SA: n = 9;<br>-70 mV Saline SA: n = 14; Cocaine SA: n = 9;<br>-80 mV Saline SA: n = 14; Cocaine SA: n = 9;<br>-90 mV Saline SA: n = 14; Cocaine SA: n = 9;<br>-100 mV Saline SA: n = 14; Cocaine SA: n = 9;<br>-110 mV Saline SA: n = 14; Cocaine SA: n = 9;<br>-120 mV Saline SA: n = 14; Cocaine SA: n = 9 | Shapiro-Wilk test | Pass | Unpaired t test | Drug | / | 0.405 | 21 | 0.6896 | / |
| Extended Data Fig. 9 | c | -50 mV Saline SA: n = 14; Cocaine SA: n = 9;<br>-60 mV Saline SA: n = 14; Cocaine SA: n = 9;<br>-70 mV Saline SA: n = 14; Cocaine SA: n = 9;<br>-80 mV Saline SA: n = 14; Cocaine SA: n = 9;<br>-90 mV Saline SA: n = 14; Cocaine SA: n = 9;<br>-100 mV Saline SA: n = 14; Cocaine SA: n = 9;<br>-110 mV Saline SA: n = 14; Cocaine SA: n = 9;<br>-120 mV Saline SA: n = 14; Cocaine SA: n = 9 | Shapiro-Wilk test | Pass | Unpaired t test | Drug | / | 0.7052 | 21 | 0.4884 | / |
| Extended Data Fig. 9 | c | -50 mV Saline SA: n = 14; Cocaine SA: n = 9;<br>-60 mV Saline SA: n = 14; Cocaine SA: n = 9;<br>-70 mV Saline SA: n = 14; Cocaine SA: n = 9;<br>-80 mV Saline SA: n = 14; Cocaine SA: n = 9;<br>-90 mV Saline SA: n = 14; Cocaine SA: n = 9;<br>-100 mV Saline SA: n = 14; Cocaine SA: n = 9;<br>-110 mV Saline SA: n = 14; Cocaine SA: n = 9;<br>-120 mV Saline SA: n = 14; Cocaine SA: n = 9 | Shapiro-Wilk test | Pass | Unpaired t test | Drug | / | 1.043 | 21 | 0.3087 | / |
|  |  | -90 mV Saline SA: n = 14;<br>Cocaine SA: n = 9;<br>-100 mV Saline SA: n = 14; Cocaine SA: n = 9;<br>-110 mV Saline SA: n = 14; Cocaine SA: n = 9;<br>-120 mV Saline SA: n = 14; Cocaine SA: n = 9 |  |  |  |  |  |  |  |  |  |
| Extended Data Fig. 9 | c | -50 mV Saline SA: n = 14; Cocaine SA: n = 9;<br>-60 mV Saline SA: n = 14; Cocaine SA: n = 9;<br>-70 mV Saline SA: n = 14; Cocaine SA: n = 9;<br>-80 mV Saline SA: n = 14; Cocaine SA: n = 9;<br>-90 mV Saline SA: n = 14; Cocaine SA: n = 9;<br>-100 mV Saline SA: n = 14; Cocaine SA: n = 9;<br>-110 mV Saline SA: n = 14; Cocaine SA: n = 9;<br>-120 mV Saline SA: n = 14; Cocaine SA: n = 9 | Shapiro-Wilk test | Pass | Unpaired t test | Drug | / | / | / | / | / |
| Extended Data Fig. 9 | d | Saline SA: n = 14;<br>Cocaine SA: n = 9 | Shapiro-Wilk test | Pass | Unpaired t test | Drug | / | 3.489 | 21 | 0.0022 | / |
| Extended Data Fig. 9 | e | -90 mV Saline SA: n = 14; Cocaine SA: n = 9;<br>-100 mV Saline SA: n = 14; Cocaine SA: n = 9;<br>-110 mV Saline SA: n = 14; Cocaine SA: n = 9;<br>-120 mV Saline SA: n = 14; Cocaine SA: n = 9 | Shapiro-Wilk test | Pass | Unpaired t test | Drug | / | 0.2346 | 21 | 0.8168 | / |
| Extended Data Fig. 9 | e | -90 mV Saline SA: n = 14; Cocaine SA: n = 9;<br>-100 mV Saline SA: n = 14; Cocaine SA: n = 9;<br>-110 mV Saline SA: n = 14; Cocaine SA: n = 9;<br>-120 mV Saline SA: n = 14; Cocaine SA: n = 9 | Shapiro-Wilk test | Pass | Unpaired t test | Drug | / | 0.1896 | 21 | 0.8514 | / |
| Extended Data Fig. 9 | e | -90 mV Saline SA: n = 14; Cocaine SA: n = 9;<br>-100 mV Saline SA: n = 14; Cocaine SA: n = 9;<br>-110 mV Saline SA: n = 14; Cocaine SA: n = 9;<br>-120 mV Saline SA: n = 14; Cocaine SA: n = 9 | Shapiro-Wilk test | Pass | Unpaired t test | Drug | / | 0.6168 | 21 | 0.544 | / |
| Extended Data Fig. 9 | e | -90 mV Saline SA: n = 14; Cocaine SA: n = 9;<br>-100 mV Saline SA: n = 14; Cocaine SA: n = 9;<br>-110 mV Saline SA: n = 14; Cocaine SA: n = 9;<br>-120 mV Saline SA: n = 14; Cocaine SA: n = 9 | Shapiro-Wilk test | Pass | Unpaired t test | Drug | / | 0.4087 | 21 | 0.6869 | / |
| Extended Data Fig. 9 | g | -50 mV Saline SA: n = 14; Cocaine SA: n = 9;<br>-60 mV Saline SA: n = 14; Cocaine SA: n = 9;<br>-70 mV Saline SA: n = 14; Cocaine SA: n = 9;<br>-80 mV Saline SA: n = 14; Cocaine SA: n = 9;<br>-90 mV Saline SA: n = 14; Cocaine SA: n = 9;<br>-100 mV Saline SA: n = 14; Cocaine SA: n = 9;<br>-110 mV Saline SA: n = 14; Cocaine SA: n = 9;<br>-120 mV Saline SA: n = 14; Cocaine SA: n = 9 | Shapiro-Wilk test | Pass | Unpaired t test | Drug | / | 3.673 | 21 | 0.0014 | / |
| Extended Data Fig. 9 | g | -50 mV Saline SA: n = 14; Cocaine SA: n = 9;<br>-60 mV Saline SA: n = 14; Cocaine SA: n = 9;<br>-70 mV Saline SA: n = 14; Cocaine SA: n = 9;<br>-80 mV Saline SA: n = 14; Cocaine SA: n = 9;<br>-90 mV Saline SA: n = 14; Cocaine SA: n = 9;<br>-100 mV Saline SA: n = 14; Cocaine SA: n = 9;<br>-110 mV Saline SA: n = 14; Cocaine SA: n = 9;<br>-120 mV Saline SA: n = 14; Cocaine SA: n = 9 | Shapiro-Wilk test | Pass | Unpaired t test | Drug | / | 3.482 | 21 | 0.0022 | / |
| Extended Data Fig. 9 | g | -50 mV Saline SA: n = 14; Cocaine SA: n = 9;<br>-60 mV Saline SA: n = 14; Cocaine SA: n = 9;<br>-70 mV Saline SA: n = 14; Cocaine SA: n = 9;<br>-80 mV Saline SA: n = 14; Cocaine SA: n = 9;<br>-90 mV Saline SA: n = 14; Cocaine SA: n = 9;<br>-100 mV Saline SA: n = 14; Cocaine SA: n = 9;<br>-110 mV Saline SA: n = 14; Cocaine SA: n = 9;<br>-120 mV Saline SA: n = 14; Cocaine SA: n = 9 | Shapiro-Wilk test | Pass | Unpaired t test | Drug | / | 3.372 | 21 | 0.0029 | / |
| Extended Data Fig. 9 | g | -50 mV Saline SA: n = 14; Cocaine SA: n = 9;<br>-60 mV Saline SA: n = 14; Cocaine SA: n = 9;<br>-70 mV Saline SA: n = 14; Cocaine SA: n = 9;<br>-80 mV Saline SA: n = 14; Cocaine SA: n = 9;<br>-90 mV Saline SA: n = 14; Cocaine SA: n = 9;<br>-100 mV Saline SA: n = 14; Cocaine SA: n = 9 | Shapiro-Wilk test | Pass | Unpaired t test | Drug | / | 3.394 | 21 | 0.0027 | / |
|  |  | -110 mV Saline SA: n = 14; Cocaine SA: n = 9;<br>-120 mV Saline SA: n = 14; Cocaine SA: n = 9 |  |  |  |  |  |  |  |  |  |
| Extended Data Fig. 9 | g | -50 mV Saline SA: n = 14; Cocaine SA: n = 9;<br>-60 mV Saline SA: n = 14; Cocaine SA: n = 9;<br>-70 mV Saline SA: n = 14; Cocaine SA: n = 9;<br>-80 mV Saline SA: n = 14; Cocaine SA: n = 9;<br>-90 mV Saline SA: n = 14; Cocaine SA: n = 9;<br>-100 mV Saline SA: n = 14; Cocaine SA: n = 9;<br>-110 mV Saline SA: n = 14; Cocaine SA: n = 9;<br>-120 mV Saline SA: n = 14; Cocaine SA: n = 9 | Shapiro-Wilk test | Pass | Unpaired t test | Drug | / | 3.067 | 21 | 0.0059 | / |
| Extended Data Fig. 9 | g | -50 mV Saline SA: n = 14; Cocaine SA: n = 9;<br>-60 mV Saline SA: n = 14; Cocaine SA: n = 9;<br>-70 mV Saline SA: n = 14; Cocaine SA: n = 9;<br>-80 mV Saline SA: n = 14; Cocaine SA: n = 9;<br>-90 mV Saline SA: n = 14; Cocaine SA: n = 9;<br>-100 mV Saline SA: n = 14; Cocaine SA: n = 9;<br>-110 mV Saline SA: n = 14; Cocaine SA: n = 9;<br>-120 mV Saline SA: n = 14; Cocaine SA: n = 9 | Shapiro-Wilk test | Pass | Unpaired t test | Drug | / | 2.08 | 21 | 0.05 | / |
| Extended Data Fig. 9 | g | -50 mV Saline SA: n = 14; Cocaine SA: n = 9;<br>-60 mV Saline SA: n = 14; Cocaine SA: n = 9;<br>-70 mV Saline SA: n = 14; Cocaine SA: n = 9;<br>-80 mV Saline SA: n = 14; Cocaine SA: n = 9;<br>-90 mV Saline SA: n = 14; Cocaine SA: n = 9;<br>-100 mV Saline SA: n = 14; Cocaine SA: n = 9;<br>-110 mV Saline SA: n = 14; Cocaine SA: n = 9;<br>-120 mV Saline SA: n = 14; Cocaine SA: n = 9 | Shapiro-Wilk test | Pass | Unpaired t test | Drug | / | 0.2993 | 21 | 0.7676 | / |
| Extended Data Fig. 9 | g | -50 mV Saline SA: n = 14; Cocaine SA: n = 9;<br>-60 mV Saline SA: n = 14; Cocaine SA: n = 9;<br>-70 mV Saline SA: n = 14; Cocaine SA: n = 9;<br>-80 mV Saline SA: n = 14; Cocaine SA: n = 9;<br>-90 mV Saline SA: n = 14; Cocaine SA: n = 9;<br>-100 mV Saline SA: n = 14; Cocaine SA: n = 9;<br>-110 mV Saline SA: n = 14; Cocaine SA: n = 9;<br>-120 mV Saline SA: n = 14; Cocaine SA: n = 9 | Shapiro-Wilk test | Pass | Unpaired t test | Drug | / | 1.602 | 21 | 0.1242 | / |
|  |  | -80 mV Saline SA: n = 14;<br>Cocaine SA: n = 9;<br>-90 mV Saline SA: n = 14;<br>Cocaine SA: n = 9;<br>-100 mV Saline SA: n = 14;<br>Cocaine SA: n = 9;<br>-110 mV Saline SA: n = 14;<br>Cocaine SA: n = 9;<br>-120 mV Saline SA: n = 14;<br>Cocaine SA: n = 9 |  |  |  |  |  |  |  |  |  |
| Extended Data Fig. 9 | h | -50 mV Saline SA: n = 14;<br>Cocaine SA: n = 9;<br>-60 mV Saline SA: n = 14;<br>Cocaine SA: n = 9;<br>-70 mV Saline SA: n = 14;<br>Cocaine SA: n = 9;<br>-80 mV Saline SA: n = 14;<br>Cocaine SA: n = 9;<br>-90 mV Saline SA: n = 14;<br>Cocaine SA: n = 9;<br>-100 mV Saline SA: n = 14;<br>Cocaine SA: n = 9;<br>-110 mV Saline SA: n = 14;<br>Cocaine SA: n = 9;<br>-120 mV Saline SA: n = 14;<br>Cocaine SA: n = 9 | Shapiro-Wilk test | Pass | Unpaired t test | Drug | / | 1.455 | 21 | 0.1604 | / |
| Extended Data Fig. 9 | h | -50 mV Saline SA: n = 14;<br>Cocaine SA: n = 9;<br>-60 mV Saline SA: n = 14;<br>Cocaine SA: n = 9;<br>-70 mV Saline SA: n = 14;<br>Cocaine SA: n = 9;<br>-80 mV Saline SA: n = 14;<br>Cocaine SA: n = 9;<br>-90 mV Saline SA: n = 14;<br>Cocaine SA: n = 9;<br>-100 mV Saline SA: n = 14;<br>Cocaine SA: n = 9;<br>-110 mV Saline SA: n = 14;<br>Cocaine SA: n = 9;<br>-120 mV Saline SA: n = 14;<br>Cocaine SA: n = 9 | Shapiro-Wilk test | Pass | Unpaired t test | Drug | / | 1.4 | 21 | 0.1761 | / |
| Extended Data Fig. 9 | h | -50 mV Saline SA: n = 14;<br>Cocaine SA: n = 9;<br>-60 mV Saline SA: n = 14;<br>Cocaine SA: n = 9;<br>-70 mV Saline SA: n = 14;<br>Cocaine SA: n = 9;<br>-80 mV Saline SA: n = 14;<br>Cocaine SA: n = 9;<br>-90 mV Saline SA: n = 14;<br>Cocaine SA: n = 9;<br>-100 mV Saline SA: n = 14;<br>Cocaine SA: n = 9;<br>-110 mV Saline SA: n = 14;<br>Cocaine SA: n = 9;<br>-120 mV Saline SA: n = 14;<br>Cocaine SA: n = 9 | Shapiro-Wilk test | Pass | Unpaired t test | Drug | / | 1.201 | 21 | 0.2432 | / |
| Extended Data Fig. 9 | h | -50 mV Saline SA: n = 14; Cocaine SA: n = 9;<br>-60 mV Saline SA: n = 14; Cocaine SA: n = 9;<br>-70 mV Saline SA: n = 14; Cocaine SA: n = 9;<br>-80 mV Saline SA: n = 14; Cocaine SA: n = 9;<br>-90 mV Saline SA: n = 14; Cocaine SA: n = 9;<br>-100 mV Saline SA: n = 14; Cocaine SA: n = 9;<br>-110 mV Saline SA: n = 14; Cocaine SA: n = 9;<br>-120 mV Saline SA: n = 14; Cocaine SA: n = 9 | Shapiro-Wilk test | Pass | Unpaired t test | Drug | / | 1.579 | 21 | 0.1294 | / |
| Extended Data Fig. 9 | h | -50 mV Saline SA: n = 14; Cocaine SA: n = 9;<br>-60 mV Saline SA: n = 14; Cocaine SA: n = 9;<br>-70 mV Saline SA: n = 14; Cocaine SA: n = 9;<br>-80 mV Saline SA: n = 14; Cocaine SA: n = 9;<br>-90 mV Saline SA: n = 14; Cocaine SA: n = 9;<br>-100 mV Saline SA: n = 14; Cocaine SA: n = 9;<br>-110 mV Saline SA: n = 14; Cocaine SA: n = 9;<br>-120 mV Saline SA: n = 14; Cocaine SA: n = 9 | Shapiro-Wilk test | Pass | Unpaired t test | Drug | / | 2.085 | 21 | 0.0495 | / |
| Extended Data Fig. 9 | h | -50 mV Saline SA: n = 14; Cocaine SA: n = 9;<br>-60 mV Saline SA: n = 14; Cocaine SA: n = 9;<br>-70 mV Saline SA: n = 14; Cocaine SA: n = 9;<br>-80 mV Saline SA: n = 14; Cocaine SA: n = 9;<br>-90 mV Saline SA: n = 14; Cocaine SA: n = 9;<br>-100 mV Saline SA: n = 14; Cocaine SA: n = 9;<br>-110 mV Saline SA: n = 14; Cocaine SA: n = 9;<br>-120 mV Saline SA: n = 14; Cocaine SA: n = 9 | Shapiro-Wilk test | Pass | Unpaired t test | Drug | / | 2.626 | 21 | 0.0158 | / |
| Extended Data Fig. 9 | h | -50 mV Saline SA: n = 14; Cocaine SA: n = 9;<br>-60 mV Saline SA: n = 14; Cocaine SA: n = 9;<br>-70 mV Saline SA: n = 14; Cocaine SA: n = 9;<br>-80 mV Saline SA: n = 14; Cocaine SA: n = 9;<br>-90 mV Saline SA: n = 14; Cocaine SA: n = 9;<br>-100 mV Saline SA: n = 14; Cocaine SA: n = 9;<br>-110 mV Saline SA: n = 14; Cocaine SA: n = 9;<br>-120 mV Saline SA: n = 14; Cocaine SA: n = 9 | Shapiro-Wilk test | Pass | Unpaired t test | Drug | / | 1.996 | 21 | 0.0591 | / |
|  |  | -110 mV Saline SA: n = 14; Cocaine SA: n = 9;<br>-120 mV Saline SA: n = 14; Cocaine SA: n = 9 |  |  |  |  |  |  |  |  |  |
| Extended Data Fig. 9 | h | -50 mV Saline SA: n = 14; Cocaine SA: n = 9;<br>-60 mV Saline SA: n = 14; Cocaine SA: n = 9;<br>-70 mV Saline SA: n = 14; Cocaine SA: n = 9;<br>-80 mV Saline SA: n = 14; Cocaine SA: n = 9;<br>-90 mV Saline SA: n = 14; Cocaine SA: n = 9;<br>-100 mV Saline SA: n = 14; Cocaine SA: n = 9;<br>-110 mV Saline SA: n = 14; Cocaine SA: n = 9;<br>-120 mV Saline SA: n = 14; Cocaine SA: n = 9 | Shapiro-Wilk test | Pass | Unpaired t test | Drug | / | / | / | / | / |
| Extended Data Fig. 9 | i | Saline SA: n = 14; Cocaine SA: n = 9 | Shapiro-Wilk test | Pass | Unpaired t test | Drug | / | 3.674 | 21 | 0.0014 | / |

**Supplementary Table 16.** Cocaine training increases HCN2 and HCN4 expression in the NAc.

| Figure | Panel | Sample size | Normality test | Normality test results | Test | Comparison | *F* | *t* | df | *P* | Post hoc test |
| --- | --- | --- | --- | --- | --- | --- | --- | --- | --- | --- | --- |
| Extended Data Fig. 10 | b | Saline SA: n = 5;<br>Sucrose SA: n = 5;<br>Yoked cocaine: n = 5;<br>Cocaine SA: n = 7 | Shapiro-Wilk test | Pass | One-way ANOVA | Drug | 0.6239 | / | 3, 18 | 0.6087 | Saline SA vs. Sucrose SA: > 0.9999;<br>Saline SA vs. Yoked cocaine: > 0.9999;<br>Saline SA vs. Cocaine SA: > 0.9999 |
| Extended Data Fig. 10 | c | Saline SA: n = 5;<br>Sucrose SA: n = 5;<br>Yoked cocaine: n = 5;<br>Cocaine SA: n = 7 | Shapiro-Wilk test | Pass | One-way ANOVA | Drug | 5.73 | / | 3, 18 | 0.0062 | Saline SA vs. Sucrose SA: 0.9951;<br>Saline SA vs. Yoked cocaine: 0.0111;<br>Saline SA vs. Cocaine SA: 0.0348 |
| Extended Data Fig. 10 | d | Saline SA: n = 6;<br>Sucrose SA: n = 6;<br>Yoked cocaine: n = 6;<br>Cocaine SA: n = 8 | Shapiro-Wilk test | Pass | One-way ANOVA | Drug | 2.35 | / | 3, 22 | 0.1001 | Saline SA vs. Sucrose SA: 0.5713;<br>Saline SA vs. Yoked cocaine: 0.5517;<br>Saline SA vs. Cocaine SA: 0.0404 |

**Supplementary Table 17.** Validation of lentiviral-mediated knockdown efficiency for different HCN isoforms.

| Figure | Panel | Sample size | Normality test | Normality test results | Test | Comparison | *F* | *t* | df | *P* | Post hoc test |
| --- | --- | --- | --- | --- | --- | --- | --- | --- | --- | --- | --- |
| Extended Data Fig. 11 | a | Cocaine SA: n = 3;<br>LV-HCN1: n = 6 | Shapiro-Wilk test | Pass | Unpaired t test | Intervention | / | 7.794 | 7 | 0.0001 | / |
| Extended Data Fig. 11 | b | Cocaine SA: n = 3;<br>LV-HCN2: n = 6 | Shapiro-Wilk test | Pass | Unpaired t test | Intervention | / | 6.632 | 7 | 0.0003 | / |
| Extended Data Fig. 11 | c | Cocaine SA: n = 3;<br>LV-HCN4: n = 4 | Shapiro-Wilk test | Pass | Unpaired t test | Intervention | / | 4.378 | 5 | 0.0072 | / |

**Supplementary Table 18.** NAcC HCN blockade suppresses fentanyl seeking, normalizes high-fat CIN firing, and spares sucrose seeking.

| Figure | Panel | Sample size | Normality test | Normality test results | Test | Comparison | F | t | df | P | Post hoc test |
| --- | --- | --- | --- | --- | --- | --- | --- | --- | --- | --- | --- |
| Extended Data Fig. 12 | b | VEH: n = 11;<br>ZD7288: n = 11 | Shapiro-Wilk test | Pass | Two-way repeated-measures ANOVA | Treatment × Time | 16.37 | / | 1, 20 | 0.0006 | Last extinction (Vehicle vs. ZD7288): 0.7027;<br>Reinstatement test (Vehicle vs. ZD7288): < 0.0001 |
| Extended Data Fig. 12 | b | VEH: n = 11;<br>ZD7288: n = 11 | Shapiro-Wilk test | Pass | Two-way repeated-measures ANOVA | Treatment | 14.85 | / | 1, 20 | 0.001 | Last extinction (Vehicle vs. ZD7288): 0.7027;<br>Reinstatement test (Vehicle vs. ZD7288): < 0.0001 |
| Extended Data Fig. 12 | b | VEH: n = 11;<br>ZD7288: n = 11 | Shapiro-Wilk test | Pass | Two-way repeated-measures ANOVA | Time | 153.5 | / | 1, 20 | <0.0001 | Last extinction (Vehicle vs. ZD7288): 0.7027;<br>Reinstatement test (Vehicle vs. ZD7288): < 0.0001 |
| Extended Data Fig. 12 | c | VEH: n = 11;<br>ZD7288: n = 11 | Shapiro-Wilk test | Pass | Two-way repeated-measures ANOVA | Treatment × Time | 2.142 | / | 1, 20 | 0.1588 | Last extinction (Vehicle vs. ZD7288): 0.658;<br>Reinstatement test (Vehicle vs. ZD7288): 0.248 |
| Extended Data Fig. 12 | c | VEH: n = 11;<br>ZD7288: n = 11 | Shapiro-Wilk test | Pass | Two-way repeated-measures ANOVA | Treatment | 0.36 | / | 1, 20 | 0.5554 | Last extinction (Vehicle vs. ZD7288): 0.658;<br>Reinstatement test (Vehicle vs. ZD7288): 0.248 |
| Extended Data Fig. 12 | c | VEH: n = 11;<br>ZD7288: n = 11 | Shapiro-Wilk test | Pass | Two-way repeated-measures ANOVA | Time | 0.005 | / | 1, 20 | 0.9451 | Last extinction (Vehicle vs. ZD7288): 0.658;<br>Reinstatement test (Vehicle vs. ZD7288): 0.248 |
| Extended Data Fig. 12 | f | Normal food<br>VEH: n = 11;<br>ZD7288: n = 11<br>High-fat food<br>VEH: n = 12;<br>ZD7288: n = 11 | Shapiro-Wilk test | Pass | Two-way ANOVA | Treatment × Diet | 7.4 | / | 1, 40 | 0.0096 | Normal food (Vehicle vs. ZD7288): 0.1176;<br>High-fat food (Vehicle vs. ZD7288): <0.0001;<br>Vehicle (Normal food vs. High-fat food): <0.0001 |
| Extended Data Fig. 12 | f | Normal food<br>VEH: n = 11;<br>ZD7288: n = 11<br>High-fat food<br>VEH: n = 12;<br>ZD7288: n = 11 | Shapiro-Wilk test | Pass | Two-way ANOVA | Treatment | 37.47 | / | 1, 40 | <0.0001 | Normal food (Vehicle vs. ZD7288): 0.1176;<br>High-fat food (Vehicle vs. ZD7288): <0.0001;<br>Vehicle (Normal food vs. High-fat food): <0.0001 |
| Extended Data Fig. 12 | f | Normal food<br>VEH: n = 11;<br>ZD7288: n = 11<br>High-fat food<br>VEH: n = 12;<br>ZD7288: n = 11 | Shapiro-Wilk test | Pass | Two-way ANOVA | Diet | 26.15 | / | 1, 40 | <0.0001 | Normal food (Vehicle vs. ZD7288): 0.1176;<br>High-fat food (Vehicle vs. ZD7288): <0.0001;<br>Vehicle (Normal food vs. High-fat food): <0.0001 |
| Extended Data Fig. 12 | g | VEH: n = 7<br>ZD7288: n = 8 | Shapiro-Wilk test | Pass | Two-way repeated-measures ANOVA | Treatment × Time | 0.0052 | / | 1, 13 | 0.9438 | Last extinction (Vehicle vs. ZD7288): 0.9979;<br>Reinstatement test (Vehicle vs. ZD7288): 0.9858 |
| Extended Data Fig. 12 | g | VEH: n = 7<br>ZD7288: n = 8 | Shapiro-Wilk test | Pass | Two-way repeated-measures ANOVA | Treatment | 0.0191 | / | 1, 13 | 0.8923 | Last extinction (Vehicle vs. ZD7288): 0.9979;<br>Reinstatement test (Vehicle vs. ZD7288): 0.9858 |
| Extended Data Fig. 12 | g | VEH: n = 7<br>ZD7288: n = 8 | Shapiro-Wilk test | Pass | Two-way repeated-measures ANOVA | Time | 41.48 | / | 1, 13 | <0.0001 | Last extinction (Vehicle vs. ZD7288): 0.9979;<br>Reinstatement test (Vehicle vs. ZD7288): 0.9858 |

**Supplementary Table 19.** NAcSh HCN blockade does not affect cue-induced cocaine or sucrose seeking.

| Figure | Panel | Sample size | Normality test | Normality test results | Test | Comparison | F | t | df | P | Post hoc test |
| --- | --- | --- | --- | --- | --- | --- | --- | --- | --- | --- | --- |
| Extended Data Fig. 13 | b (left) | VEH: n = 6;<br>ZD7288: n = 8 | Shapiro-Wilk test | Pass | Two-way repeated-measures ANOVA | Treatment × Time | 1.724 | / | 4, 48 | 0.1601 | day 6: 0.3442;<br>day 7: >0.9999;<br>day 8: 0.9737;<br>day 9: 0.6003;<br>day 10: > 0.9999 |
| Extended Data Fig. 13 | b (left) | VEH: n = 6;<br>ZD7288: n = 8 | Shapiro-Wilk test | Pass | Two-way repeated-measures ANOVA | Time | 1.404 | / | 4, 48 | 0.2470 | day 6: 0.3442;<br>day 7: >0.9999;<br>day 8: 0.9737;<br>day 9: 0.6003;<br>day 10: > 0.9999 |
| Extended Data Fig. 13 | b (left) | VEH: n = 6;<br>ZD7288: n = 8 | Shapiro-Wilk test | Pass | Two-way repeated-measures ANOVA | Treatment | 0.5203 | / | 1, 12 | 0.4845 | day 6: 0.3442;<br>day 7: >0.9999;<br>day 8: 0.9737;<br>day 9: 0.6003;<br>day 10: > 0.9999 |
| Extended Data Fig. 13 | b (right) | VEH: n = 6;<br>ZD7288: n = 8 | Shapiro-Wilk test | Pass | Two-way repeated-measures ANOVA | Treatment × Time | 1.743 | / | 4, 48 | 0.1560 | day 6: 0.9998;<br>day 7: 0.2226;<br>day 8: 0.9945;<br>day 9: >0.9999;<br>day 10: 0.9995 |
| Extended Data Fig. 13 | b (right) | VEH: n = 6;<br>ZD7288: n = 8 | Shapiro-Wilk test | Pass | Two-way repeated-measures ANOVA | Time | 11.37 | / | 4, 48 | <0.0001 | day 6: 0.9998;<br>day 7: 0.2226;<br>day 8: 0.9945;<br>day 9: >0.9999;<br>day 10: 0.9995 |
| Extended Data Fig. 13 | b (right) | VEH: n = 6;<br>ZD7288: n = 8 | Shapiro-Wilk test | Pass | Two-way repeated-measures ANOVA | Treatment | 0.4332 | / | 1, 12 | 0.5229 | day 6: 0.9998;<br>day 7: 0.2226;<br>day 8: 0.9945;<br>day 9: >0.9999;<br>day 10: 0.9995 |
| Extended Data Fig. 13 | c | VEH: n = 6;<br>ZD7288: n = 8 | Shapiro-Wilk test | Pass | Two-way ANOVA | Treatment × Nose poke type | 1.386 | / | 1, 12 | 0.2619 | Active nose poke (Vehicle vs. ZD7288): 0.0915;<br>Inactive nose poke (Vehicle vs. ZD7288): 0.8163 |
| Extended Data Fig. 13 | c | VEH: n = 6;<br>ZD7288: n = 8 | Shapiro-Wilk test | Pass | Two-way ANOVA | Treatment | 3.06 | / | 1, 12 | 0.1057 | Active nose poke (Vehicle vs. ZD7288): 0.0915;<br>Inactive nose poke (Vehicle vs. ZD7288): 0.8163 |
| Extended Data Fig. 13 | c | VEH: n = 6;<br>ZD7288: n = 8 | Shapiro-Wilk test | Pass | Two-way ANOVA | Nose poke type | 12.35 | / | 1, 12 | 0.0043 | Active nose poke (Vehicle vs. ZD7288): 0.0915;<br>Inactive nose poke (Vehicle vs. ZD7288): 0.8163 |
| Extended Data Fig. 13 | e (left) | VEH: n = 7;<br>ZD7288: n = 8 | Shapiro-Wilk test | Pass | Two-way repeated-measures ANOVA | Treatment × Time | 0.4810 | / | 4, 52 | 0.7495 | day 6: >0.9999;<br>day 7: 0.9867;<br>day 8: >0.9999;<br>day 9: 0.9986;<br>day 10: 0.7937 |
| Extended Data Fig. 13 | e (left) | VEH: n = 7;<br>ZD7288: n = 8 | Shapiro-Wilk test | Pass | Two-way repeated-measures ANOVA | Time | 20.97 | / | 4, 52 | <0.0001 | day 6: >0.9999;<br>day 7: 0.9867;<br>day 8: >0.9999;<br>day 9: 0.9986;<br>day 10: 0.7937 |
| Extended Data Fig. 13 | e (left) | VEH: n = 7;<br>ZD7288: n = 8 | Shapiro-Wilk test | Pass | Two-way repeated-measures ANOVA | Treatment | 0.2219 | / | 1, 13 | 0.6454 | day 6: >0.9999;<br>day 7: 0.9867;<br>day 8: >0.9999;<br>day 9: 0.9986;<br>day 10: 0.7937 |
| Extended Data Fig. 13 | e (right) | VEH: n = 7;<br>ZD7288: n = 8 | Shapiro-Wilk test | Pass | Two-way repeated-measures ANOVA | Treatment × Time | 0.4238 | / | 3, 39 | 0.7370 | day 7: 0.5520;<br>day 8: 0.9666;<br>day 9: 0.9880;<br>day 10: >0.9999 |
| Extended Data Fig. 13 | e (right) | VEH: n = 7;<br>ZD7288: n = 8 | Shapiro-Wilk test | Pass | Two-way repeated-measures ANOVA | Time | 25.60 | / | 3, 39 | <0.0001 | day 7: 0.5520;<br>day 8: 0.9666;<br>day 9: 0.9880;<br>day 10: >0.9999 |
| Extended Data Fig. 13 | e (right) | VEH: n = 7;<br>ZD7288: n = 8 | Shapiro-Wilk test | Pass | Two-way repeated-measures ANOVA | Treatment | 0.8120 | / | 1, 13 | 0.3839 | day 7: 0.5520;<br>day 8: 0.9666;<br>day 9: 0.9880;<br>day 10: >0.9999 |
| Extended Data Fig. 13 | f | VEH: n = 7;<br>ZD7288: n = 8 | Shapiro-Wilk test | Pass | Two-way ANOVA | Treatment × Nose poke type | 0.16 | / | 1, 13 | 0.6957 | Active nose poke (Vehicle vs. ZD7288): 0.9826;<br>Inactive nose poke (Vehicle vs. ZD7288): 0.7950 |
| Extended Data Fig. 13 | f | VEH: n = 7;<br>ZD7288: n = 8 | Shapiro-Wilk test | Pass | Two-way ANOVA | Treatment | 0.2177 | / | 1, 13 | 0.6485 | Active nose poke (Vehicle vs. ZD7288): 0.9826;<br>Inactive nose poke (Vehicle vs. ZD7288): 0.7950 |
| Extended Data Fig. 13 | f | VEH: n = 7;<br>ZD7288: n = 8 | Shapiro-Wilk test | Pass | Two-way ANOVA | Nose poke type | 85.06 | / | 1, 13 | <0.0001 | Active nose poke (Vehicle vs. ZD7288): 0.9826;<br>Inactive nose poke (Vehicle vs. ZD7288): 0.7950 |
